# A fuzzy sequencer for rapid DNA fragment counting and genotyping

**DOI:** 10.1101/2023.10.24.563729

**Authors:** Wenxiong Zhou, Li Kang, Shuo Qiao, Haifeng Duan, Chenghong Yin, Chao Liu, Zhizhao Liao, Mingchuan Tang, Ruiying Zhang, Lei Li, Lei Shi, Meijie Du, Yipeng Wang, Wentao Yue, Yan Xiao, Lin Di, Xiannian Zhang, Yuhong Pang, Mingkun Li, Lili Ren, Jianbin Wang, Zitian Chen, Yanyi Huang

## Abstract

High-throughput sequencing technologies generate a vast number of DNA sequence reads simultaneously, which are subsequently analyzed using the information contained within these fragmented reads. The assessment of sequencing technology relies on information efficiency, which measures the amount of information entropy produced per sequencing reaction cycle. In this study, we propose a fuzzy sequencing strategy that exhibits information efficiency more than twice of currently prevailing cyclic reversible terminator sequencing methods. To validate our approach, we developed a fully functional and high-throughput fuzzy sequencer. This sequencer implements a highly efficient fluorogenic sequencing-by-synthesis chemistry and underwent testing across various application scenarios, including copy-number variation detection, noninvasive prenatal testing, transcriptome profiling, mutation genotyping, and metagenomic profling. Our findings unequivocally demonstrate that the fuzzy sequencing strategy outperforms existing methods in terms of information efficiency and delivers accurate resequencing results with faster turnaround times.

**One Sentence Summary:** A fuzzy sequencer can exceed current limit of information efficiency of DNA sequencers for resequencing applications.

---

High-throughput ‘next-generation’ sequencing (NGS)^1–2^ has been extensively employed for precise quantification of nucleic acids at a whole genome, exome, or transcriptome scale^3–4^. Among the primary applications of NGS is resequencing, which encompasses copy-number variation (CNV) assessment^5–6^, transcriptome profiling (RNA-seq)^7–8^, and noninvasive prenatal testing (NIPT)^9^ that rely on counting sequence-based DNA fragments. The accuracy of counting largely depends on the effectiveness of extracting information from DNA fragments and accurately mapping it onto reference sequences. Although mapping algorithms continue to evolve, the sequencing steps themselves have remained largely unchanged for several years.

Next-generation sequencing relies on the incorporation of nucleotides to encode information, and the efficiency of information encoding varies across different systems. The majority of high-throughput DNA sequencers utilize sequencing-by-synthesis (SBS) chemistries, with signal generation schemes falling into two major variations: single-nucleotide addition (SNA) or cyclic reversible terminator (CRT)^10–11^. We define the intrinsic information efficiency of a sequencer as the information entropy per cycle of the sequencing signals. According to this definition, SNA chemistry produces natural DNA duplex but suffers from relatively low intrinsic information efficiency, averaging at 0.67 nt/cycle or 1.33 bit/cycle for encoding random sequences. In contrast, multi-step CRT chemistry, although producing scar-containing duplex, offers 2 bit/cycle intrinsic information efficiency with an exact 1 nt/cycle extension rate (Supplementary Fig. 1a). Enhancing the information entropy encoding efficiency becomes a clear avenue for improving sequencing performance.

While the information entropy embedded in the nucleotide sequence plays a decisive role in mapping a read to a reference, mapping accuracy is also influenced by the corresponding region of the reference. One example is the mate-pair reads which, compared with conventional paired-end reads, are more mappable because of larger reference target size instead of higher information entropy. This effect becomes particularly evident in genomes with a high proportion of repeat sequences. From the perspective of the sequencer, we define the extrinsic information efficiency as the information entropy per cycle of the sequenced DNA fragment, or two times of the read length per cycle. This reference-side extrinsic information is equally significant but often overlooked. This oversight is primarily due to prevalent DNA sequencers providing explicit sequences of four bases, where the intrinsic information efficiency is equal to the extrinsic information efficiency, despite the differences in efficiency between SNA and CRT. Enhancing the extrinsic information efficiency could also lead to improvements in sequencing technology.

Decoupling the intrinsic and extrinsic information entropy within the same read suggests a change in sequencing read formatting. When aligning a DNA fragment sequence to reference sequences, the alignment process relies not on the string format, but rather on the information entropy contained within it. The reference genome is typically represented by the conventional and natural four base-letters (A, C, G, and T), but the read itself does not necessarily need to adhere to the same formatting. As long as the information is sufficiently rich for unique alignment, alternative formatting can be employed. Although sequencing by oligonucleotide ligation and detection (SOLiD) is being phased out in the market due to low-efficiency chemistry and very short read lengths, it demonstrated the possibility of sequencing with alternate formatting through a color-space strategy^12^.

From a sequencing strategy standpoint, achieving higher information efficiency (> 2 bit/cycle) for both intrinsic and extrinsic information entropy could revolutionize the entire sequencing field. In this report, we introduce a fuzzy SBS strategy that sacrifices explicit sequencing information in exchange for improved information encoding performance. The fuzzy SBS sequencing strategy is based on the fluorogenic sequencing chemistry using the terminal phosphate-labeled fluorogenic nucleotides^13–15^ (TPLFN, Fig. 1a) we reported earlier. In this chemistry, the fluorophore Tokyo Green (TG) labeled TPLFN can turn from a dark state to a light state after catalysis by the Bst polymerase and alkaline phosphatase, and its fluorescence intensity indicates the number of the incorporated nucleotides (Fig. 1b). We constructed DNA fragments into a library with two specific end adapters for clonal amplification on hydrogel microbeads carrying amplification primers. The DNA fragments hybridized on the beads were then amplified into clones through emulsion PCR^16^ or surface-tethered recombinase polymerase amplification (RPA) (Fig. 1c). Each clone typically contained approximately 30,000 copies of DNA for the SBS reaction. After annealing with the sequencing primer, the DNA fragments can be effective sequenced through a conventional SNA strategy (Fig. 1d).

**Fig. 1.**
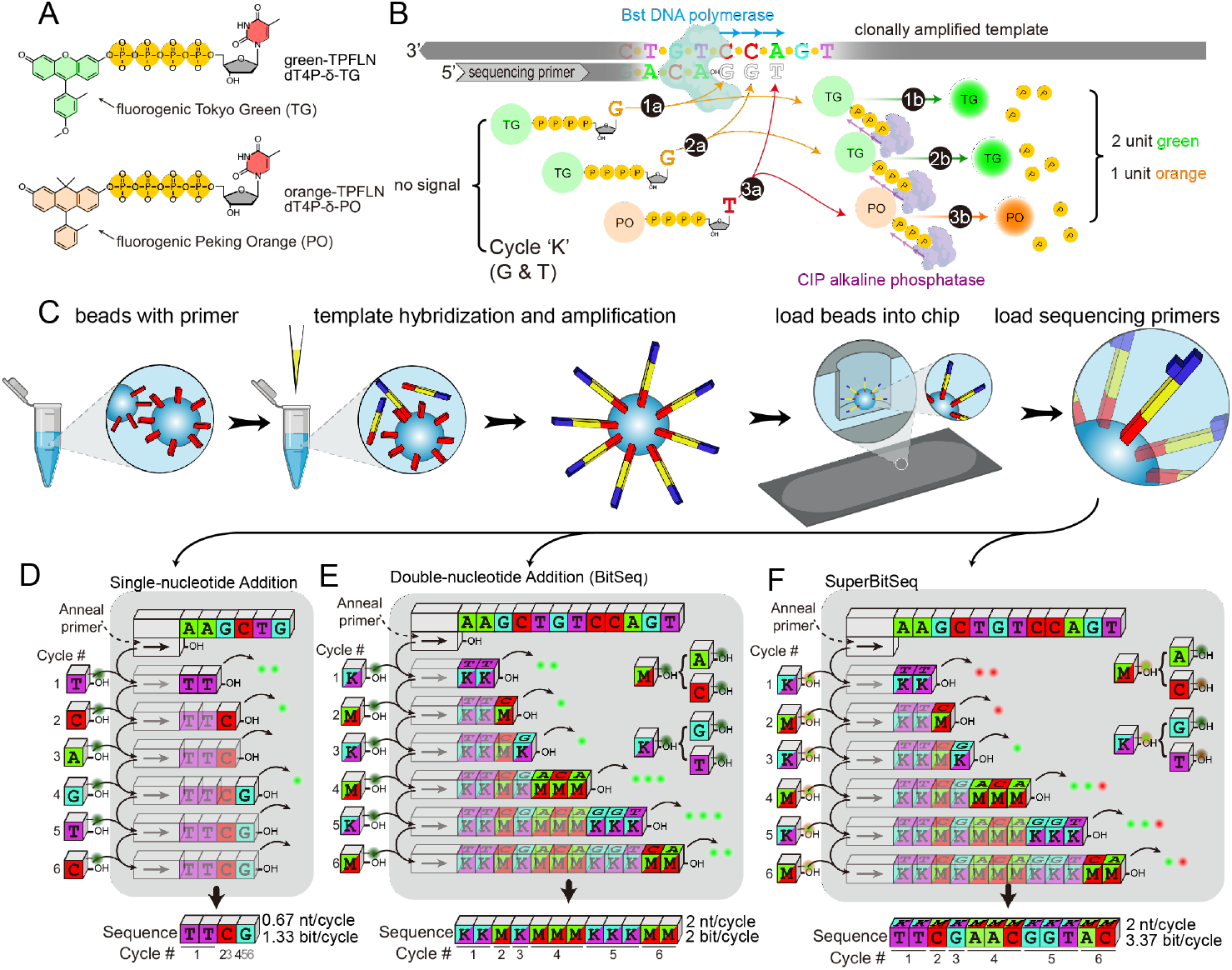
Schematic of fuzzy sequencing. (A) Chemical structure of the terminal phosphate-labeled fluorogenic nucleotides (TPLFN) used in fuzzy sequencing. (B) Schematic of the fluorogenic sequencing chemistry adopted by fuzzy sequencing. (C) Schematic of the fuzzy sequencing pipeline. The DNA templates are amplified on the beads and loaded into the wells of the sequencing chip. After hybridized by the sequencing primers, the DNA templates are primed for sequencing. (D-F) Schematic of the nucleotide flowgrams used by single-nucleotide addition (D), monochromatic double-nucleotide addition (BitSeq, E) and dichromatic double-nucleotide addition (SuperBitSeq, F).

Fuzzy SBS can be implemented using different flowgrams (Fig. 1e-f, Supplementary Fig. 1b), allowing the introduction of nucleotides beyond the limitation of a single type per cycle (see Supplementary Text 2.1). Briefly, *BitSeq* employs a double-nucleotide addition flowgram, where a combination of two nucleotides is alternately added in each cycle. For instance, K (a mix of G/T) is added during odd cycles, while M (a mix of A/C) is added during even cycles (see Fig. 1e). BitSeq generates fuzzy sequences since it’s not possible to distinguish the signal associated with the incorporation of the two nucleotides in the mix. Similarly, TritSeq employs a triple-nucleotide addition flowgram, where a combination of three nucleotides is alternately added in each cycle (Supplementary Fig. 1b). All of these fuzzy SBS approaches can achieve intrinsic and/or extrinsic information efficiency beyond 2 bits per cycle, effectively reducing the cycle number required to achieve a specific read length and ultimately leading to faster turn-around speed (Supplementary Table 1). In this paper, the focus will primarily be on BitSeq and its dichromatic form SuperBitSeq as they strike a reasonable balance between information efficiency and signal dynamic range per reaction cycle.

We constructed a laboratory prototype sequencer for conducting high-throughput BitSeq (Fig. 2a,b). The bit sequencer comprises a microfluidic flow cell for implanting clonally amplified DNA fragments for sequencing and an imaging system for signal acquisition (Supplementary Fig. 2-4). We implemented fluorogenic SBS chemistry to naturally align with the BitSeq flowgram^13–15,17^.

**Fig. 2.**
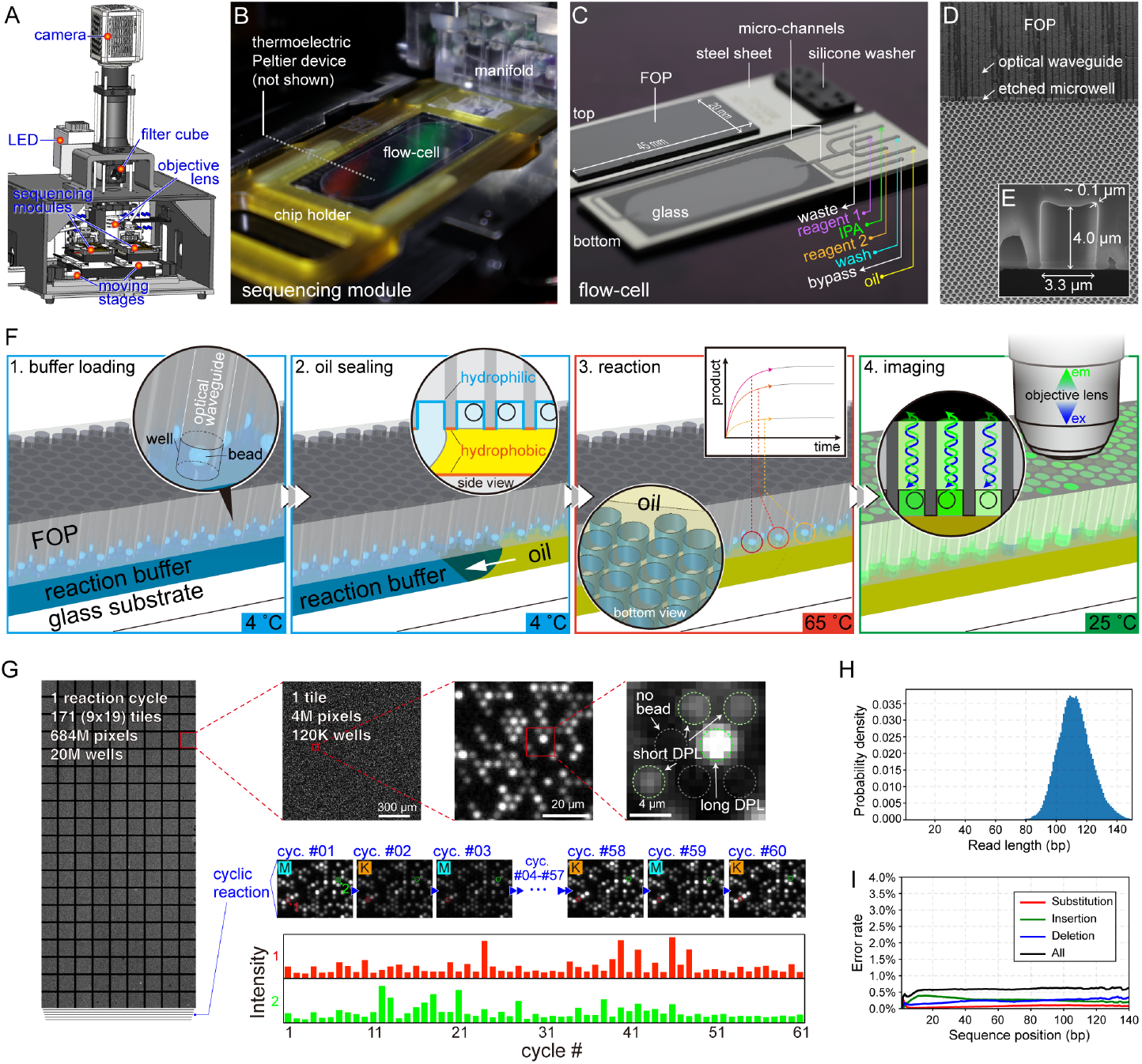
Implementation of BitSeq. (A) The design of a prototype for high-throughput fluorogenic fuzzy sequencing. (B) The photo of a sequencing module, which consists of a flow-cell connected with a manifold for liquid routing, and the flow-cell is placed on a thermoelectric Peltier device for temperature control. (C) The photo of a sequencing flow-cell, which is made of a glass slide and a fiber optical plate (FOP) and a flow channel in between with shape defined by laser-cut double side adhesive. (D) Scanning electron microscopy image of the microwell array made of selective etching of FOP. These femtoliter microwells are placed inward the flow chamber. (E) The size of a microwell. (F) The running procedure of one sequencing cycle. Firstly, the micro-beads with clonal amplified templates for sequencing are introduced into the microwells and tethered on the inner surface of the microwells. Then, the reaction buffer is primed into the flow chamber and all microwells are sealed by oil flowing into the chamber to separate each microwell from cross-talk. The SBS reaction is triggered by elevation of the temperature. The fluorogenic product is proportional to the bases that incorporated during each reaction cycle. When the reaction is finished, the flow-cell is cooled to take fluorescent images. (G) One typical field of view from 171 tiles a cycle and the procedure of fluorescence intensity extraction. The microwells that contained beads will generate fluorescence after reaction of each cycle, and the higher intensity indicates longer degenerate polymer length (DPL) in that reaction cycle. Each microwell is addressable and indexed to produce an intensity series that can be later deduced into bit sequences. (H) The read length distribution of bit sequences. (I) The error rate of BitSeq.

The microfluidic flow cell (Fig. 2c) contains 28 million 30-femtoliter microwells (Fig. 2d,e). Each cycle of the SBS reaction can extend one or more bases and produce a corresponding amount of fluorophore. The microwells can be sealed to prevent signal cross-talk due to fluorophore diffusion. These microwells were fabricated by selective wet-etching of fiber-optic plates (FOP), allowing the fluorogenic signal to propagate through the fiber waveguide to the outer surface of the flow cell for imaging (Fig. 2f). The inner surface of the microwells was made hydrophilic, while the adjoining surface between microwells was topologically coated to be hydrophobic, enabling the highly parallel sealing of the SBS reactions using oil (Supplementary Fig. 5,6).

The oil-based microwell sealing is robust, reversible, and compatible with fluorogenic SBS reaction conditions. In each cycle of the fluorogenic SBS reaction, we first filled the microwells with reaction buffer containing polymerase and two fluorogenic unnatural nucleotides at low temperature of 4 °C. We then sealed the microwells by steadily flowing in fluorinated oil (Fig. 2f). Next, we quickly raised the temperature of the flowcell to 65°C to initiate the synthesis reaction, followed by cooling down to 25°C for fluorescence image acquisition. After that, we removed the sealing by flowing in isopropanol and performed an aqueous wash to reset the reaction conditions for the next cycle. The acquired images were aligned to identify each microwell, which was then registered for tracking throughout all reaction cycles (Fig. 2g).

Defected regions resulting from local sealing or unsealing failures were eliminated before further processing (Supplementary Fig. 7A). Empty microwells and microwells with multiple beads were identified and filtered out of further analysis.

The fluorescence signal of each monoclonal microwell was extracted from the images after noise reduction and background correction. Due to the asynchronization of the molecular reaction within each clone, the intensity-cycle series could not be directly converted into bit sequences without a meticulous dephasing process. To supervise the dephasing algorithm, we spiked a standard DNA library, constructed from the lambda phage genome, into the samples to be sequenced together. As a result, the sequenced reads could be easily classified into two categories: standard dots (SD) and library dots (LD), based on the characteristic signal profiles of the first few cycles.

We employed the dual-base flowgram dephasing model to fit the SD signals and obtain the parameters^18^, which were then applied to LD signals to construct a flux matrix for dephasing correction and to convert the signals into bit sequences.

We initially tested our prototype sequencer on a lambda phage genomic library (Supplementary Fig. 8, Supplementary Table 2). After 61 cycles of repeated sealing and sequencing, more than 90% of the chip area remained unaffected by stains (Supplementary Fig. 7B, 11). In a single run, we generated 6,578,364 reads with an average length of 112.2 bp (Fig. 2h, Supplementary Fig. 9-15). To map the BitSeq signals, we encode them as sequence reads that can be aligned using conventional mappers such as Burrows-Wheeler Aligner (BWA)^19–20^ and Bowtie2 (Supplementary Text 2.2, Supplementary Table 3-6)^21–22^. By mapping the bit sequences back to the reference genome, we successfully aligned 99.11% of the reads, which had an average length of 111.6 bp. The error rate of BitSeq was found to be 0.39% in the first 10 bp and 0.62% in the 130-140 bp region, which were acceptably low. This indicates the feasibility of accurate read mapping and DNA fragment counting (Fig. 2i).

Using the same aligning strategy, we then evaluated the performance of the BitSeq approach in four widely used resequencing applications: copy number variation (CNV) identification, noninvasive prenatal testing (NIPT), transcriptomic analysis (RNA-seq), and metagenomics sequencing (mNGS).

We sequenced two genomic DNA samples collected from a normal male and a Down syndrome male patient at low coverage (0.1X). The sequencing coverage across the whole genome was mostly uniform for both samples (Supplementary Fig. 16,17) except for the sex chromosomes which were half of that of autosomes. The additional copy of chromosome 21 in the Down syndrome patient was accurately counted through BitSeq, as validated by Ion Torrent sequencer as well.

Besides aneuploidies, BitSeq can also faithfully detect the clinically relevant small size CNVs. We used BitSeq to identify the 2.9-Mb deletion at Chr22.q11.21 that was associated with DiGeorge syndrome from a patient, and two CNVs (20.8-Mb gain at Chr 11.q and 10.0-Mb gain at Chr 22.q11) from another patient with development delay (Fig. 3a) with high concordance with other conventional sequencing approaches. Notably, BitSeq showed lower median of the absolute values of all pairwise differences (MAPD) than CRT when truncated to read lengths with the same sequencing cycles, indicating more precise CNV determination (Fig. 3b).

**Fig. 3.**
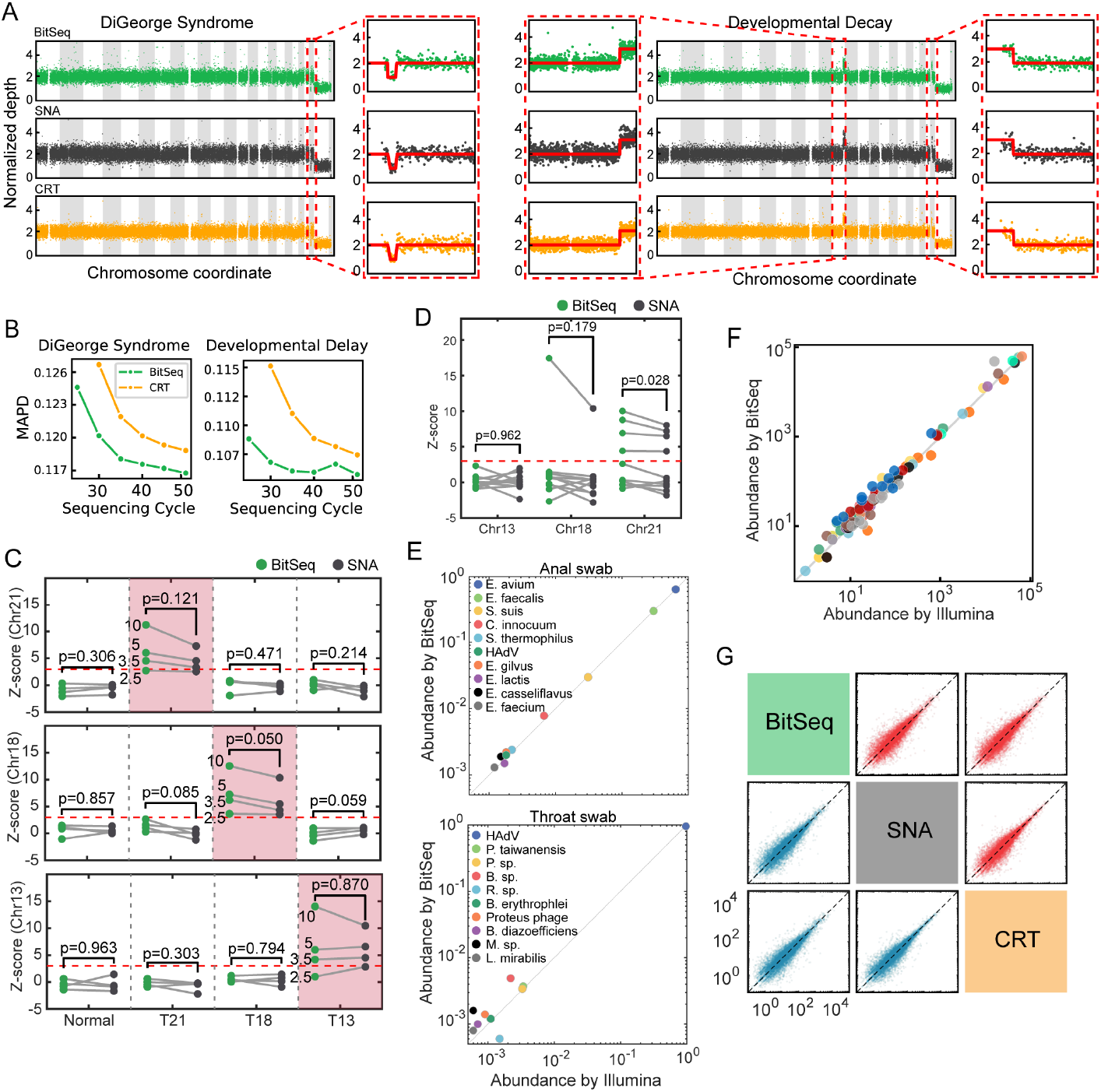
Comparison of BitSeq with commercial sequencers in resequencing. (A) CNV of a DiGeorge syndrome patient (left) and a development-delayed patient (right). (B) BitSeq shows lower MAPD than CRT under the same sequencing cycles. MAPD for CRT under 25 cycles is missing because such short reads cannot be mapped uniquely to the genome. (C) Z-scores of normal and mock trisomy NIPT ccfDNA samples. (D) Z-scores of single-blinded NIPT ccfDNA samples. The red dash in B-C indicates the threshold of 3. (E-F) Microbe abundance comparison between BitSeq and Illumina by metagenomic sequencing (E) and targeted metagenomic sequencing (F). (G) TPM comparison of MEF (blue) and mES (red) cell line between BitSeq and commercial sequencers.

We then tested the effectiveness of BitSeq on NIPT, a widely used resequencing application in prenatal diagnostics that requires high accuracy of DNA counting. We started the test with mock samples prepared by spiking fragmented genomic DNA from trisomy patients into circulating cell-free DNA (ccfDNA) of an unpregnant woman with different ratios. These 12 mock NIPT samples had trisomy in Chr21, Chr18 and Chr13, respectively, and the mixing proportion was 10%, 5%, 3.5% and 2.5%. We also collected 28 NIPT ccfDNA samples from pregnant women with normal fetuses. We applied BitSeq to all the samples and used 24 out of the 28 normal samples as the control set (the other 4 used as the negative test set) to calculate the Z-score for each NIPT sample (Fig. 3c). Using Z=3 as the cut-off, BitSeq shows comparable power to identify the positive trisomy NIPT samples when the mock fetal DNA fraction is not less than 3.5%. We further tested 10 single-blinded true NIPT samples, 5 of which were tested positive based on the same cut-off using BitSeq (Fig. 3d). These results were confirmed by parallel tests by Ion Torrent sequencers as well as the clinical records (Supplementary Fig. 18,19a).

Combined with metagenomic sequencing, BitSeq may identify pathogens in acute infections timely. We tested the capability of BitSeq in metagenomics by sequencing one anal swab and one throat swab from a previously reported pediatric patient suffered from multi-organ abscesses^23^. The 10 species types with the highest abundance identified by BitSeq and Illumina sequncer are identical. And their abundances, ranging from less than 0.1% to over 95%, are also close (Fig. 3e). In addition, using a custom panel, we also tested BitSeq on targeted metagenomics by sequencing bronchoalveolar lavage fluid samples from 14 patients with community-acquired pneumonia. The species types as well as their normalized read number reflecting their relative abundance, are also consistent (Fig. 3f).

We next verified that BitSeq could be used for RNA-seq. Using mouse embryonic fibroblasts (MEF) and embryonic stem (mES) cells, BitSeq could provide almost identical results as Ion Torrent or Illumina sequencers did. The number of genes detected was comparable, and the gene expression levels were correlated with high linearity between these approaches (Fig. 3g, Supplementary Fig. 19b-c). We also checked the Gene Ontology (GO) terms of genes only detected by one sequencer and found no specific preferences in between (Supplementary Fig. 19d-f).

All these experiments show that BitSeq is fully capable for those applications based on DNA fragment counting. We expect that other re-sequencing approaches that do not require single-base resolution but focus on pattern alterations such as bisulfite sequencing for DNA methylation^24–26^ or Hi-C for chromatin topology analysis^27–28^ are also possible with BitSeq. Thanks to its high information and long extension length per cycle as well as the fast fluorogenic sequencing chemistry, BitSeq would enable more efficient DNA fragment counting that is valuable in both basic science research and clinical testing.

We further considered the possibility to extend fuzzy sequencing strategy from BitSeq to SuperBitSeq by labeling the two different nucleotides in each reaction cycle with distinguishable fluorophores (Fig. 1f). SuperBitSeq possesses the same extrinsic information efficiency as BitSeq, but even higher intrinsic information efficiency (Supplementary Fig. 1d).

Like BitSeq, we also designed an encoding strategy for SuperBitSeq that is compatible with prevailing short reads mappers like BWA (Fig. 4a). We conducted *in silico* analysis to validate the alignment accuracy of the fuzzy SBS encoding strategy. For our analysis, we randomly selected 1×10^7^ positions from the human reference genome and 1×10^6^ positions from the *Arabidopsis* reference genome. We simulated the BitSeq, SuperBitSeq and CRT sequencing from these positions by extending the nucleotide strings with certain cycles (Supplementary Fig. 20), and then aligned the simulated reads back to their respective reference genomes. We observe that reads generated with more sequencing cycles provided greater information, resulting in a higher unique mapping rate (UMR) for both fuzzy sequencing and CRT approaches. Notably, SuperBitSeq consistently exhibited the highest UMR (Fig. 4b, Supplementary Fig. 21, Supplementary Table 7-58). Additionally, the UMR of BitSeq with the M-K form was consistently higher than that of CRT and BitSeq with R-Y form. This preference for the M-K form may be attributed to the occurrence of transition base substitutions follow gene duplication during evolution^29^.

**Fig. 4.**
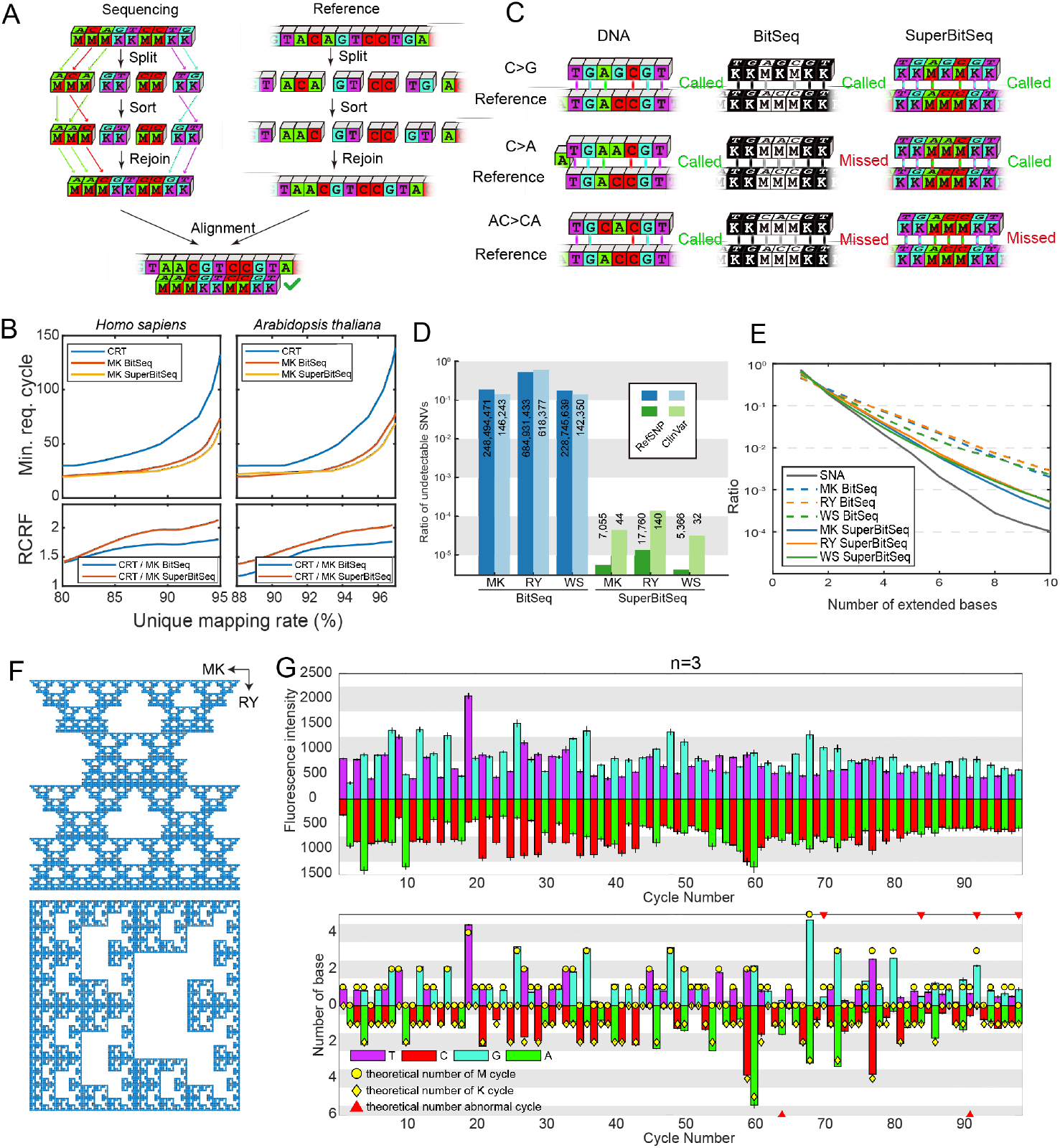
Properties of SuperBitSeq. (A) Encoding strategy of SuperBitSeq in M-K form as an example.(B) Simulated unique mapping rate of DNA reads by fuzzy sequencing and CRT for different genomes. For mapping the specific proportion of the genome, both BitSeq and SuperBitSeq need much shorter minimally required sequencing cycles, and this advantage can also be reflected by reaction cycle reduction factor (RCRF) that defined by the ratio of minimally required cycles between technologies.(C) Distinguishable and indistinguishable SNV types of BitSeq and SuperBitSeq. (D) Ratio of indistinguishable SNV by BitSeq and SuperBitSeq. (E) Freqencies of extended base number of different flowgrams. (F) Fractal of encoded SuperBitSeq signals. (G) The fluorescence intensities (top) and their dephasing corrected signals (bottom) of a single-template SuperBitSeq experiment.

The elevated information efficiency enables the detection of significantly more SNVs that cannot be identified through BitSeq (Fig. 4c). For example, in BitSeq the A>C/G>T mutations cannot be identified in the M-K form, nor the A>G/C>T mutations in the R-Y form. In SuperBitSeq they can all be well captured, except for only specific indistinguishable base swap events such as the AC>CA in the M-K form. To evaluate the potential of SuperBitSeq for SNP/SNV detection, we filtered dbSNP (build 155)^30^ and ClinVar (20210908)^31^ to exclude items with N, and obtained 1,297,977,577 and 1,009,591 items respectively. About 17-52% of known human SNP/SNVs cannot be distinguished by BitSeq, whereas the undetection rates are ∼0.01% or even lower for SuperBitSeq (Fig. 4d). This means that SuperBitSeq can *de facto* detect almost all known SNP/SNVs. More importantly, in each reaction cycle the fluorescence signal released by extended bases can be split into two channels, resulting in reduced requirement for signal detection dynamic range and enhanced accuracy in detecting long copolymers (Fig. 4e, Supplementary Fig. 22). Interestingly, we also found that the encoded superbit sequences can form a fractal when mapped the points in a square (Fig. 4f, Supplementary Text 2.4).

To implement SuperBitSeq experimentally, we synthesized a novel fluorophore, Peking Orange (PO), which was designed to exhibit excellent fluorogenic properties (on/off ratio ∼ 2000) when terminally labeled on the tetraphosphate nucleotides (PO-TPLFNs, or PO-dN4P, Fig. 1a, Supplementary Fig. 23,24,25a). PO was used to pair with TG for labeling two different nucleotides, respectively, in each reaction cycle. PO and TG have similarly high fluorescence quantum efficiency but distinct excitation and emission spectra to avoid signal cross-talk (Supplementary Fig. 25b). PO-dN4P has great photo and thermal stability, and showed excellent signal-to-base linearity upon nucleotide extension reaction (Supplementary Fig. 25c-e). The same dephasing algorithm we optimized for TG-dN4P substrates could be seamlessly adopted to PO-dN4P when sequencing a single template (Supplementary Fig. 26).

According to our previous *in silico* simulation, dephasing correction for SuperBitSeq could be built upon the algorithm of BitSeq by separating two channels for independent correction with the same dephasing parameters^18^. We also experimentally confirmed this strategy using the same single template for SuperBitSeq (Fig. 4g, Supplementary Fig. 27).

We added a fluorescence channel to the bit sequencer and converted it to the superbit sequencer (Fig. 5a, Supplementary Fig. 28), while the image processing and signal extraction pipeline was largely adapted from before. We demonstrated the SNV detectability of SuperBitSeq by sequencing the G719S and T790M mutations of EGFR gene (Fig. 5b). The signal acquired from 9 imaging tiles (0.67 × 0.67 mm^2^ each, 212,874 reads in total) could be clearly clustered into 4 groups (Fig. 5c,d), and each group represented one of the DNA templates with single nucleotide difference. With dephasing correction, the signals of these clusters clearly unveiled the significant difference at the specific reaction cycle, which is associated with the SNVs we targeted.

**Fig. 5.**
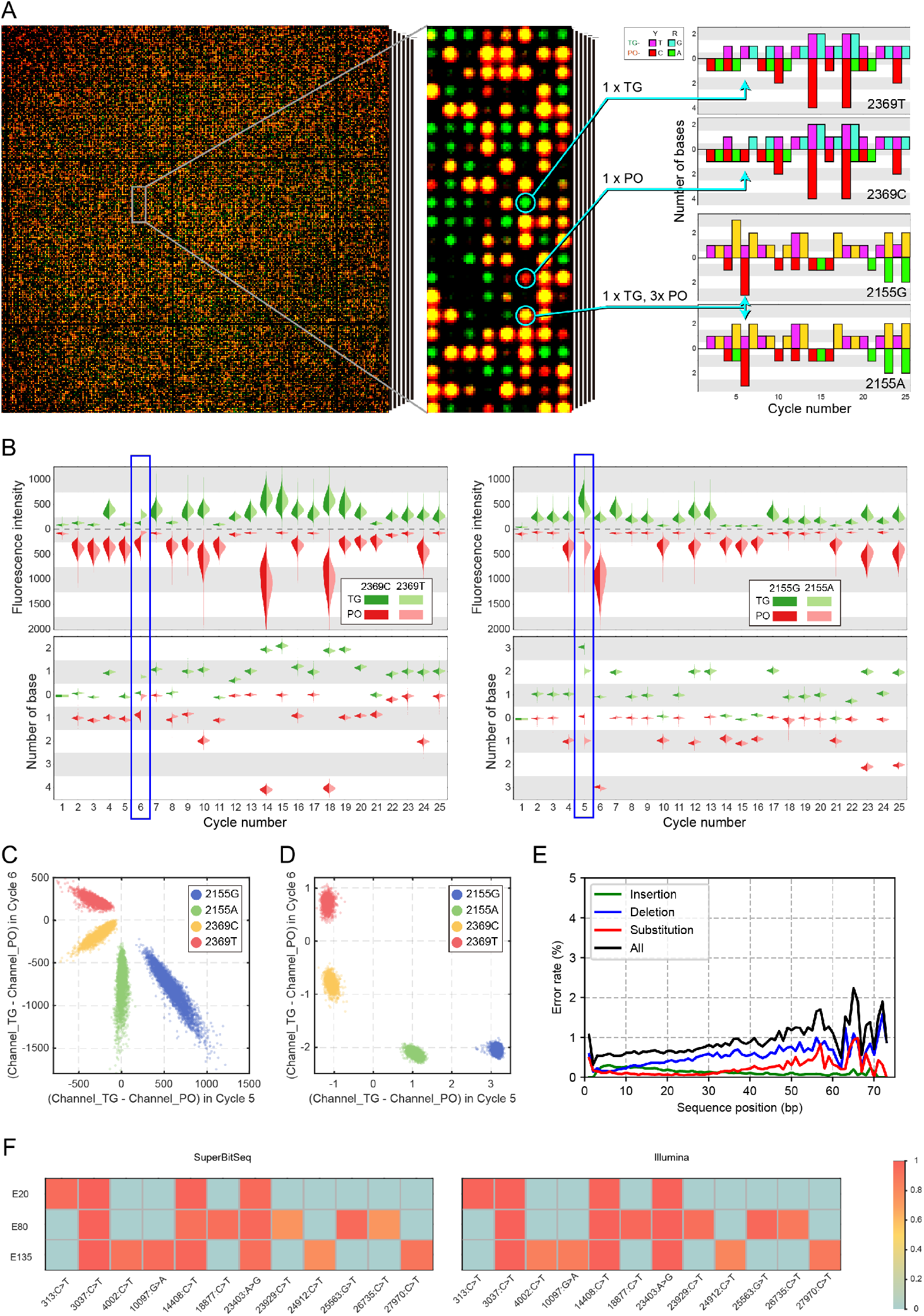
High-throughput SuperBitSeq identifies single-nucleotide variants (SNVs). (A) A merged fluorescence image of green (TG-labeled T or G) and red (PO labeled C or A) channels, from which the signal intensities are extracted. The image was associated with Cycle 6 in this specific experiment and yellow dots represent the co-existing of green and red-signal. The signal series of four different DNA templates are shown in the right. (B) Violin plot of fluorescence intensity and dephasing corrected signals of four different DNA templates. The blue boxes show the distinguishable signals that differentiate the SNVs using SuperBitSeq. (C-D) The four different DNA templates can be distinguished by the difference of fluorescence intensities (C) or dephasing-corrected signals (D) in Cycle 5-6. (E) Error rate of SuperBitSeq in SARS-CoV-2 samples. (F) Allele frequencies in SARS-CoV-2 samples detected by SuperBitSeq and an Illumina sequencer.

While BitSeq can only determine the pathogen species and abundances in metagenomic sequencing, SuperBitSeq allows fast pathogen genotyping and origin tracing, which is valuable in epidemic outbreaks. We sequenced 3 SARS-CoV-2 samples from a previously reported epidemiology study^32^ for 26 cycles using SuperBitSeq. The total reaction time is only 42.6 min and we got about 200k reads. After mapping the SuperBitSeq reads to the reference genome, we restored the SNV information by separating the read and the reference into two semi-sequences, aligning individually, and then merging together (Supplementary Fig. 29). The experimental data demonstrated low error rate (≈1%, Fig. 5e) and an average read length of 43 bp (Supplementary Fig. 30a). The SNVs called by SuperBitSeq are consistent with those called by an Illumina sequencer (Fig. 5f).

One reasonable extension of our information-richer SuperBitSeq is to combine three rounds of SuperBitSeq with orthogonal dual-base mixes for error correction code (ECC) sequencing, which may surpass the previously demonstrated high accuracy ECC. Our *in silico* simulation showed that compared to traditional ECC, error-correction using three rounds of superbit sequences has lower false negative rate in detecting potential sequencing errors under the same noise level (Supplementary Fig. 31). Both BitSeq and SuperBitSeq are specific forms of the fuzzy SBS strategy that encode DNA with high information efficiency for sequencing. More advanced forms such as TritSeq or SuperTritSeq are definitely worth of more expectation.

However, the signal dynamic range of TritSeq would be too large to practice for fluorogenic chemistry, despite TritSeq would generate longer extension length and higher information efficiency than BitSeq. There are three major challenges that have not been well addressed in this work. First, the fundamental mathematical structure of the fuzzy coding of DNA is yet fully understood. For example, after transforming the bit sequences into binary faction number and then converting into decimal fraction numbers, every infinite long DNA sequence can be mapped to a point in the square region [0,1) × [0,1) (Fig. 4f), and formed fractal patterns for both SuperBitSeq with M-K and W-S forms. These two fractal patterns have identical Hausdorff dimension of ≈ 1.7716, although the nature of it is still unknown (Supplementary Text 2.4).

Second, currently available fragment mapping algorithms and softwares, such as BWA or Bowtie2, were designed for recognizing natural base format but not the fuzzy sequences. Although we have circumvented this problem by renaming the bit strings to be compatible with current softwares, a more general mapping tool that directly handles the nature of fuzzy sequence is still highly desirable. Last, further optimizations are still required to reduce the overall operational time and cost.

Although cycle number reduction by fuzzy sequencing leads to reduced reaction time and reagent cost, multiple steps such as template amplification, fluidics and imaging still take up a large proportion of time, and the throughput should be further increased to reduce the average cost.

## Supporting information

The Supplementary Information

## Acknowledgments

The authors thank Ying Shang, Wentao Li, Lingguo Du, Xiaomeng Wu, Bingxiao Feng, Jiawei Huang and Runxin Du for experimental and analytical assistance. **Funding**: Funding was partially provided by National Natural Science Foundation of China Grant 21927802 (Y.H., J.W.), T2225005 (J.W.) and T2188102 (Y.H.), Beijing Municipal Science and Technology Commission Grant Z201100005320016 (Y.H.), Z211100003321006 (Y.H.) and Z221100007022003 (Y.H.), and Beijing Advanced Innovation Center for Genomics.

## Author contributions

Conceptualization: Y.H., Z.C., and W.Z. Experiment: W.Z., L.K., S.Q., H.D., C.L., R.Z., L.S., Y.X., W.Y., Y.W., L.D., Y.P., M.L., L.R., and C.Y. Data Analysis: W.Z., L.K., S.Q., Z.L., M.T., L.L., M.D., X.Z., J.W., Z.C., and Y.H. Writing: W.Z., L.K., J.W., Z.C., and Y.H.

## Competing interests

W.Z., L,K., S.Q., H.D., Z.C. and Y.H. are inventors on the patent applications of fuzzy sequencing technologies. H.D., Z.C. and Y.H. are founders, and R.Z., L.L. and L.S. are employees, of Cygnus Biosciences. Other authors declare no competing interests.

## Notes

### Summary of Updates

Figure 1-4 revised. Supplemental files updated. One author added.

