## Supplementary material for "A fuzzy sequencer for rapid DNA fragment counting and genotyping": The Supplementary Information

### 1. Materials and Methods

#### *1.1 Ethics and sample collection*

The lambda phage genomic DNA was obtained from New England Biolabs (NEB). Peripheral blood DNA samples of one congenital heart disease (CHD) patient was obtained from a previous CHD study, which was approved by the Medical Ethics Committee of Tsinghua University. Peripheral blood plasma cfDNA samples of pregnant women were obtained from a previous prenatal diagnostic study, which was approved by both Beijing Obstetrics and Gynecology Hospital and Tsinghua University. All samples were deidentified prior to use in this study.

#### *1.2 DNA extraction and library preparation*

*Genomic DNA sequencing library preparation.*

50 ng genomic DNA in 130 μL TE buffer was fragmented to ca. 180 bp using sonication (Covaris S220) with the following parameter setting: 175 W peak incident power, 10% duty factor, 200 cycles per burst, 300 s treatment time. Libraries were constructed from fragmented genomic DNA using NEBNext Ultra II DNA Library Prep Kit for Illumina (New England Biolabs) following manufacturer's protocol. Standard index PCR primers were replaced by custom dual-unique indexing primers for Illumina sequencing, or Adaptor A and P1 (synthesized by Thermo Fisher, Figure S8, Table S2) for Ion Torrent and BitSeq.

*Circulating cell-free DNA (ccfDNA) extraction.*

10 mL blood samples were centrifuged at 1600 g for 10 min at 4°C within 8 h after collection. The extracted plasma was centrifuged again at 16000 g for 10 min at 4°C to further remove cellular component, before being stored at -80°C . ccfDNA from the plasma was extracted using the CWhipro Circulating Nucleic Acid Kit (CW Biotech, China) following the manufacturer's instruction. The quantity and quality of the ccfDNA was checked by Qubit 3.0 (Thermo Fisher) and 2100 Bioanalyzer (Agilent).

*Mock trisomy ccfDNA preparation.*

Genome DNA from the trisomy patient was sheared with Covaris S220. The 150-250 bp part of the fragmented DNA was gel purified. The mock trisomy ccfDNA was prepared by mixing fragmented trisomy genomic DNA with ccfDNA extracted from nonpregnant female volunteers. The mixing ratios were validated through droplet digital PCR.

*ccfDNA sequencing library preparation.*

2-5 ng ccfDNA was end-repaired in a total 60 μL mix containing 1X T4 polynucleotide kinase buffer (Tiangen, China), 0.167 mM dNTP mix (NEB), 3U T4 DNA polymerase (Tiangen), 10U T4 polynucleotide kinase (Tiangen), 5U Klenow fragment (Tiangen) and sterile water at 25°C for 25 min, 70°C for 10 min and held in 4°C . The end-repaired DNA was then ligated in a total 100 μL mix with 1X T4 DNA ligase buffer (Tiangen), 3600U T4 DNA ligase (Tiangen), 40U Bst DNA polymerase (Tiangen), 50 nM Adaptor A and P1 at 25°C for 15 min, 65°C for 5 min and held at 4°C . The ligated DNA was cleaned up using 0.9X volume of Agencourt AMPure XP beads (Beckman Coulter). The library was sent to Beijing Jinnuo Ruijie Gene Science and Technology Co.,Ltd for ion torrent sequencing. This library was also for BitSeq.

*RNA extraction and reverse-transcription.*

Total RNA was extracted from 106 feeder-free mouse ES cells or MEF cells separately using RNeasy Micro kit (Qiagen) following the manufacturer's instructions. On-column DNase digestion was performed to remove genomic DNA. The quantity and quality of the RNA was checked using a Qubit RNA HS Assay kit (Invitrogen) and 2100 Bioanalyzer. Messenger RNA was isolated and fragmented from 1μg of extracted total RNA using NEBNext Poly(A) mRNA Magnetic Isolation Module, followed by reverse-transcription using NEBNext Ultra II RNA First and Second Strand Synthesis Module according to the manufacturer's instructions. Libraries were constructed from 10 ng double strand cDNA obtained above using NEBNext Ultra II DNA Library Prep Kit for Illumina following manufacturer's protocol. Standard index PCR primers were replaced by custom dual-unique indexing primers for Illumina sequencing, or Adaptor A and P1 for Ion Torrent and BitSeq. After 10 cycles PCR amplification, the final libraries were quantified using a Qubit dsDNA HS Assay kit (Invitrogen) and Agilent Fragment Analyzer.

*Targeted metagenomics sequencing (tNGS)*

Briefly, 600 μL BALF isolated from each enrolled patient was collected and mixed with lysozyme and glass beads. DNA extraction was performed by using the TIANamp Micro DNA Kit (TIANGEN BIOTECH, Beijing, China). The libraries were prepared using the tNGS panel of Cygnus Biosciences following the manufacturer's protocol.

#### *1.3 Sequencer design*

A 10× microscope objective (NA 0.45, CFI Plan Apo Lambda, Nikon) and an sCMOS sensor (Flash 4.0, Hamamatsu) were used for image acquisition in the sequencer prototype. A high-power blue LED with wavelength around 460 nm (CBT-90-B-TE, Luminus) was used for excitation. Both excitation (482±20 nm) and emission (536±40 nm) bandpass filters were purchased from Semrock. The x-axis translation stage (MSMF012, Panasonic) had 2 μm precision, taking ≈ 1 s to move 1.33 mm in each step. The y-axis translation stage (IKO) had 1 μm precision and 65 mm moving range, taking ≈ 300 ms to move 1.33 mm in each step. We imaged 171 tiles (9 by 19) in each reaction cycle to cover the whole reaction chamber. Reagent flow was driven by a syringe pump (Cavro XC, Tecan) and a series of solenoid valves (LVM10R6, SMC). The temperature of reaction was controlled by a Peltier device (TE tech), with a water-cooling system connected to the heat-sink. We used LabVIEW (National Instruments) to control the whole prototype.

#### *1.4 Synthesis of Peking Orange (***PO***)*

*Synthesis of (2-bromo-5-methoxyphenyl)(3-methoxyphenyl)methanol (*1a *in* Figure S23*).*

In a 250 mL round bottom flask (flame dried) equipped with a constant pressure dropping funnel, 80 mL anhydrous tetrahydrofuran solution of m-methoxyphenyl magnesium chloride (1.2 eq) was added and cooled to -40˚C with stirring. Then 20 mL anhydrous tetrahydrofuran solution of 3-methoxy-o-bromobenzaldehyde (10.8g) was added dropwise to the reaction flask through dropping funnel while kept at -40˚C with stirring. After addition the reaction was kept for 2-6 hours, monitored by TLC, and stopped when the raw material 3-methoxy-o-bromobenzaldehyde disappeared. Add 20 mL saturated ammonium chloride aqueous solution to quench the reaction. The reaction mixture was subjected to a rotary evaporator to remove most of the solvent, and then extracted twice with dichloromethane (100 mL each). After separation, the organic phases were combined, washed with saturated brine, dried over $Na_2SO_4$, concentrated by a rotary evaporator, and then purified by silica gel column chromatography with developing solvent: petroleum ether/ethyl acetate ~10/1, collect Rf~0.3 product to get 15 g colorless liquid 1a in 93% yield.

$^{1}$H NMR ($CDCl_3$, 500 MHz): δ 7.39 (d, J=10 Hz, 1H, Ar-H), 7.23 (t, J=10 Hz, 1H, Ar-H ),7.13 (d, J=5 Hz, 1H, Ar-H ), 6.96 (m, 1H, Ar-H), 6.81-6.79 (m, 1H, Ar-H), 6.96 (m, 1H, Ar-H), 6.70 (dd, J=10 Hz , 5 Hz, 1H, Ar-H), 6.08 (d,1H, CH), 3.77 (s, 3H), 3.75 (s, 3H).

$^{13}$C NMR (126 MHz, $CDCl_3$): δ 159.70, 159.29, 143.76, 143.48, 133.41, 129.51, 119.33, 115.12, 114.01, 113.11, 113.05, 112.80, 74.63, 55.49, 55.23.

HRMS: Calcd for $C_{15}H_{15}BrO_3Na$ (M+Na), 345.0097. Found, m/z 345.0096.

*Synthesis of (2-bromo-5-methoxyphenyl)(3-methoxyphenyl) methanone (*1b *in* Figure S23*).*

Dissolve 1a (6.5g) in 80 mL of dichloromethane in a 250 mL flask, add 10 g of pyridinium chlorochromate (PCC) and 10 g celite to the solution and the mixture was kept for 2-5 hours under rapid stirring at room temperature. The reaction is monitored by TLC until the raw materials disappear. The reaction solution is filtered through a short column pad with celite and washed with dichloromethane. The filtrate is collected, concentrated on rotary evaporator and subjected to silica gel column chromatography. Developing solvent: petroleum ether/ethyl acetate~10/1, Rf~0.6. 6.1g light yellow solid is obtained after concentration, 95% yield.

$^{1}$H NMR ($CDCl_3$, 500 MHz): δ 7.50 (d, J=10 Hz, 1H, Ar-H), 7.45 (m, 1H, Ar-H), 7.35 (d, J=10 Hz, 1H, Ar-H), 7.31-7.29 (m, 1H, Ar-H), 7.16-7.14 (m, 1H, Ar-H), 6.91-6.88 (m, 1H, Ar-H), 6.87 (d, J=5 Hz, 1H, Ar-H ), 3.85 (s, 3H), 3.79 (s, 3H).

$^{13}$C NMR (126 MHz, $CDCl_3$) δ□195.40, 159.92, 158.74, 141.48, 137.29, 133.93, 129.61, 123.52, 120.47, 117.36, 114.18, 113.76, 109.69, 77.05, 55.65, 55.49.

HRMS: Calcd for $C_{15}H_{14}BrO_3$ (M+H), 321.0121. Found, m/z 321.0120.

*Synthesis of 1-bromo-4-methoxy-2-(2-(3-methoxyphenyl)propan-2-yl) benzene (*1c *in* Figure S23*).*

70 mL of dichloromethane was added to a dry 250mL round-bottomed flask and then cooled to -40°C. Then $TiCl_4$ (9 mL) and dimethyl zinc solution (80 mL of 1M toluene solution) were slowly added respectively with stirring under the protection of argon. After stirring at 40°C for 15 minutes, 30ml of dichloromethane solution of 1b (6.1g) was added dropwise to the above solution and kept at -40°C for 3 hours. After that the temperature was slowly raised to 0°C and the reaction was continued for 5-10 hours. The reaction was monitored by TLC and stopped when raw materials 1b disappeared. The brown reaction solution was quenched by pouring into crushed ice. The mixture was extracted with dichloromethane, dried over $Na_2SO_4$, concentrated on a rotary evaporator and purified by silica gel column chromatography with developing solvent: petroleum ether/ethyl acetate ~15/1. Collect Rf~0.6 product and concentrate to obtain 5.5 g colorless liquid 1c with yield of 87%.

$^{1}$H NMR ($CDCl_3$, 500 MHz): δ 7.38 (d, *J*=10 Hz, 1H, Ar-H), 7.21 (d, *J*=5Hz, 1H, Ar-H),7.18-7.14 (m, 1H, Ar-H), 6.87-6.81 (m, 1H, Ar-H), 6.72-6.69 (m, 1H, Ar-H), 6.63 (dd, *J*=10 Hz , 5 Hz, 1H, Ar-H), 3.82 (s, 3H), 3.74 (s, 3H), 1.73 (s, 6H).

$^{13}$C NMR (126 MHz, $CDCl_3$): δ 159.51,158.70, 151.23, 148.67, 135.90, 133.52, 128.94, 119.03, 115.78, 113.10, 111.93, 109.87, 55.38, 55.14, 44.88, 30.16, 13.29.

HRMS: Calcd for $C_{17}H_{20}BrO_2$ (M+H), 335.0642. Found, m/z, 335.0641.

*Synthesis of 1-bromo-4-methoxy-2-(2-(3-methoxyphenyl)propan-2-yl) benzene (*1d *in* Figure S23*).*

Compound 1c (3g) in 30 mL anhydrous tetrahydrofuran was added into a dry 100 mL round-bottomed flask, cooled to -78°C under argon. *t*-butyllithium n-hexane solution (9~10 mmol) was added *with extremely caution* by well dried syringe. The solution was kept stirring at -78°C~-60°C for 1 hour. Then 10 mL anhydrous tetrahydrofuran solution of *o*-methyl benzaldehyde (11.5 mmol) was slowly add to the reaction at -78°C with syringe pump. After the addition is completed, the temperature was allowed to increase slowly with stirring for 2-6 hours and monitored by TLC. The reaction was quenched by adding 10 mL of saturated ammonium chloride solution. After removing most of the solvent by rotary evaporator, the residue mixture was extracted three times with dichloromethane (100 mL each), the organic phase was collected, dried over $Na_2SO_4$, and concentrated to obtain the crude compound 1d, which was directly used in the next reaction step.

*Synthesis of 1-bromo-4-methoxy-2-(2-(3-methoxyphenyl)propan-2-yl) benzene (*1e *in* Figure S23*).*

Dissolve the compound 1d from the previous step in 40 mL of dichloromethane, 4g of pyridinium chlorochromate (PCC) was added under rapid stirring. The reaction was kept for 2-4h at room temperature, monitored by TLC until the raw material disappears. The reaction solution is filtered through a short column pad with celite aid and washed with dichloromethane. The filtrate is collected, concentrated on rotary evaporator and subjected to silica gel column chromatography. Developing solvent: petroleum ether/ethyl acetate ~ 7/1, collect Rf ~ 0.5 product. After concentration, 2.9 g of yellowish solid was obtained as crude compound 1e, which was directly used in the next reaction step without further purification needed.

*Synthesis of 1-bromo-4-methoxy-2-(2-(3-methoxyphenyl)propan-2-yl) benzene (*PO*).*

Compound 1e (2.9g) from above step was dissolved in 40 mL anhydrous dichloromethane under ice-water bath cooling, and boron tribromide (2-5eq) was added dropwisely (*extremely caution*). The reaction was kept for 2~5 hours after the addition is completed. Then, 20 mL of ice water was carefully added to quench the reaction with stirring for 30 minutes. After that, the reaction was adjusted to pH ~7.0 with saturated aqueous $NaHCO_3$, extracted twice with 150 mL of dichloromethane, washed the organic phase with water and saturated brine, and dried over anhydrous $Na_2SO_4$. The organic phase was concentrated by rotary evaporator, and then drained by a vacuum oil pump under reduced pressure to obtain a slightly yellow oil. Then 5 mL of Methanesulfonic acid was added to the oil and heated to 80~100°C with stirring. The reaction was stopped and cooled to room temperature after 1 hour heating. The reactant was poured into crushed ice while stirring, the precipitated solid was collected by filtration, washed with water, and dried under vacuum to obtain about 2.2 g of the crude product of the target product 1. The crude product was separated and purified by silica gel column chromatography to obtain 1.8 g of orange-red solid "Peking Orange" (PO) with yield of 72%.

$^1$H NMR ($CDCl_3$, 500 MHz): δ 7.32-7.29 (m, 1H, Ar-H), 7.24-7.18(m, 2H, Ar-H), 7.02(d, *J*=5 Hz, 1H, Ar-H), 6.95(s, 1H, Ar-H), 6.94(s, 1H, Ar-H), 6.86 (d, *J*=10 Hz , 1H, Ar-H), 6.50 (dd, J=10 Hz , 5 Hz, 2H, Ar-H), 1.94 (s, 3H), 15.2 (s, 3H), 1.48 (s, 3H).

$^{13}$C NMR (126 MHz, $CDCl_3$) δ 174.51, 157.24, 155.28, 136.62, 135.96, 130.24, 129.10, 128.84, 125.72, 122.37, 120.66, 118.89, 114.02, 46.98, 40.15, 32.50, 31.93, 19.52.

HRMS: Calcd for $C_{23}H_{19}O_2$ (M-H), 327.1386. Found, m/z 327.1390.

#### *1.5 Chip preparation and sequencing*

*Etching of fiber optical plates.*

Fiber optical plates (FOPs) were fabricated by Guangzhou Honsun Opto-Electronic Co., Ltd. The etchant was prepared by adding 85 mL concentrated hydrochloric acid and 20 mL hydrogen peroxide to 895 mL water. One face of the FOP chip was covered by a piece of adhesive tape, while the other face was exposed to etchant. Etching was operated at 25°C for 2 h, then the chip was rinsed by water and air-dried. A thin layer (ca. 100-400 nm) of silica was deposited on the etched FOP by ion-beam deposition (IBD).

*Silane deposition.*

The FOP chips were firstly cleaned by air plasma, and then put into a vacuum oven for hydrophobic modification under argon atmosphere and trichloro(1H, 1H, 2H, 2H-perfluorooctyl)silane vapor under 100°C and 100 Pa for 1 h. Then the chips were cooled to room temperature and transferred to another vacuum oven under argon atmosphere and (3-mercaptopropyl)tri-methoxysilane vapor at 100°C and 100 Pa for 30 min. Finally, the chips were put on a hot stage of 130°C to age the silane modification.

*PEG and streptavidin coating.*

The silane-coated chips were rinsed by isopropanol and then 1X PBS buffer. The 0.45 mM maleimide-PEG-biotin (M.W. 5000, Laysan Bio) solution was injected into the chip, incubated for 30 min and washed by 10 mL MilliQ water. Then 100 μg/mL streptavidin (Sigma Aldrich) in 1X PBS buffer was injected into the chip, incubated for 10 min, and washed by 10 mL 1X PBS buffer.

*Preparation and loading sequencing beads.*

The Ion Sphere Particles (ISPs) were prepared through emulsion PCR and enrichment following the Ion Torrent OneTouch2 protocol (ThermoFisher). Then, 100 μL ISP solution dispersed in 1× TdT buffer was treated by TdT solution and 10μM biotin-16-dUTP (NEB) at 37°C for 1 h, followed by adding 100 μM ddNTP at 37°C for 4 h to further block all free 3' DNA terminals. After that the ISP solution was diluted to 0.3X by 1X PBS, and injected into the sequencing chip. The chip was centrifuged under 1000 g for 10 min and washed by wash buffer to remove ISPs that were not firmly immobilized on-chip.

*Sequencing.*

Sequencing was done by sequentially introducing dual-base reaction mix (the dual-base flowgram) as we have described before (Ref. 15). The sequencing reaction mix contains Bst polymerase, calf intestinal alkaline phosphatase (CIP, NEB), $MnCl_2$ (Sigma Aldrich) and two types of fluorogenic nucleotides to provide specific degenerate combinations. In each sequencing cycle, the chip was filled with the sequencing reaction mix at 4°C and then sealed by fluorinated oil (FC-40, 3M), the sequencing-by-synthesis reaction was triggered by heating the sealed chip to 65°C for 30 s. Then the chip was cooled to 25°C to stop the reaction and to perform the tiling-image acquisition. The chip was then washed by isopropanol and wash buffer at the end of each reaction cycle. The sequencing operation was controlled by a LabVIEW program.

#### *1.6 Base-calling*

The microwells on the FOP were imaged as multiple bright dots with different intensities in the fluorescent images. For each fluorescent image, the microwells were recognized as local maxima by image morphological dilation. Then the images were up-scaled to increase the resolution. The microwell positions were refined by weighted average of pixels around the local maxima on the resized image. The intensities of the microwells were extracted by sum of pixels around the refined positions on the resized image. The extracted intensities were further corrected by subtracting intensities of neighboring dark microwells. Light stains and dark stains were identified as large areas with over bright or over dark intensities, and microwells from stained regions were discarded in the following steps (Figure S6A). Images of the same tile in different cycle were registered through the positioning markers on the bright-field image (Figure S6B).

We plotted a histogram for microwell intensities extracted from each cycle (Figure S10A). Two thresholds were identified in the histogram to separate the first peak as the dark dots, the second peak as the monoclonal bright dots and the rest as polyclonal bright dots. Only monoclonal bright dots were retained for further processing.

We mixed two kinds of DNA samples in each run: the standard dots (SD) and library dots (LD). All SDs have known identical sequences while LDs are from the sample to be sequenced and have variant sequences. All LDs have an identical starting sequence GTAGCC (the key sequence in Figure S8) which is different from SDs. We calculated the Pearson's correlation coefficients between intensities of Cycle 1-6 and the degenerate polymer length (DPL) of LD/SD (Figure S10B). Dots with correlation>0.9 with the LD DPL were identified as LD, and dots with correlation>0.95 with the SD DPL were identified as SD. Dots that were neither LD nor SD were identified as error dots (ED).

The SD intensities $f$ were corrected from dephasing using our previous algorithm (Ref. 15,18). Briefly, we constructed a flux matrix $T$ from the lead $(\varepsilon_1, \varepsilon_2)$, lag $(\lambda_1, \lambda_2)$ and SD DPL $h$ to transform $h$ to a dephased signal $s$, where $\varepsilon_1$ and $\lambda_1$ are for odd cycles and $\varepsilon_2$ and $\lambda_2$ are for even cycles. Then the predicted intensities was $f^* = a \cdot (1-b)t \cdot s + c \cdot s_1 + d \cdot s_2,$ where $a$ is the unit signal, $b$ is the decay coefficient, $c$ is hydrolysis1 (hydrolysis of odd cycles), $d$ is hydrolysis2 (hydrolysis of even cycles), $s_1$ is 1 in odd cycles and 0 otherwise, $s_2$ is 1 in even cycles and 0 otherwise. We minimized the lost function $y = \|f - f^*\|^2 + r^2[(\varepsilon_1 - \varepsilon_2)^2 + (\lambda_1 - \lambda_2)^2]$ to fit all the parameters $(\varepsilon_1, \varepsilon_2, \lambda_1, \lambda_2, a, b, c, d)$. The regularization coefficient $r$ was set to $10^9$. The corrected signal was obtained by reverse transformation of intensity $f$ to DPL $h$ using the fitted parameters.

The LD intensities were first normalized by the known DPLs of their key sequence and then corrected from dephasing using the mean dephasing coefficients obtained from SD. The corrected DPL of LD were rounded to their nearest integers and then transformed to bit sequences.

The dephasing correction of SuperBitSeq is similar to that of BitSeq, except for that the signals from the two channels are processed separately, according to our previous simulation results (Ref. 18).

#### *1.7 Mapping the simulated error-free bit sequences to the genome*

To test the accuracy of the mapping of bit and superbit sequences, we simulated the mapping of error-free sequences to three genomes: *Homo sapiens* (GRCh38, or hg38), *Arabidopsis thaliana* (TAIR10.1) and a simulated random genome. The simulated genome has 3 $\times 10^9$ bases equally distributed in 20 mock chromosomes (150Mb each) since BWA can only process chromosomes less than 2Gb. The simulated genome is fully constituted of base A, C, G and T without any ambiguous bases such as N. Each base has equal occurrence probability of 0.25 and the base type in each site of the genome is independent from any other sites. $1 \times 10^7$, $1 \times 10^6$ and $1 \times 10^7$ sites were uniformly sampled from the GRCh38, TAIR10.1 and the simulated genome, respectively. Regions with ambiguous bases (marked as N in the reference genome) were omitted in the sampling. For each site, by retrieving sequences with different lengths, we simulated sequencing reads under the MK, RY, WS and CRT flowgram with 5-150 cycles, respectively (Figure S20). BitSeq and SuperBitSeq flowgrams were named with a heading 'b' and 's', respectively. For example, bMK stands for BitSeq in MK flowgram, and sRY stands for SuperBitSeq in RY flowgram. This nomenclature is consistent through this manuscript. No errors were added to these sequences. Sequencing signals were encoded as both bit and superbit sequences according to the method described in section 2.1. These sequences were mapped to their corresponding reference genomes using Bowtie2, BWA-MEM or BWA-SW, respectively. We calculated the percentage of the following kinds of sequences from the BAM files (Table S7-S24, S47-S52):

1) Total mapping: mapped to the reference genome by the software;
2) Unique mapping: no XS tag, or the value of XS tag is less than that of AS tag;
3) Q20 mapping: the mapping quality is no less than 20;
4) Q30 mapping: the mapping quality is no less than 30;
5) Correct mapping: the distance between the mapped site and the sampling site is no greater than 5 bp;
6) Unique and correct mapping: both unique and correct mapping.

#### *1.8 Mapping the simulated erroneous sequences to the human genome*

To test the accuracy of the mapping of bit sequences with a few errors, $1 \times 10^6$ sites were uniformly sampled from the hg38. Regions with ambiguous bases were omitted in the sampling. For each site, we also simulated sequencing reads under the MK and RY flowgram with 15-150 cycles, respectively, as in the error-free sequence simulation. Erroneous sequences under the CRT flowgram were not simulated because they have different error profiles compared to bit sequences (more substitutions than indels). To simulate errors in bit sequences, we used a matrix $P_{n\times n}$ to describe the error pattern, in which $P_{ij}$ is the probability that DPL $i$ is sequenced as $j$. $P_{n\times n}$ was generated as follows:

1. $$P_{ij} = \begin{cases} \frac{\lambda}{i+2\lambda} + \frac{1}{2} & i = j \\ \left(\frac{P_{ii}-1}{P_{ii}-2}\right)^{|i-j|} & i \neq j \end{cases}$$

2. Normalize $P$ such that the sum of each row is 1.

$n$ was set to 50, an integer sufficiently greater than most DPLs. $\lambda$ was set to 447.6, 222.6, 110.1, 53.8 and 25.7, corresponding to error rate of 0.25%, 0.5%, 1%, 2% and 4%, respectively. In the simulation, DPLs were transformed from the sampled bit sequences, then modified according to the probability given by the matrix $P$. For example, the probability that DPL $i$ remains unchanged is $P_{i,i}$, and the probability that DPL $i$ is erroneously sequenced as $i + 1$ is $P_{i,i+1}$, etc. The modified DPLs were transformed back to bit sequences and mapped to the reference genome using Bowtie2, BWA-MEM or BWA-SW, respectively. The percentage of total mapping, unique mapping, Q20 mapping, Q30 mapping, correct mapping and unique and correct mapping reads (Table S25-S42) were calculated using the same method in the previous section that mapping the simulated error-free sequences.

#### *1.9 Mapping simulated random sequences to the human genome*

As a negative control, random sequences were used to validate that non-genome bit sequences would not be erroneously mapped onto the genome. We generated $1\times10^6$ random DNA sequences of 1 kb. The probability of each 4 kinds of base is 0.25 and each site was generated independent from each other. For each random sequence, by retrieving subsequences with different lengths, we simulated sequencing reads under the MK, RY and CRT flowgram with 5-50 cycles, respectively, as in the error-free sequence simulation. These sequences were mapped to their corresponding genomes using Bowtie2, BWA-MEM or BWA-SW, respectively. The percentage of total mapping, unique mapping, Q20 mapping and Q30 mapping reads were calculated (Table S43-46) using the same method as previously described. Correct mapping or unique and correct mapping reads were not calculated since random sequences should not have been mapped.

#### *1.10 Whole-genome screening of faithful mapping sites of human genome*

We defined *faithful mapping* as:

1) the sequence was mapped to where it was generated (correct mapping);
2) the mapping quality is no less than 20 (Q20 mapping);
3) no XS tag, or the value of AS tag minus that of XS tag is no less than 30 (unique mapping);

We enumerated every 50, 70 and 100 bp sequences on the GRCh38. For example, in the case of 50 bp, the sequences were from sites of 1-50, 2-51, 3-52, etc, till the end of each chromosome. These sequences were transformed into MK and RY bit sequences, and mapped to their corresponding reference genome along with their original DNA sequences using Bowtie2 and BWA-MEM. We selected sites where the generated sequences can be faithfully mapped as the faithful mapping sites. The number of faithful mapping site of different read length and software can be seen in Table S53-58.

#### *1.11 Analysis of lambda phage genomic DNA sequencing*

The bit sequences from the lambda phage genomic DNA were mapped to its corresponding encoded reference genome using BWA-MEM with the following parameters: -B 5 -O 3 -E 3. Only reads with mapping quality no less than 20 were used for calculating the error rate, using the method described by Ref. 33. Soft-clipped bases were omitted during the calculation.

#### *1.12 Identification of copy number variations (CNVs)*

1) *Generating the BED files.*

We firstly screened for the faithful mapping sites (see section 1.10). We then retained faithful mapping sites that occur consecutively for over 200 bp and discard the rest sporadic faithful mapping sites. We then transformed the retained sites to BED form files. Two different BED files were generated for normal 4-base DNA sequences and bit sequences, respectively.

2) *Binning.*

The reference genome is divided into multiple bins in which every bin contains exactly L faithful mapping sites recorded in the BED files. If there are less than L sites when binning at the end of a chromosome, discard this bin. The binning resolution L is 2 Mb for identification of aneuploidies and 100 kb for small CNVs.

3) *Filtering.*
Sequencing reads are mapped to the reference genome using the default settings of BWA-MEM and retain reads that:

a) are mapped to regions recorded in the BED files;
b) with mapping quality no less than 20;
c) contain no XS tag in the alignment result, or the value of AS tag minus XS tag is no less than 5.

3) *Counting.*
Count the number $N_i$ of reads mapped to each bin of the reference genome as the sequencing depth.

4) *GC correction.*
Suppose the GC content of each bin is $g_i$, use the locally-weighted polynomial regression (LOWESS) to fit the curve of $\log(N_i)$ with respect to $g_i$. Suppose the fitted values are $f_i$, calculate $x_i = \exp(\log(N_i) - f_i)$ as the sequencing depth after GC correction.

5) *Normalization.*
Calculate the mean value $\bar{x}$ of $x_i$ on all autosomes, and calculate $x_i' = 2x_i/\bar{x}$ as the normalized depth of each bin.

6) *CNV identification.*
Use the circular binary segmentation (CBS) algorithm of the DNAcopy package[34] in the R language to calculate the segment mean $m_i$ from $x_i$. The segment mean $m_i$ is the copy number in this bin.

#### *1.13 Noninvasive prenatal testing (NIPT)*

The bioinformatic pipeline for NIPT is:
1) Use the same pipeline as the CNV identification to get the normalized depth $x_i'$ using a binning resolution of L = 2 Mb.
2) Calculate the representative value of Chr21 of each sample as

$$y_{chr21} = \frac{\sum_{chr21} x_i'}{\sum_{autosome} x_i'}$$

3) Calculate the mean $\mu_{chr21}$ and standard deviation $\sigma_{chr21}$ of the Chr21 representative values of all 24 negative controls, and the Chr21 Z-score of a sample $s$ to be detected is:

$$Z_{chr21}^{s} = \frac{y_{chr21}^{s} - \mu_{chr21}}{\sigma_{chr21}}$$

4) The Z-scores for Chr18 and Chr13 are calculated in the same manner. A Z-score greater than 3 is tested as trisomy positive, otherwise negative.

The bioinformatic method for gender classification is:

1) Use the same pipeline as the CNV identification to get the normalized depth $x_i'$ using a binning resolution of L = 2 Mb.

2) Calculate the mean depth of ChrY as:

$$d_{chrY}^{s} = \frac{1}{n_{chrY}} \sum_{chrY} x_i'$$

In the calculation of $d_{chrY}^{s}$, omit the second bin of ChrY for both BitSeq and conventional 4-base DNA sequencing, because this bin is akin to the centromere and its depth varies significantly.

#### *1.14 RNA-seq*

For conventional 4-base DNA sequences, the transcripts per kilobase million (TPM) of each gene were directly obtained by Salmon v1.4.0 [35].

For bit sequences, we first used the default settings of BWA-MEM to map the reads to the corresponding reference genome, retained reads with less than 2 errors and an alignment read length no less than 70 bp, then used Salmon v1.4.0 to obtain the TPM.

#### *1.15 metagenomics sequencing (mNGS)*

For the untargeted metagenomics sequencing, the raw sequencing data were filtered using fastp v0.23.2 to remove reads with length less than 50 bp or mean base quality less than 20, and trim reads over 100 bp to 100 bp. The data were further deduplicated using seqkit v2.3.0 and removed reads with >25 bp homopolymers. The filtered data were mapped to the human genome GRCh38 (MK encoded) using hs-blastn v0.0.5 with default parameters, and the unmapped reads were again mapped to RefSeq (downloaded in May 2022, MK encoded) using hs-blastn with the parameter -evalue 1e-30 -outfmt 6 -dust no -word_size 50. For each read, only the query with the highest score was retained for species identification. For targeted metagenomics sequencing (tNGS), the sequences of targeted regions of each species were compiled to form the reference sequences. The sequencing reads were mapped to these reference sequences using BWA-MEM (v0.7.17-r1188). The read numbers mapped to each species are counted using a bespoke Python script and normalized by the total read number with unit in 100k reads.

### 2. Supplementary Text

#### *2.1 Read length per cycle and information efficiency of different sequencing strategies*

Assume that the bases in the DNA molecules sequenced are independent and identically distributed:

$$P(base = A) = P(base = C) = P(base = G) = P(base = T) = 0.25$$

We have reported the read length and information entropy per cycle of CRT, SNA, BitSeq and SuperBitSeq in Ref. 15. Briefly:

1) The cyclic reversible terminator (CRT) sequences exactly one base per cycle, so its read length per cycle is:

$$L_{CRT} = 1\ bp/cycle$$

The intrinsic information efficiency is:

$$H_{CRT} = -4 \cdot \frac{1}{4} \log_2 \frac{1}{4} = 2\ bit/cycle$$

2) The single-nucleotide addition (SNA) sequences one homopolymer per cycle, so its read length per cycle is:

$$L_{SNA} = \frac{1}{2} \cdot \sum_{i=1}^{\infty} i \cdot \frac{3}{4^i} = \frac{2}{3}\ bp/cycle$$

The intrinsic information efficiency is:

$$H_{SNA} = 2L_{SNA} = \frac{4}{3}\ bit/cycle$$

3) The signal of BitSeq follows the distribution $P(x = i) = \frac{1}{2^i}$, so its read length per cycle is:

$$L_{bit} = \sum_{i=1}^{\infty} i \cdot \frac{1}{2^i} = 2\ bp/cycle$$

The intrinsic information efficiency is:

$$H_{bit} = \sum_{i=1}^{\infty} \frac{\log_2 2^i}{2^i} = 2\ bit/cycle$$

4) SuperBitSeq has the same read length per cycle as BitSeq:

$$L_{superbit} = L_{bit} = 2\ bp/cycle$$

The signal of SuperBitSeq follows the distribution $P(x = i, y = j) = \frac{C_{i+j}^{i}}{4^{i+j}}$, so its intrinsic information efficiency is:

$$H_{superbit} = -\sum_{i=0,j=0,i+j>0}^{\infty} \frac{C_{i+j}^{i}}{4^{i+j}} \cdot \log_2 \frac{C_{i+j}^{i}}{4^{i+j}} \approx 3.37\ bit/cycle$$

5) Trit, SuperTrit and HyperTritSeq added three different bases in each cycle, i.e., B (C/G/T), D (A/G/T), H (A/C/T), V (A/C/G). In TritSeq, the three bases in the same cycle are labeled with the same dye. In SuperTritSeq, one of the three bases is labeled with the first kind of dye while the other two bases are labeled with the second kind of dye. In HyperTritSeq, the three bases are each labeled with a different dye. For Trit, SuperTrit and HyperTritSeq, the number of bases extended in each cycle follows the distribution:

$$P(x = i) = \frac{1}{3} \cdot \left(\frac{3}{4}\right)^i$$

So their read length per cycle is:

$$L_{trit} = L_{supertrit} = L_{hypertrit} = \sum_{i=1}^{\infty} \left[ i \cdot \frac{1}{3} \cdot \left(\frac{3}{4}\right)^i \right] = 4\ bp/cycle$$

The intrinsic information efficiency of TritSeq is:

$$H_{trit} = -\sum_{i=1}^{\infty} \left( \frac{3^{i-1}}{4^i} \cdot \log_2 \frac{3^{i-1}}{4^i} \right) = 8 - 3\log_2 3 \approx 3.25\ bit/cycle$$

For SuperTrit and HyperTritSeq, there is a hidden fact that the first base extended in one cycle can be inferred from the previous cycle. For example, we add B (C/G/T) in cycle 1 and D (A/G/T) in cycle 2, then the first base extended in cycle 2 must be A, otherwise the nascent DNA strand will not stop there in cycle 1.

Given that, suppose $n$ bases were extended in one cycle of SuperTritSeq, $n = 1 + a + b$, where 1 denotes the first known base, $a$ and $b$ denote the number of rest bases labeled with a green and a red dye, respectively. The probability distribution of a SuperTritSeq signal is:

$$p_{a,b} = \frac{1}{3} \cdot \left(\frac{3}{4}\right)^{1+a+b} \cdot \frac{C_{a+b}^{a}}{2^{a+b}} = \frac{C_{a+b}^{a}}{4} \cdot \left(\frac{3}{8}\right)^{a+b}$$

So the intrinsic information efficiency of SuperTritSeq is:

$$H_{supertrit} = -\sum_{a\geq 0, b\geq 0} \left(p_{a,b} \cdot \log_2 p_{a,b}\right) \approx 4.59\ bit/cycle$$

Likewise, suppose $n$ bases were extended in one cycle of HyperTritSeq, $n = 1 + a + b + c$, where 1 denotes the first known base, a, b and c denote the number of the rest three kinds of bases, respectively. The probability distribution of a HyperTritSeq signal is:

$$p_{a,b,c} = \frac{1}{3} \cdot \left(\frac{3}{4}\right)^{1+a+b+c} \cdot \frac{C_{a+b+c}^{a} C_{b+c}^{b}}{3^{a+b+c}} = \frac{C_{a+b+c}^{a} C_{b+c}^{b}}{4^{1+a+b+c}}$$

The intrinsic information efficiency of HyperTritSeq is:

$$H_{hypertrit} = -\sum_{a\geq 0, b\geq 0} \left(p_{a,b,c} \cdot \log_2 p_{a,b,c}\right) \approx 5.59\ bit/cycle$$

The extrinsic information efficiency is defined as the information of DNA template sequenced per cycle, and is calculated as twice of the read length per cycle in the unit of bit/cycle. Hence the extrinsic information efficiency of CRT, SNA, BitSeq, SuperBitSeq, TritSeq, SuperTritSeq and HyperTritSeq is 2, 4/3, 4, 4, 8, 8, 8 bit/cycle, respectively.

#### *2.2 Coding strategy compatible with the prevalent mapping software*

Common DNA mapping software, such as Burrows-Wheeler Aligner (BWA) (Ref. 19,20) and Bowtie2 (Ref. 21,22) only recognizes natural bases A, C, G and T, and any other letters will be treated as 'N' in the sequence. It is critical to figure out a way that can encode the bit sequences to fit existing bioinformatic pipelines, and to check the performance of fuzzy sequencing. Fuzzy sequencing has been envisaged before without a feasible coding and mapping strategy (*36,37*). To map bit sequences onto the reference genome, both the bit sequences and the reference genome need to be encoded using the recognizable letter A, C, G or T. The critical point is how sequencing reads from the reverse-complement strand can also be mapped. For the sequenced reads, they are reverse-complementary themselves and then encoded. For the reference genome, it is encoded first and then transformed to reverse-complement. If reads from the reverse-complement strand are to be mapped, the reverse-complement operation and the encode operation must be commutative, i.e., a sequence reverse-complemented first and then encoded, or encoded first and then reverse-complemented, should result in the same sequence. This is called the commutative criterion. As bit sequences can have three possible ways to be encoded (MK, RY, or WS), we then consider each encoding scheme separately.

There might be two kinds of encoding strategies for the MK bit sequences (Table S3): (1) encode every M to A and every K to T, and (2) encode every M to A and every K to C. Only Strategy 1 satisfies the commutative criterion. The following encoding strategy for the RY bit sequence satisfies the commutative criterion: encode every R to A and every Y to T. However, the WS bit sequence is different (Table S4), and an additional strategy (the Strategy 3, which encodes every W to AT and every S to CG) is needed to satisfy the commutative criterion. Nevertheless, this encoding strategy will double the reference genome and make precise mapping more challenging and less practical.

Therefore, we developed a new approach for WS bit sequence to circumvent the commutative criterion. We put the original genome sequences and their reverse-complement together in a single fasta file, encoded every W to A and every S to C, then build the indices for the mapping software. In other words, there were both Chr1 and Chr1_rc in the genome. After mapping, we discarded alignments with flag=16. Reads from the reverse-complement strands should be mapped to chromosomes tagged with rc with flag=0. However, the commutative criterion allows bit sequence mapping in a fashion closer to conventional 4-base sequence mapping. Hence, we omitted the detail discussion of WS bit sequence in this manuscript.

The encoding strategy for superbit sequences of all MK, RY and WS is as follows: split the 4-base DNA sequence into groups according to the bases extended in each cycle, sort each group alphabetically, then rejoin the sorted groups. We gave an example of the sequence AGATGC in Table S5 and demonstrated that this encoding strategy follows the commutative criterion (Table S6).

An intuitive explanation of the commutative criterion is that the encoding strategy should keep the complement relationship of the two degenerate bases. M and K are complement, i.e., if base $X \in M = \{A, C\}$, then its complement base $X^* \in K = \{G, T\}$. So we should use two complement bases A and T to encode M and K. R and Y are also complement, so we use A and T to encode them, too. The complement bases C and G can also be used to encode MK or RY. WS is different because W is complementary to W itself and S is complementary to S itself, too. However, none of A, C, G or T is complement to itself, so neither A/T nor A/C coding of WS satisfies the commutative criterion. Strategy 3 in Table S4 satisfies the commutative criterion because the complement of sequence AT is still AT and the complement of sequence CG is still CG, thus keeping the complement relationship of W and S.

To summarize, the coding strategy we have used in this work is: (1) for MK, encode every M to A and every K to T, and (2) for RY, encode every R to A and every Y to T.

#### *2.3 The geometric interpretation of fuzzy sequencing*

The relationship between fuzzy sequencing and ECC sequencing can be interpreted in a geometric perspective.

Firstly, we represent a 4-base DNA sequence as a 4-dimensional line. We construct a 4-dimensional Euclidean space whose 4 unit orthogonal bases represent the 4 nucleotides, respectively: $\vec{A} = (1,0,0,0), \vec{C} = (0,1,0,0), \vec{G} = (0,0,1,0), \vec{T} = (0,0,0,1)$. Therefore, the vectors of degenerate nucleotides (M, K, R, Y, W, S) are the sum of vectors of their corresponding nucleotides, e.g., $\vec{M} = (1,1,0,0), \vec{K} = (0,0,1,1)$. For a given DNA sequence, we start from the original point $(0,0,0,0)$ and read the nucleotides one-by-one. Every time we read one nucleotide, we move towards the direction of this nucleotide's corresponding vector. After reading the whole sequence, we obtain a 4-dimensional line representing a DNA sequence.

Secondly, vector $\vec{M}$ and $\vec{K}$ span a plane $P_{MK}$. The projection of the 4-dimensional line on plane $P_{MK}$ represents the MK bit sequence of the DNA. Likewise, vector $\vec{R}$ and $\vec{Y}$ span the plane $P_{RY}$, vector $\vec{W}$ and $\vec{S}$ span a plane $P_{WS}$, and the projections of the 4-dimensional line on plane $P_{RY}$ and $P_{WS}$ represent the RY and WS bit sequences, respectively.

Then it would be obvious that ECC sequencing is actually attempting to reconstruct the 4-dimensional line using its projections on 3 orthogonal planes. This is similar to 3D reconstruction through multiple 2D projections (Ref. 38). On the other hand, BitSeq is actually attempting to infer the DNA sequence using one of its 2D projections. Albeit the projection doesn't contain the full information of the sequence, two sufficiently long bit sequences can be distinguished with the help of the reference genome.

The same viewpoint can be applied to SuperBitSeq as well. Taking the MK round as an example, suppose A and G are labeled with one dye (green) and C and T are labeled with the other dye (red). Then the geometric image of the MK superbit sequence is two projections of the 4D line representing the DNA sequence: one is the projection on the plane spanned by the vector $\vec{A}$ and $\vec{G}$, the other is on the plane spanned by the vector $\vec{C}$ and $\vec{T}$. Other rounds or labels of superbit sequences can be interpreted as two projections in the same way. SuperBitSeq is attempting to infer the DNA sequence using two of its projections. Similarly, two or three different rounds of superbit sequences can be decoded to a DNA sequence, in analogy to the 4D line reconstruction from 4 or 6 projections. The sequence decoding (or reconstruction) from superbit sequences is expected to be even more accurate than those from bit sequences.

#### *2.4 The construction and Hausdorff dimension of 'SuperBitSeq dust'*

The superbit sequences can be naturally presented as fractal patterns. Every infinite long DNA sequence can be mapped to a point in the square region $[0,1) \times [0,1)$ by two of its three bit sequences. For example, the MK and RY bit sequence of AAGCT⋯ are 11010⋯ and 11100⋯, respectively. We transform the two bit sequences into binary fraction numbers b0.11010⋯ and b0.11100⋯, then to decimal fraction numbers 0.8125⋯ and 0.8750⋯, so the sequence AAGCT⋯ can be mapped to the point $(0.8125\cdots,0.8750\cdots)$. If we choose MK and RY BitSeq as the X and Y axis, and only plot points mapped by sequences that are identical with their encoded MK or WS superbit sequences, we shall get the two fractals 'SuperBitSeq dust' depicted in Fig. 4F.

The shape of the SuperBitSeq dust is dependent on which bit sequences the two axes stands for and which superbit sequences to plot. If the type of superbit sequences to be plotted is one of the bit sequence types of the two axes, the shape of the fractal will be the top one of Fig. 4F (or its rotations). Otherwise, it will be the bottom one of Fig. 4F (or its rotations).

To calculate the Hausdorff dimension of the 'SuperBitSeq dust', first we calculate the number $x_n$ of n-bp superbit sequences. Taking MK as an example and let's consider the relationship between $x_{n+2}$, $x_{n+1}$ and $x_n$. We can add one base to the end of all (n+1)-bp superbit sequences and will get $4x_{n+1}$ different (n+2)-bp sequences. However, among the $4x_{n+1}$ sequences, there are two situations that are not superbit sequences:

1) Adding an A to the end of sequences ending with C;

2) Adding an T to the end of sequences ending with G;

So there are $2x_n$ sequences to be excluded from the $4x_{n+1}$ sequences. Hence we get the following relationship:

$$x_{n+2} = 4x_{n+1} - 2x_n$$

It is easy to know that $x_1 = 4, x_2 = 14$. Solving this equation, we get:

$$x_n = \frac{1}{2}\left[(1+\sqrt{2})(2+\sqrt{2})^n + (1-\sqrt{2})(2-\sqrt{2})^n\right]$$

Secondly, the rescaling factor of 'SuperBitSeq dust' (Fig. 4F) is 1/2, so its Hausdorff dimension is:

$$D = \lim_{n\to\infty}\frac{\log x_n}{\log 2^n} = \frac{\log(2+\sqrt{2})}{\log 2} \approx 1.77$$

Since the symmetry nature between MK, RY and WS, the Hausdorff dimensions of RY and WS are equal to that of MK.

Note that the Hausdorff dimension is close but not equal to the per-base information of SuperBitSeq:

$$\frac{H_{super-bit}}{L_{super-bit}} \approx \frac{3.37}{2} = 1.69$$

***2.5 SNV calling of <u>SuperBitSeq</u>***

Although the encoding strategy for superbit sequences satisfies the commutative criterion, it rearranges the original sequence and the SNV information may thus be changed. To reconstruct the SNV information, we implemented an SNV reconstruction algorithm:

1. When encoding the genome, we record the site map, i.e., the positions of encoded bases in the original genome. For example, the sequence ACAGTCCTG will be encoded as AACGTCCGT in MK, and the site map will be (1, 3, 2, 4, 5, 6, 7, 9, 8).
2. When encoding the sequencing signal into superbit sequences, the first 4 and last 3 cycles are discarded.
3. The sequence read is separated into two semi-sequences S1 and S2, each semi-sequence is composed of the two kinds of bases labeled with the same fluorophore.
4. The mapped region on the reference genome is retrieved and also separated into two semi-sequences R1 and R2 in the same manner as S1 and S2.
5. S1 and R1, S2 and R2 are locally aligned, respectively, using the Smith-Waterman algorithm.
6. The two local alignments of semi-sequences are merged together according to the site map.
7. The allele frequency of each site is calculated using pysam v0.15.4. Only SNVs with depth > 30 and allele frequency > 70% are called.

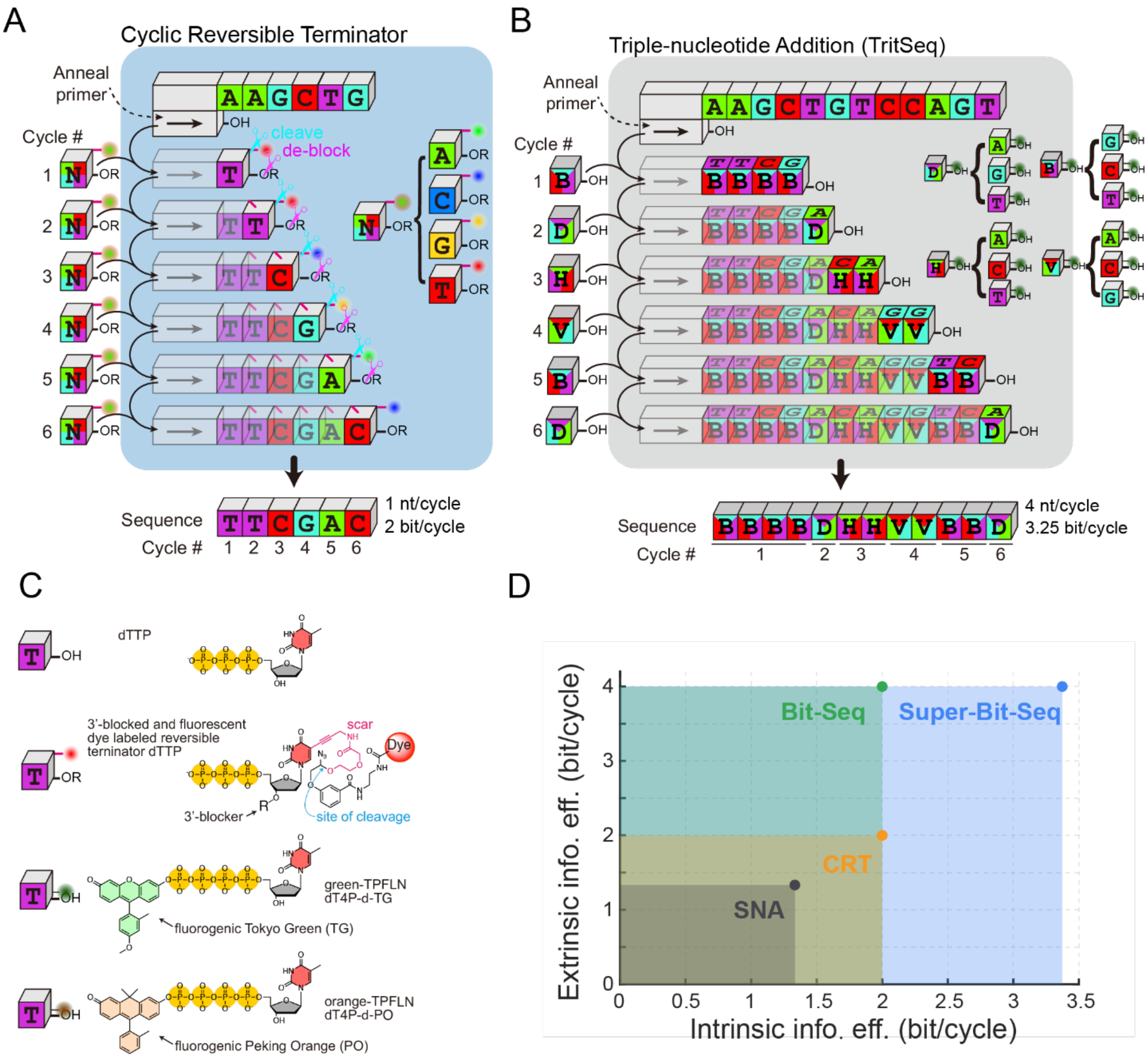


**Figure S1.** Schematic of commercial high-throughput sequencing chemistries. (A) Schematic of Cyclic reversible terminator (CRT) used by Illumina and MGI. (B) Schematic of triple-nucleotide addition (TritSeq). (C) Nucleotide substrates used in different sequencing chemistries. The TPLFN is used in fluorogenic sequencing, i.e., error-correction code (ECC) sequencing, BitSeq and SuperBitSeq. (D) The information efficiency of different sequencing technologies. Compared with SNA or CRT, BitSeq and SuperBitSeq show higher information efficiency that can be illustrated as the product of intrinsic and extrinsic components.

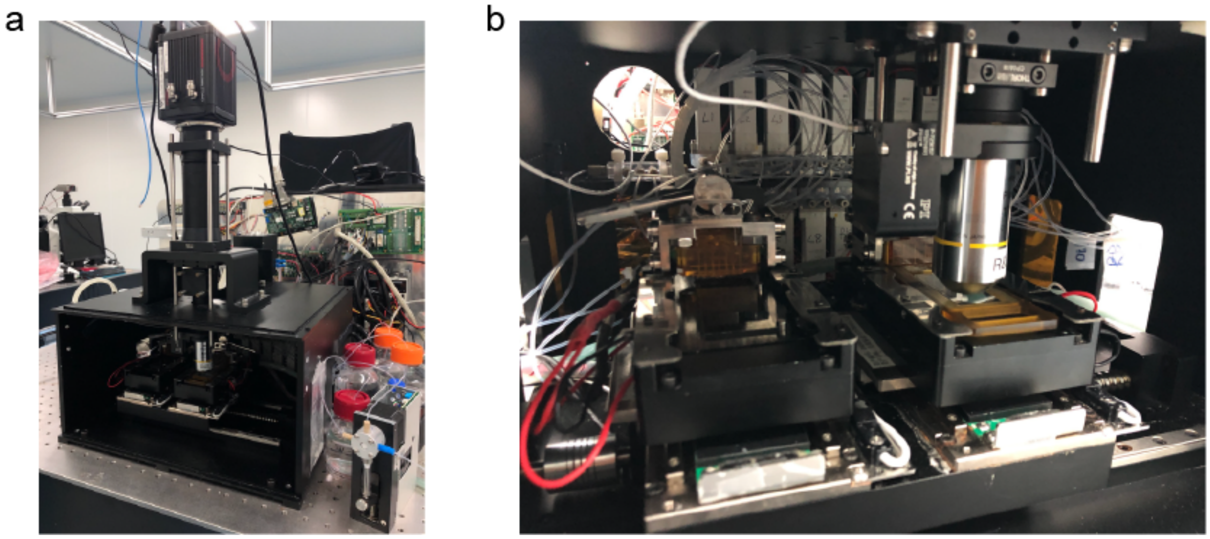


**Figure S2.** Photos of the sequencer prototype. (a) The ‘black-box’ prototype with major components including the camera, the imaging optics, the motion stages equipped with sequencing modules, as well as the liquid delivery. (b) Inside the ‘black-box’. Two sequencing modules sit on the moving stages. Above one of the sequencing module is the microscopic objective lens mounted on a piezo focusing adjustment stage. The manifolds are used for connection between the flow cell and the fluidic system (the selector and solenoid valves are shown in the photo).

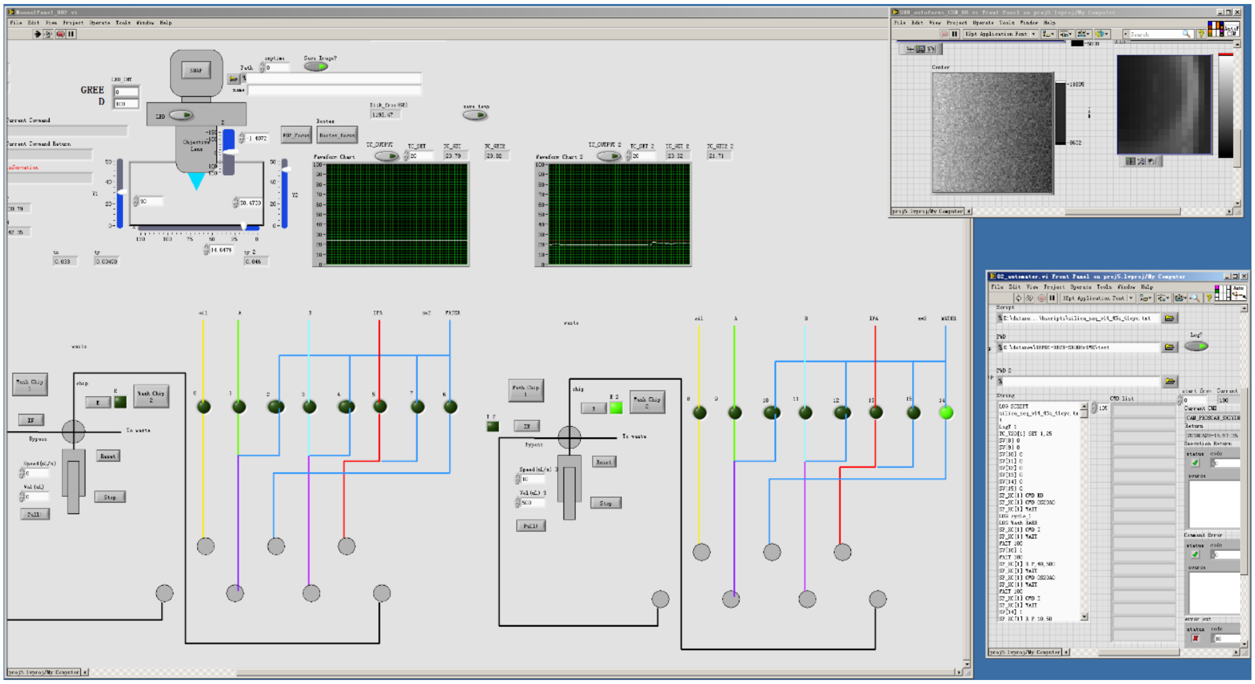

**Figure S3.** The graphical user interface of the sequencer operating software (written in LabVIEW).

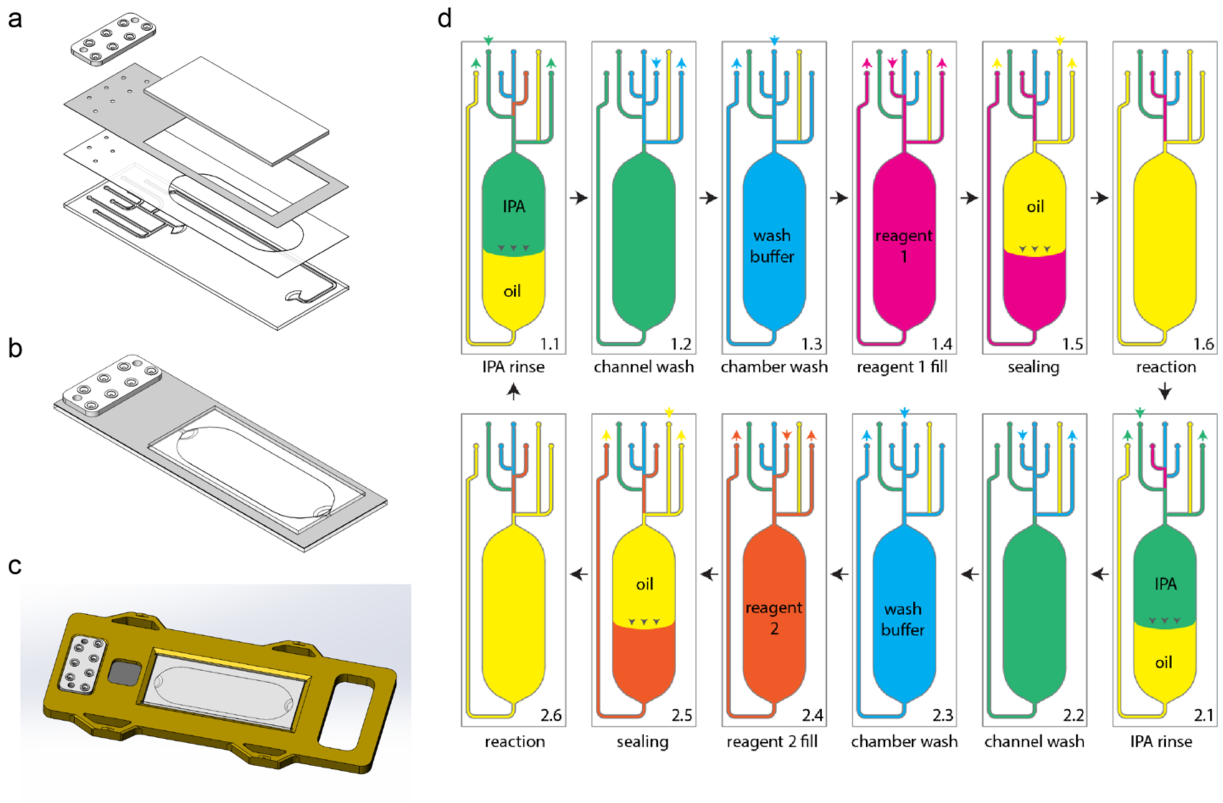


**Figure S4.** Chip. (a) The schematic of the structure and fabrication of the flow-cell chip. (b) The fully assembled flow-cell chip. (c) The flow-cell chip and its holder. (d) The fluid flowing process in each reaction period that contains two reaction cycles.

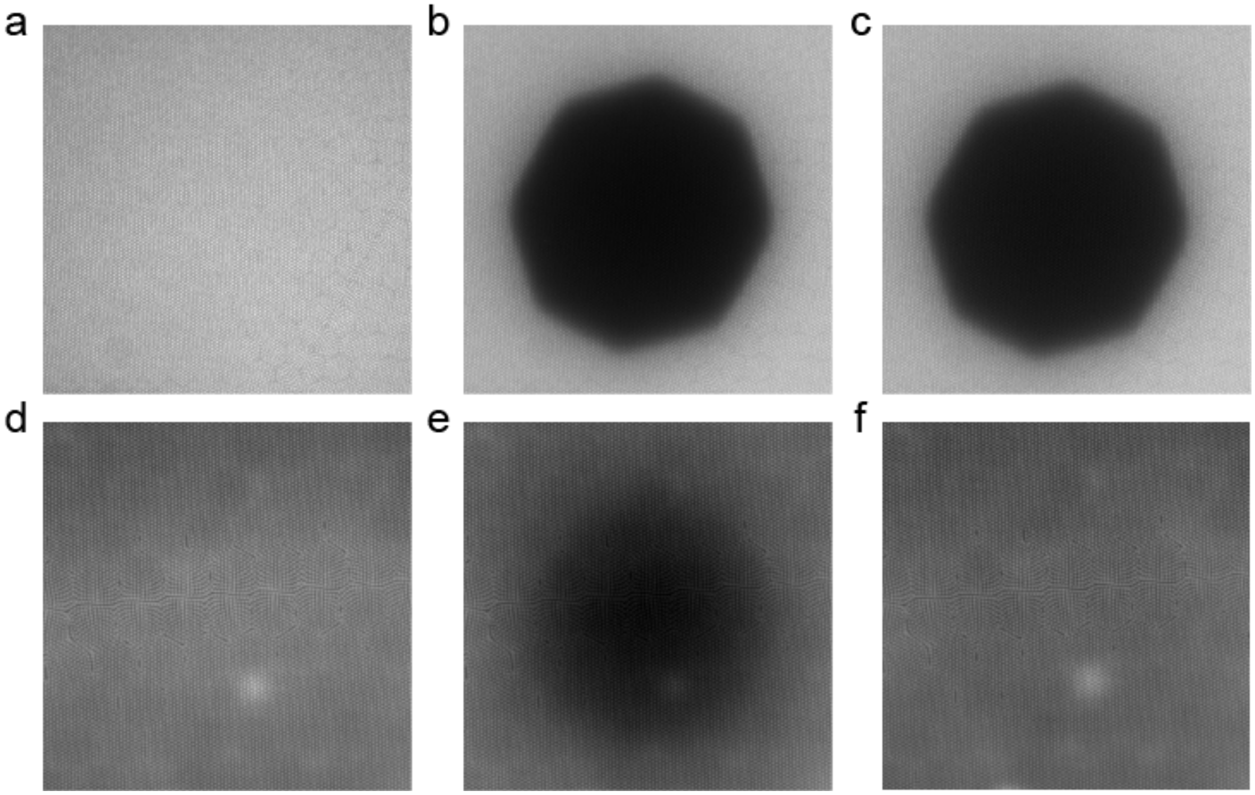


**Figure S5.** The bleaching test to confirm the effectiveness of sealing of the microreactors. (a) Field-of-view (FOV) where fluorophores were sealed inside the microwells by FC-40. (b) Using the pinhole to selectively photo-bleach part of the FOV, the fluorescent intensity is lower in the exposed area. (c) After several minutes, the margin of the exposed area remains sharp and clear, indicating the sealing is successful so that non-bleached fluorophores in the unexposed area cannot diffuse into the bleached area. (d-f) When sealing fails, the margin of exposed area is blurred, and the intensity of the exposed area recovers as a result of fluorophore diffusion.

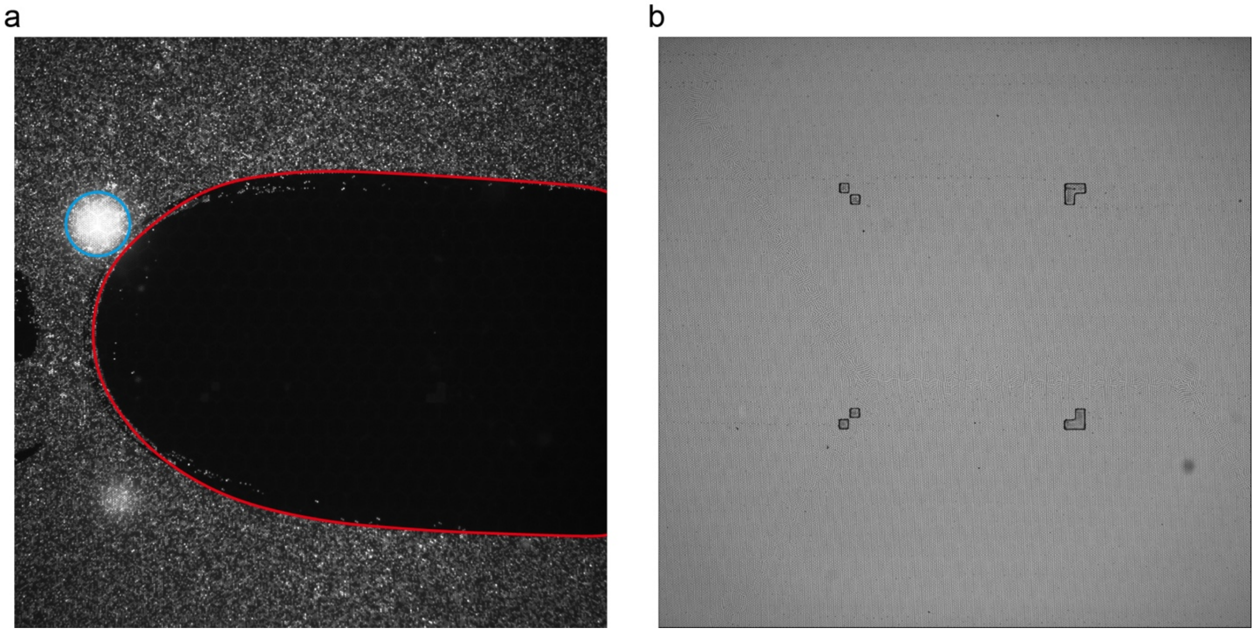


**Figure S6.** Image. (a) A typical example of light stains (blue) and dark stains (red). (b) A typical example of bright-field image with tetris-like positioning markers.

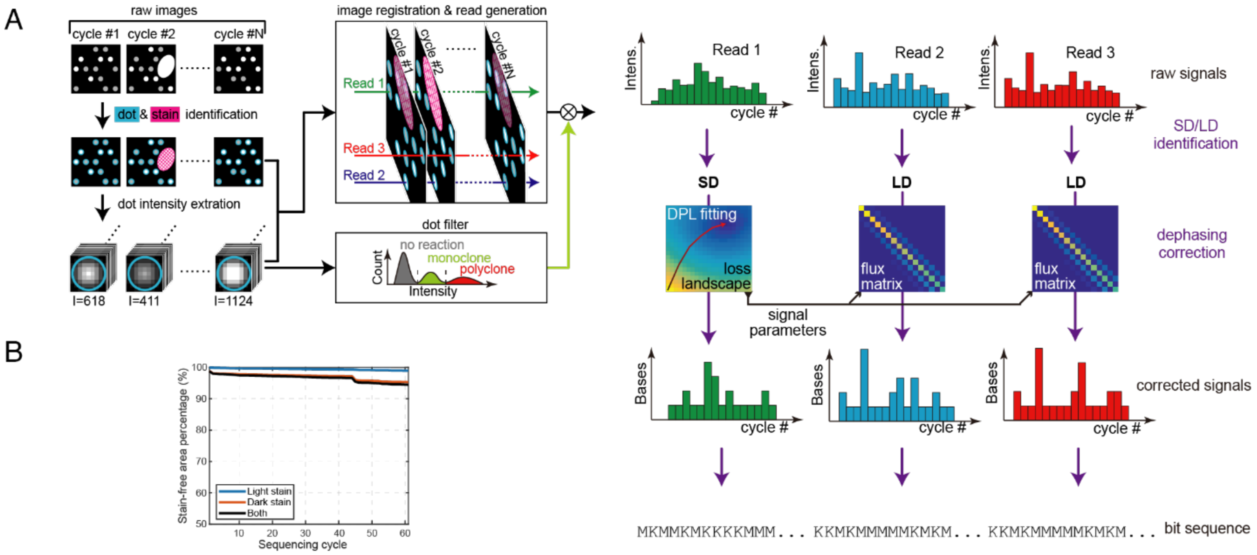


**Figure S7.** The data processing pipeline. (A) The schematic of the data processing pipeline. Defects are filtered and the intensity of each microwell at each reaction cycle is extracted and further filtered to keep monoclones for dephasing correction. The correction parameters are determined by fitting the signal to the standard dots (SD) which have been spiked into the sequencing library.(B) >90% of the area remains stain-free after 61 cycles of sequencing.

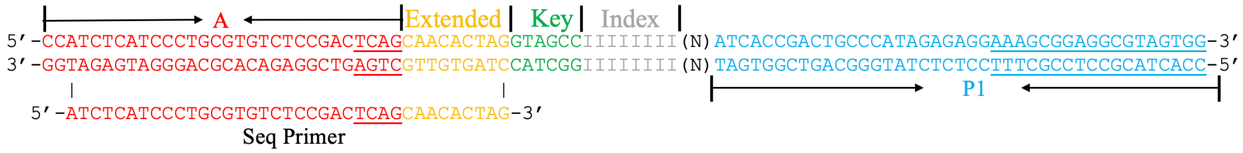


**Figure S8.** The structure of the DNA library. The 8-bp index sequence is written as I's and the inserted part is written as N. The library is modified from ion torrent's library by adding an extended sequence to enhance the melting temperature of the sequencing primer and a key sequence for LD identification. The underlined TCAG is used in ion torrent for library identification and signal normalization.

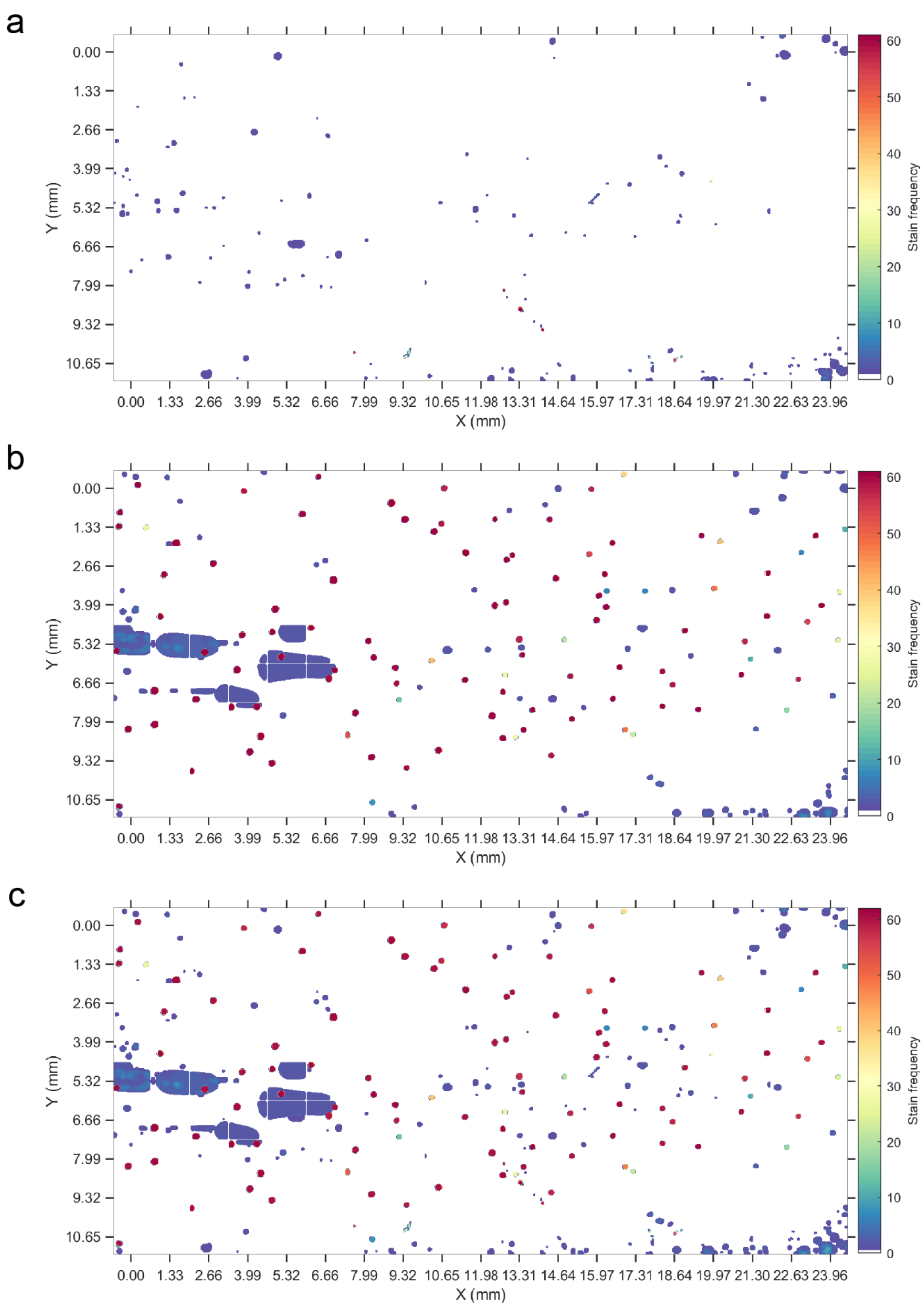


**Figure S9.** A typical result of the spatial distribution of light stains (a), dark stains (b) and both (c) in one sequencing experiment, the analysis shows the stains that observed during a 61-cycle sequencing run. The beads sitting within these stains are excluded from further analysis.

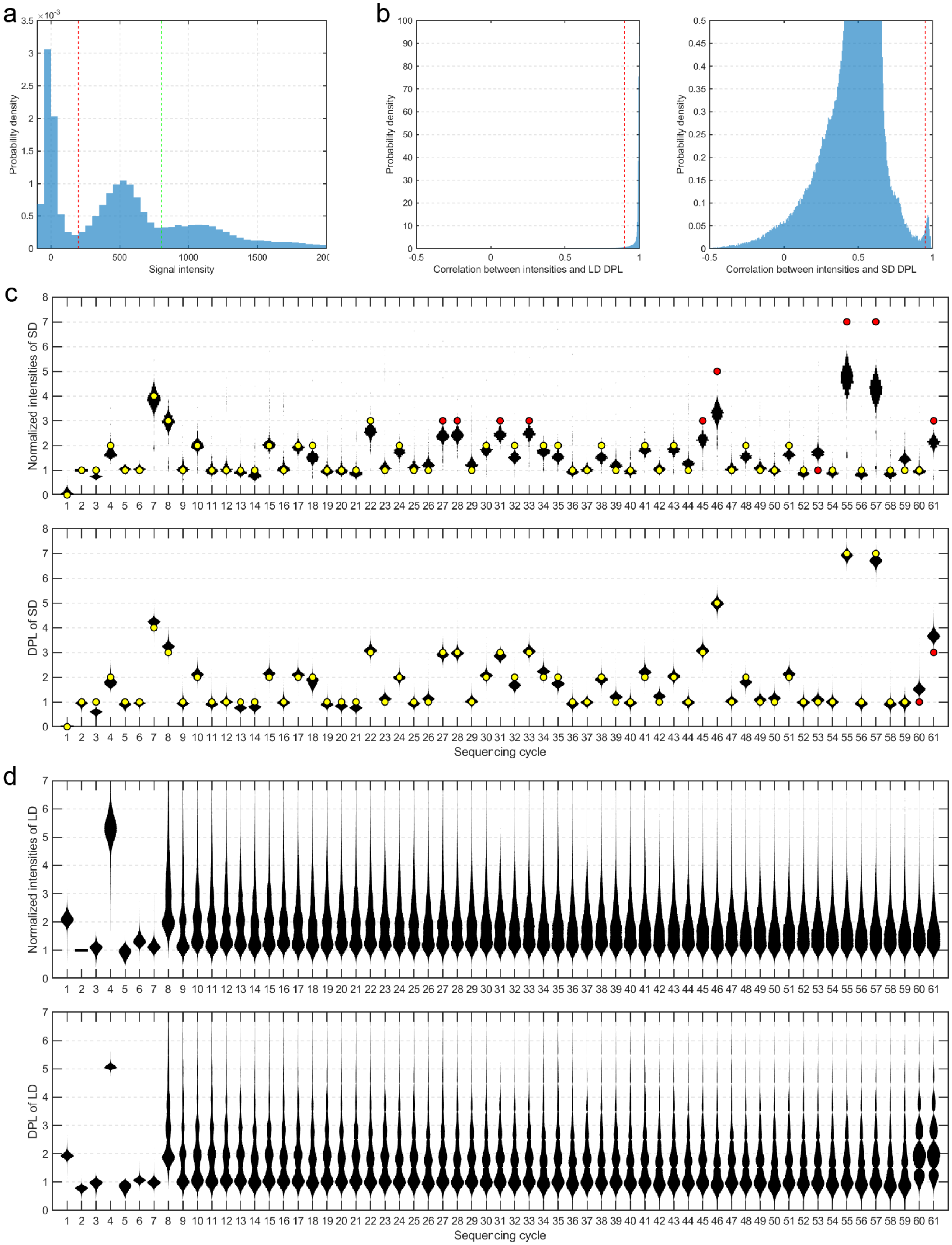


**Figure S10.** Filter pipelines in the fuzzy sequencing. (a) Schematic of how dots were selected according to their intensities. There were roughly three peaks in the distribution of dot intensities in each cycle. The troughs were found as the thresholds (the red and green vertical dash). Dots with intensities lower than the red dash were deemed as dark dots, and those greater than the green dash were deemed as polyclone. (b) LD and SD were identified through the Pearson's correlation coefficient ρ between the raw intensities of each dot and the starting DPL of LD/SD. We used $\rho > 0.9$ and $\rho > 0.95$ for LD and SD, respectively, as indicated by the red vertical dash. (c) Intensity distribution of SD in each sequencing cycle. Top: intensities normalized by dividing by the intensities of Cyc 2. Bottom: The DPLs corrected from dephasing. Black: intensity distributions. Yellow dots: Ideal DPLs equal to the rounded mean intensity. Red dots: Ideal DPLs not equal to the rounded mean intensity. The SD all had the same sequence and were shown as single peaks in all cycles. (d) Intensity distribution of LD in each sequencing cycle. Top: intensities normalized by dividing by the intensities of Cyc 2. Bottom: The DPLs corrected from dephasing. The first 7 cycles of LD were from the key region, showing as a single peak. The latter cycles of LD were from the inserted random region, showing as multiple peaks. The peaks of LD merged gradually due to the dephasing effect (top panel) but were re-separated by the dephasing correction (bottom panel).

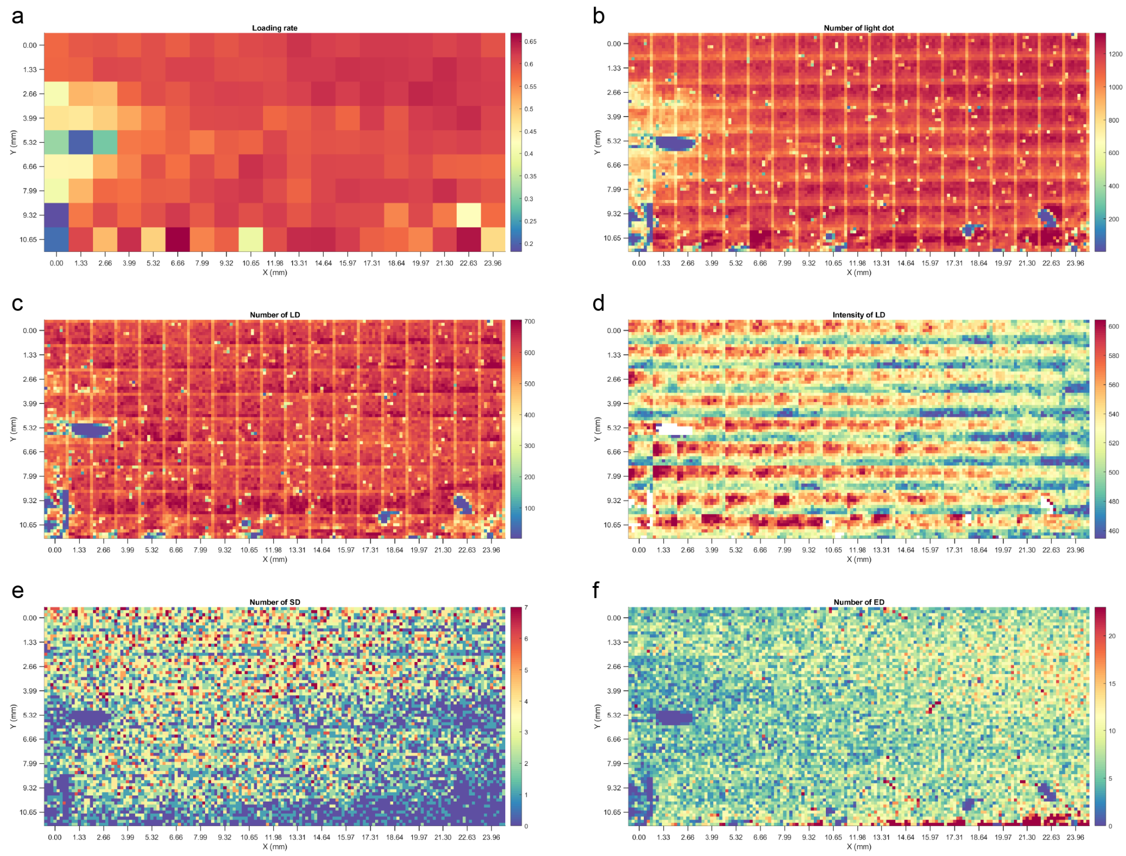


**Figure S11.** Dot number. (a) The spatial distribution of loading rate (the percentage of the wells containing beads) per tile. (b) The spatial distribution of number of light dot number. (c) The spatial distribution of LD number. (d) The spatial distribution of unit fluorogenic signal intensity of LD dots. (e) The spatial distribution of SD number. (f) The spatial distribution of ED number. In b-f, numbers were counted in 8×8 regions that each tile was divided into.

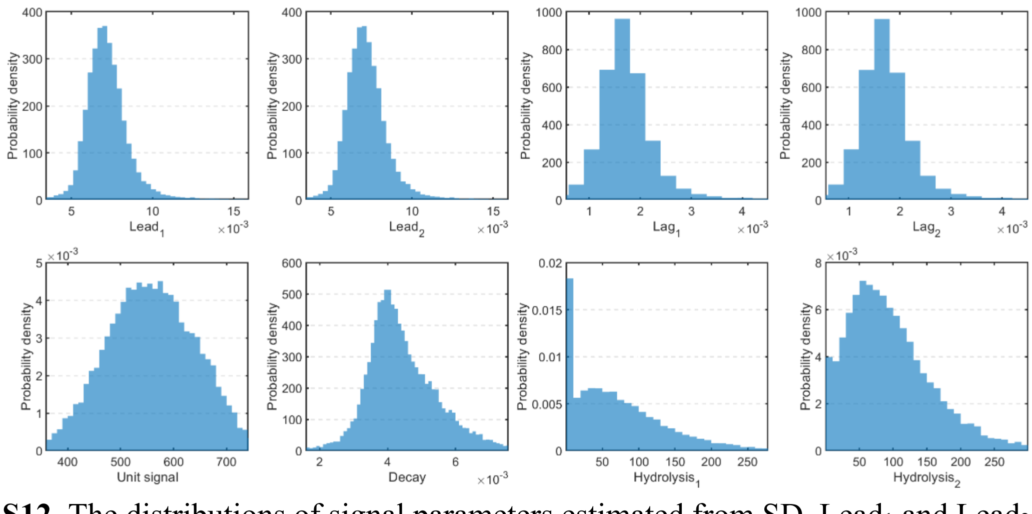


**Figure S12.** The distributions of signal parameters estimated from SD. $Lead_1$ and $Lead_2$ are almost identical, and $Lag_1$ and $Lag_2$ are almost identical due to the very large regularization term.

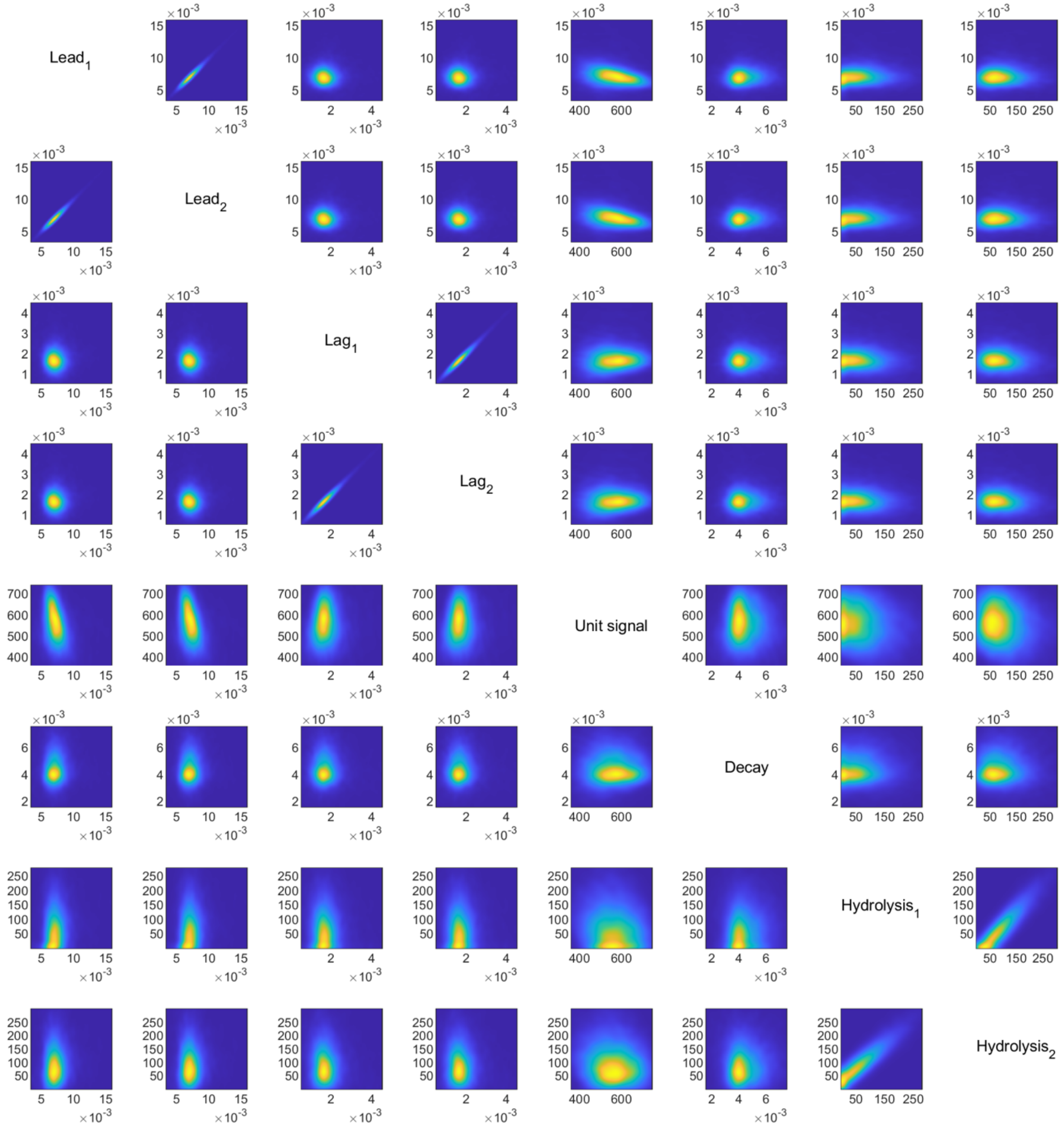


**Figure S13.** The correlation between signal parameters estimated from SD. $Lead_1$ and $lead_2$ are almost identical, and $lag_1$ and $lag_2$ are almost identical due to the very large regularization term.

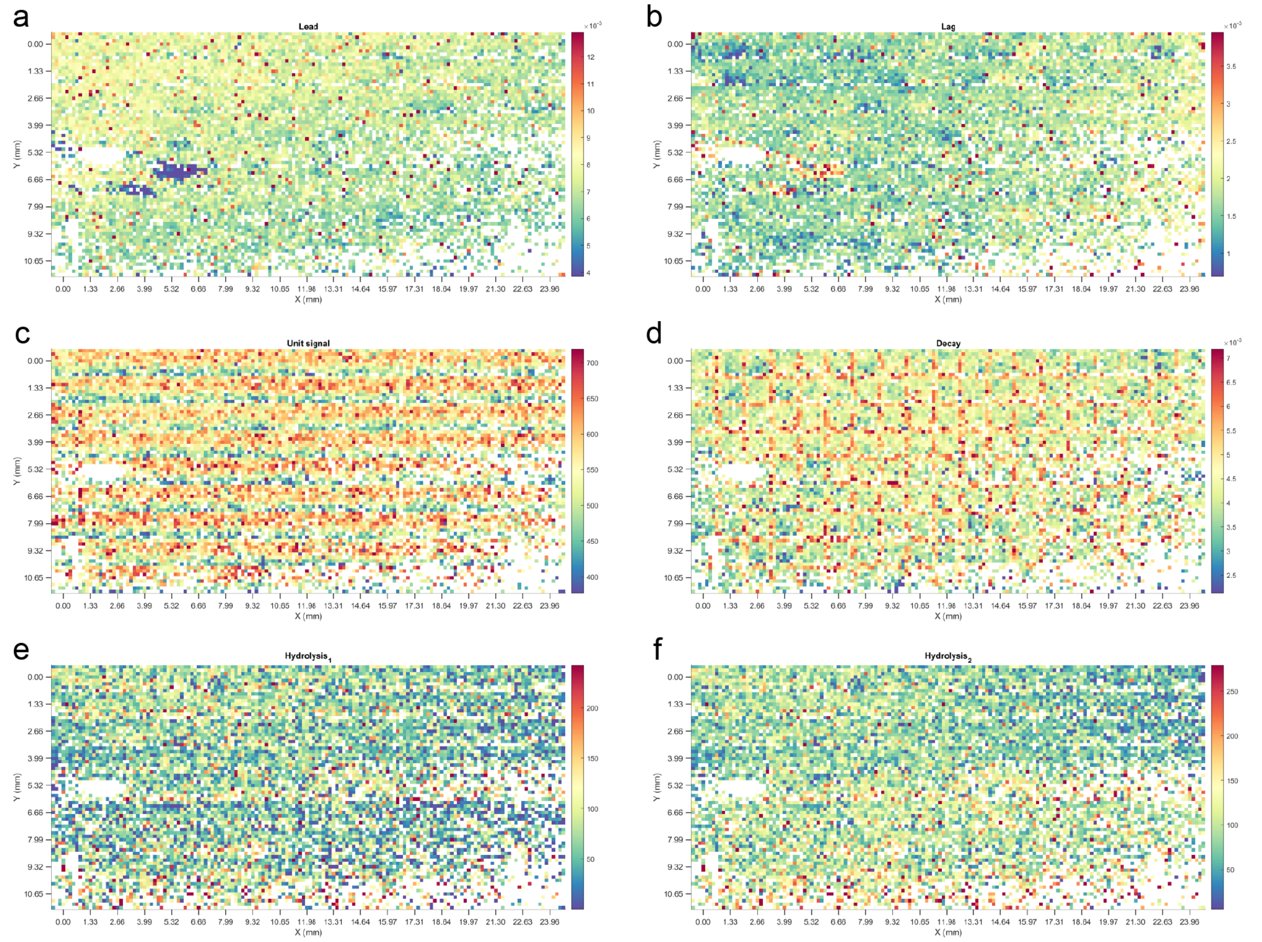


**Figure S14.** The spatial distribution of lead (a), lag (b), unit fluorogenic signal (c), decay (d), hydrolysis$_1$ (e) and hydrolysis$_2$ parameter (f) estimated from SD. Lead$_1$ and lead$_2$, lag$_1$ and lag$_2$ are almost identical due to the very large regularization term. We divided each tile into 8×8 regions and calculated the mean parameter value of dots in each region.

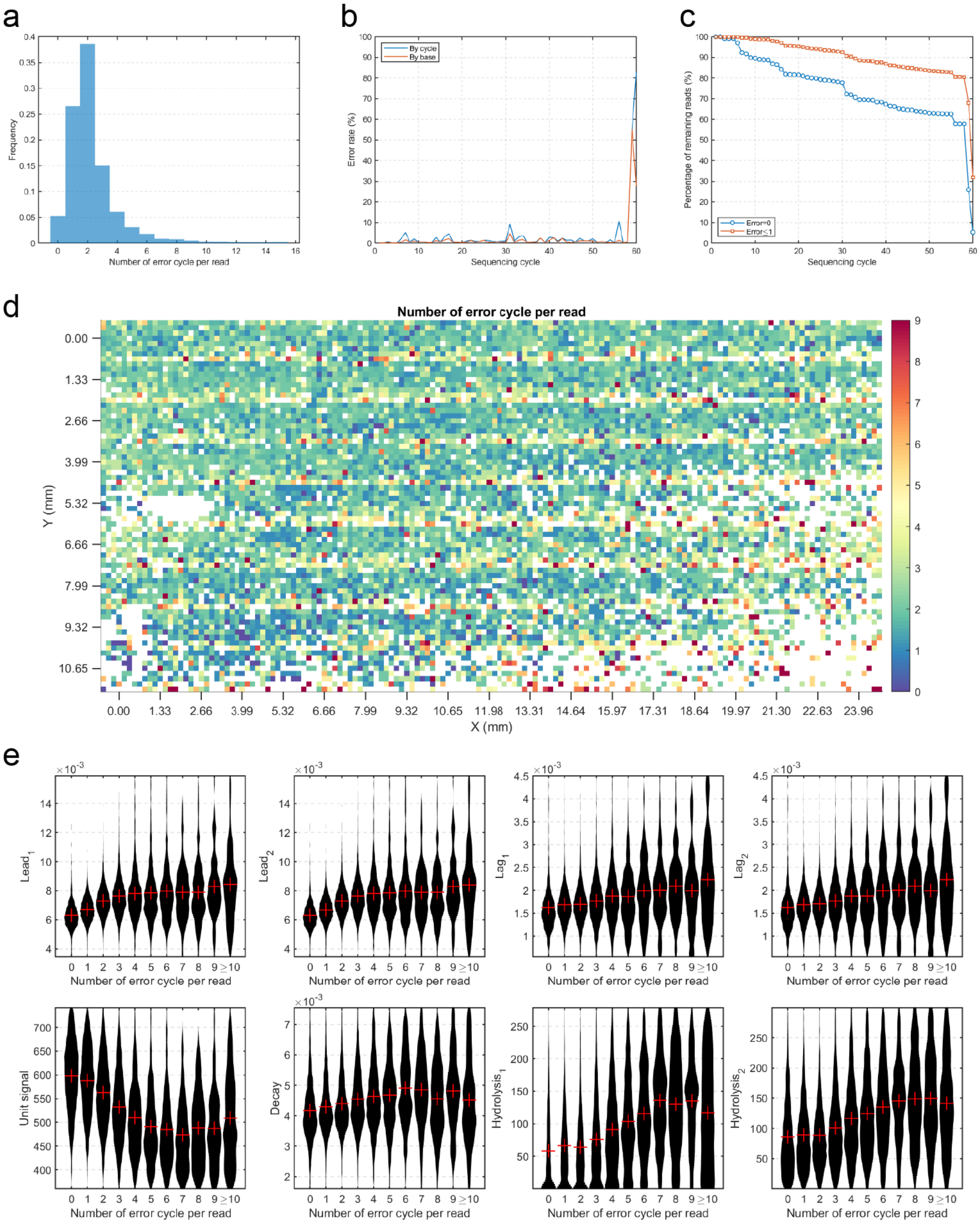


**Figure S15.** SD error. (a) The distribution of number of errors per read in SD. (b) Error rate of SD. The error rate by base is the error rate by cycle divided by the corresponding degenerate polymer length (DPL). (c) The percentage of reads with zero or one error in the first n cycles. The two lines both drop greatly in the last two cycles, indicating that most errors occur in the last two reaction cycles. (d) The spatial distribution of number of errors per read in SD. Numbers were counted in 8×8 regions that each tile was divided into. (e) Relationship between signal parameters and error number per read (SD).

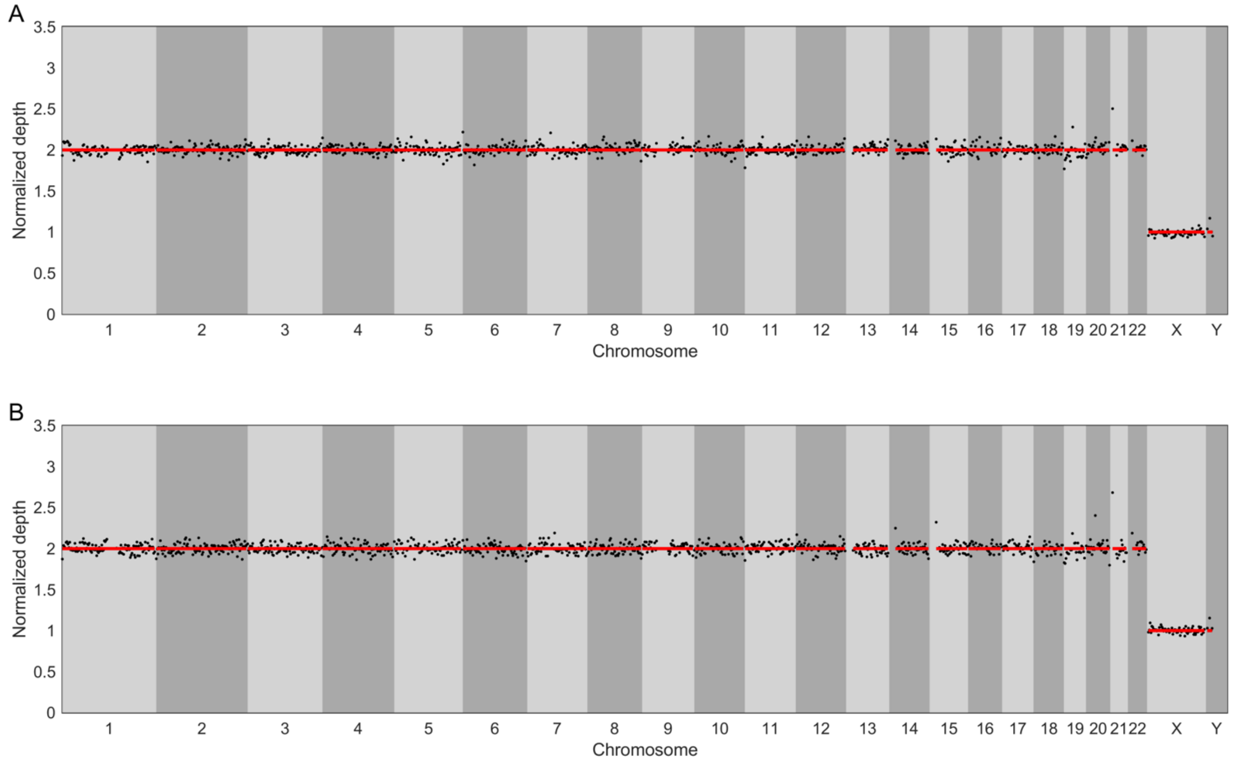


**Figure S16.** The normalized sequencing depth of a normal male adult. Top: BitSeq. Bottom: Ion Torrent.

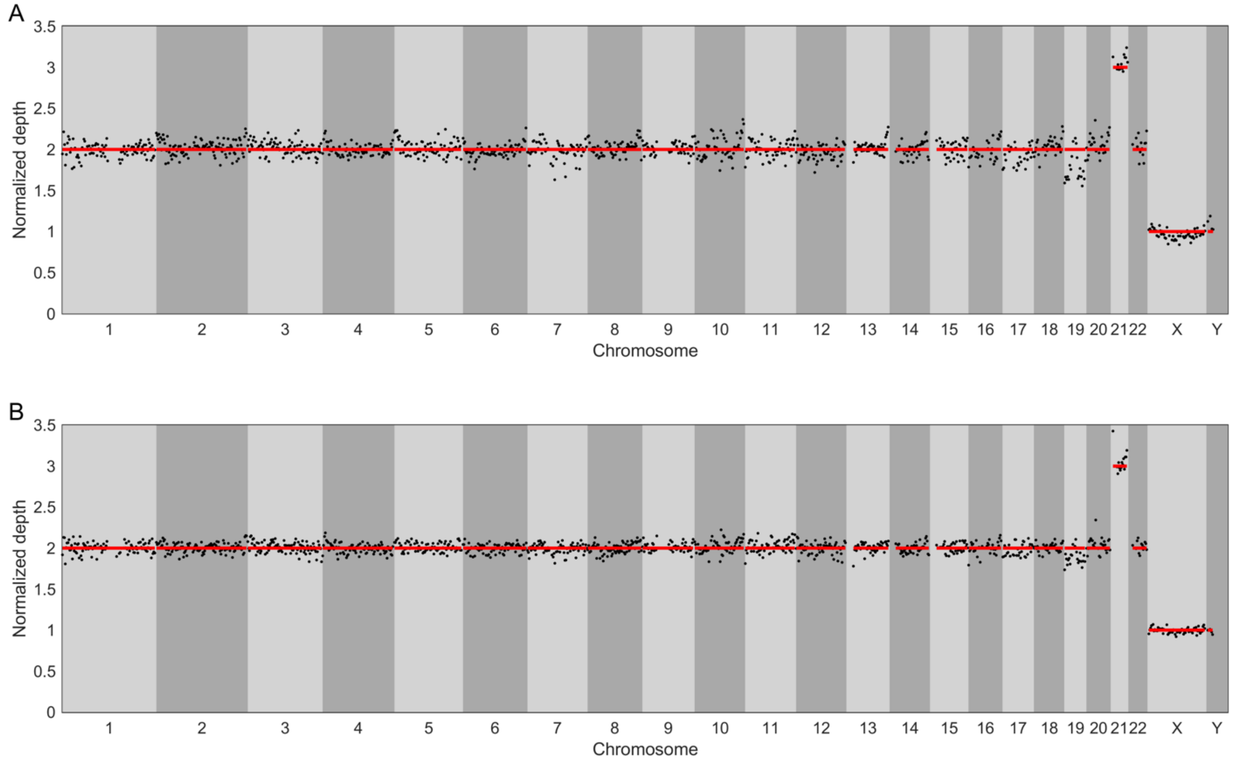


**Figure S17.** The normalized sequencing depth of a Trisomy 21 patient. Top: BitSeq. Bottom: Ion Torrent.

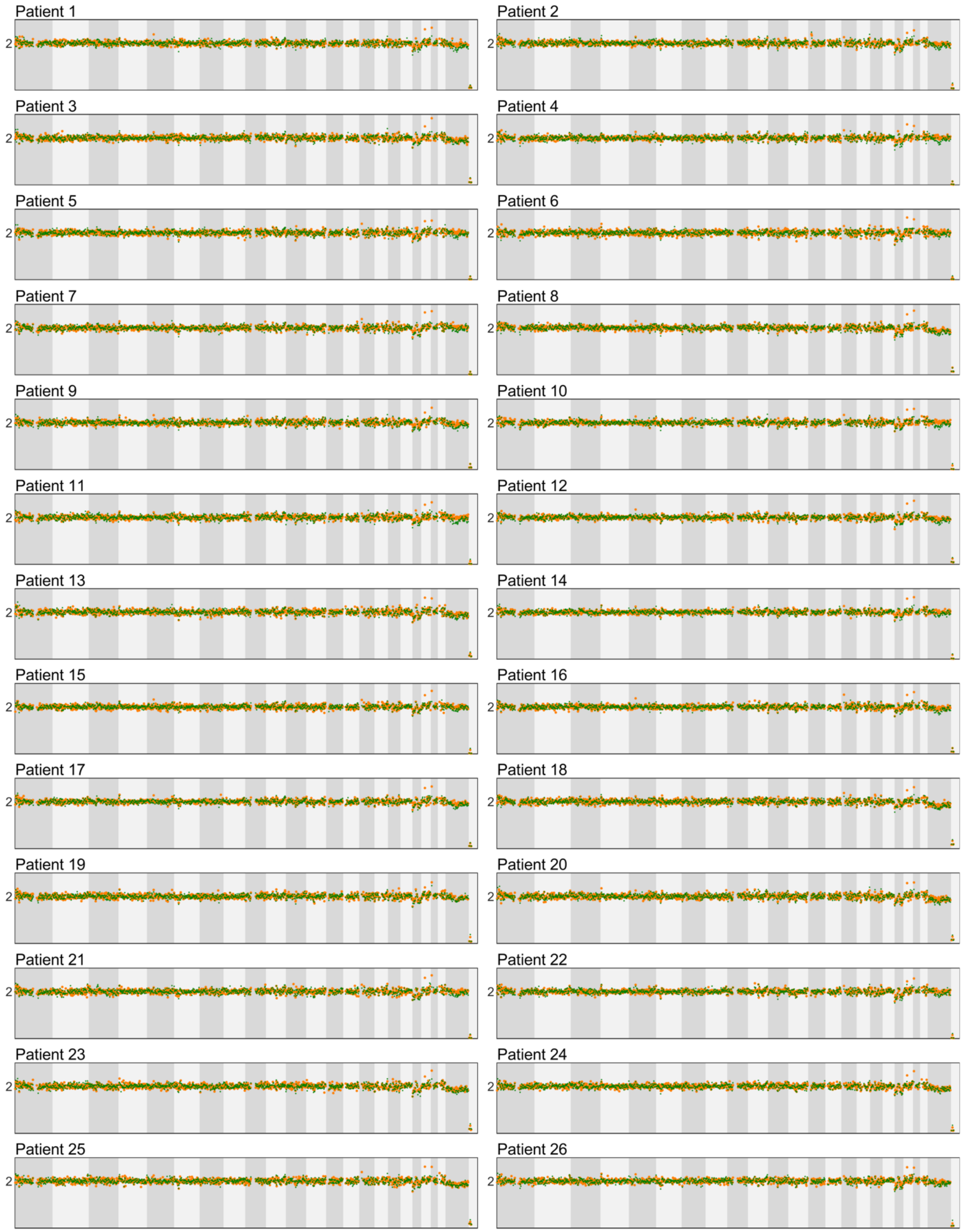
Patient 1
2
Patient 2
2
Patient 3
2
Patient 4
2
Patient 5
2
Patient 6
2
Patient 7
2
Patient 8
2
Patient 9
2
Patient 10
2
Patient 11
2
Patient 12
2
Patient 13
2
Patient 14
2
Patient 15
2
Patient 16
2
Patient 17
2
Patient 18
2
Patient 19
2
Patient 20
2
Patient 21
2
Patient 22
2
Patient 23
2
Patient 24
2
Patient 25
2
Patient 26
2

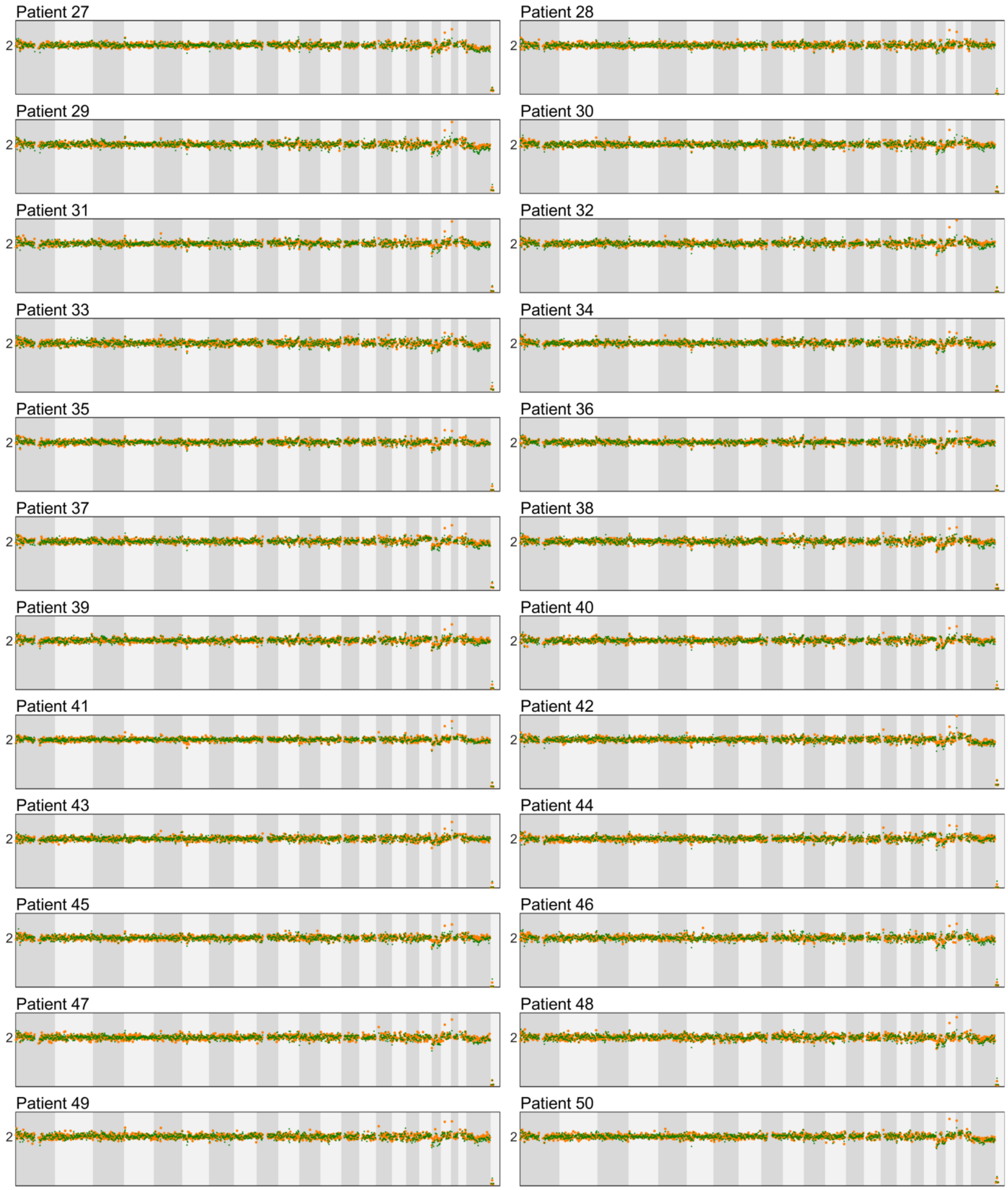


**Figure S18.** The normalized sequencing depth of 53 patients involved in the NIPT study. X axis: Genome sites. Green dots: BitSeq. Orange dots: Ion Torrent. The dark and light gray backgrounds indicate the chromosomes.

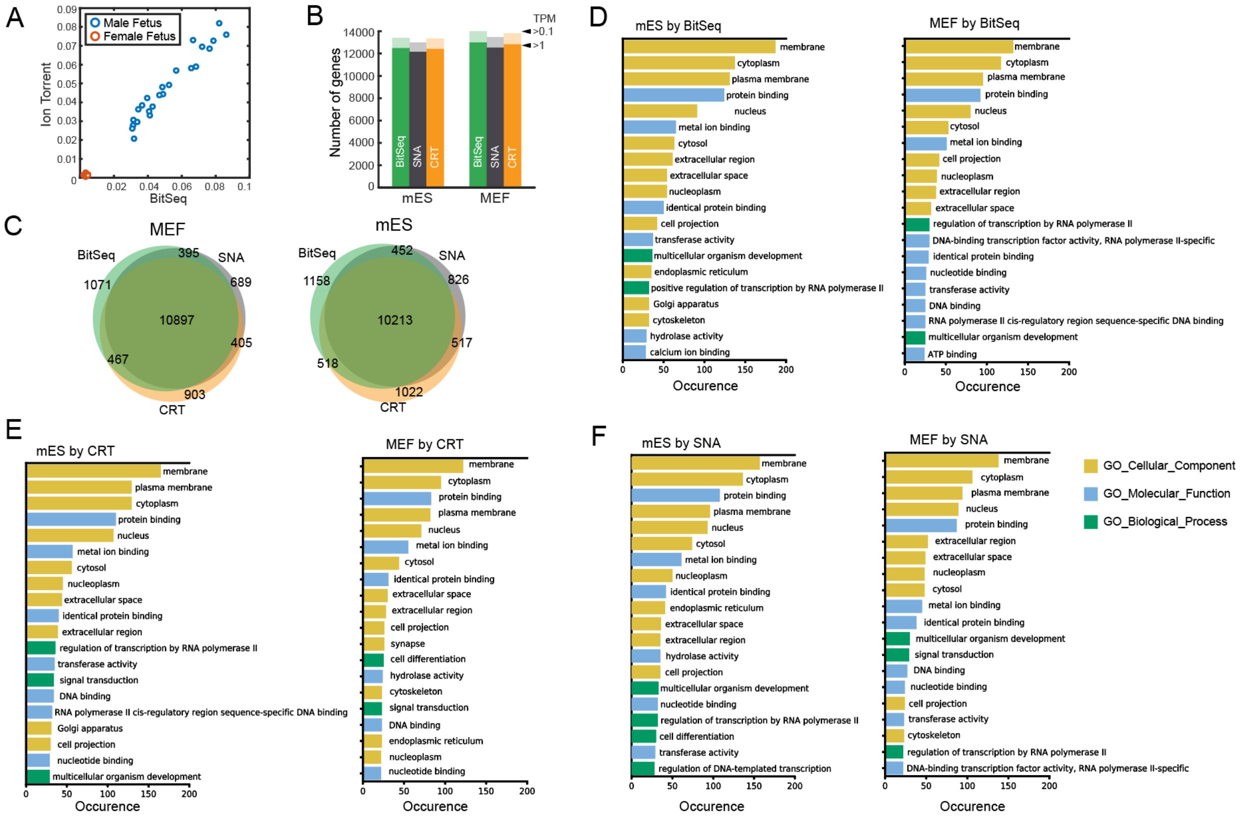


**Figure S19.** Additional comparison of BitSeq with commercial sequencers in resequencing. (A) The average sequencing depth of ChrY of each sample. These values by both BitSeq and ion torrent differ significantly between male and female fetuses. (B) Number of genes detected using different thresholds for RNA-seq data of mES and MEF cells. (C) Venn diagram of TPM>1 genes. (D-F) Gene Ontology analysis of genes that were only detected by each sequencer in RNA-seq.

**BS in MK**

```
Template: 5'-GACAATGCTACCTTACCGGTCGGAACTCGA-3'
   1 cycle : 0
   2 cycles: 01111
   3 cycles: 0111100
   5 cycles: 011110010
  10 cycles: 0111100101110011110001
```

**BS in RY**

```
Template: 5'-GACAATGCTACCTTACCGGTCGGAACTCGA-3'
   1 cycle : 11
   2 cycles: 110
   3 cycles: 11011
   5 cycles: 1101101
  10 cycles: 11011010010000100
```

**CRT**

```
Template: 5'-GACAATGCTACCTTACCGGTCGGAACTCGA-3'
   1 cycle : G
   2 cycles: GA
   3 cycles: GAC
   5 cycles: GACAA
  10 cycles: GACAATGCTA
```

**Figure S20.** How error-free sequences were simulated. BS: BitSeq. CRT: cyclic reversible terminator (the sequencing chemistry used by Illumina and MGI sequencers).

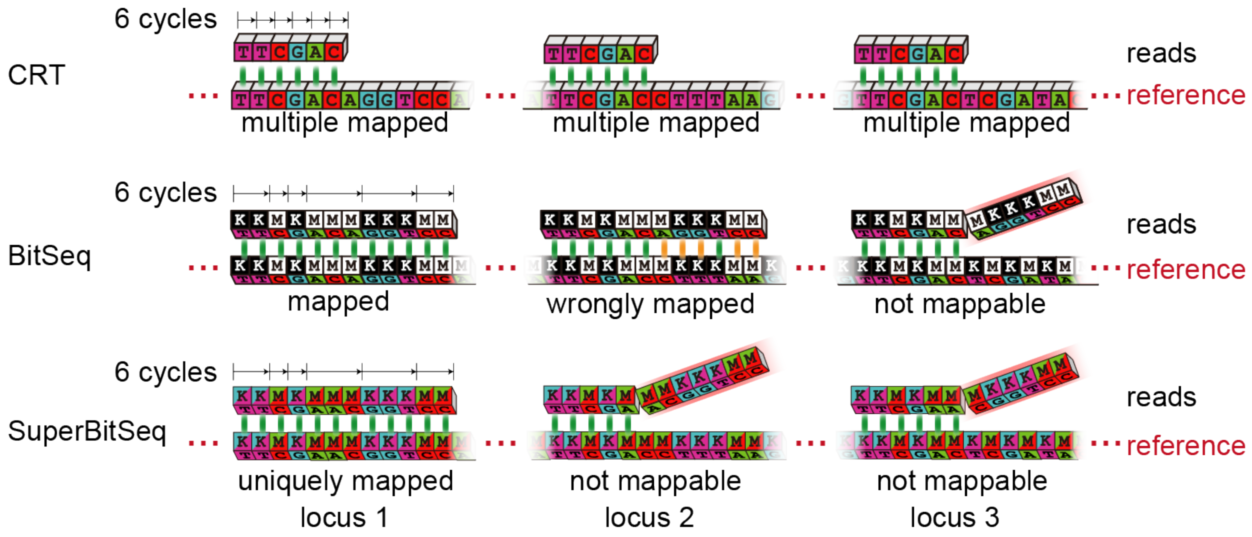


**Figure S21.** Longer read length or more information enable more accurate genome mapping.

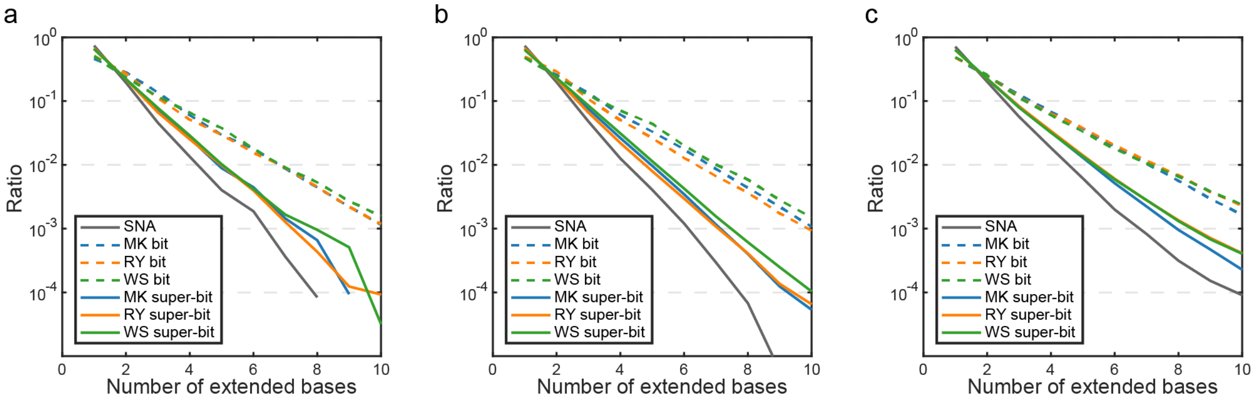


**Figure S22.** Polymer distributions of genomes of lambda phage (a), *Escherichia coli* (b) and *Saccharomyces cerevisiae* (c).

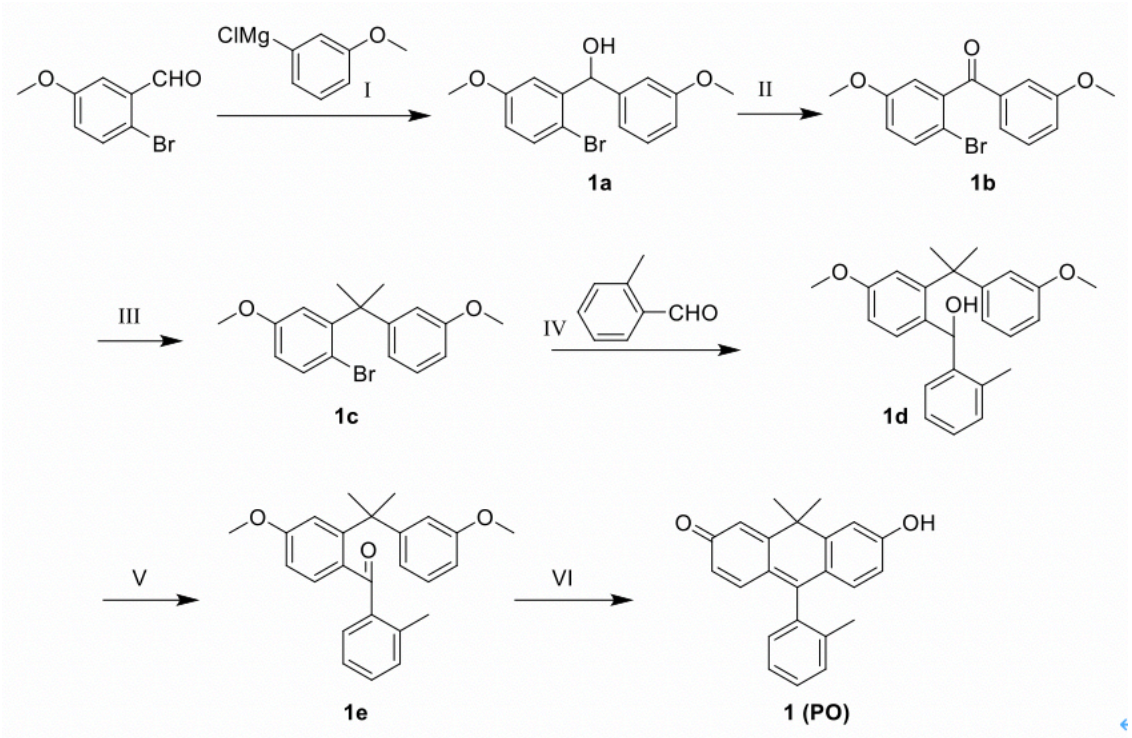


**Figure S23.** The synthetic route of a newly developed fluorophore ‘Peking Orange’ (PO).

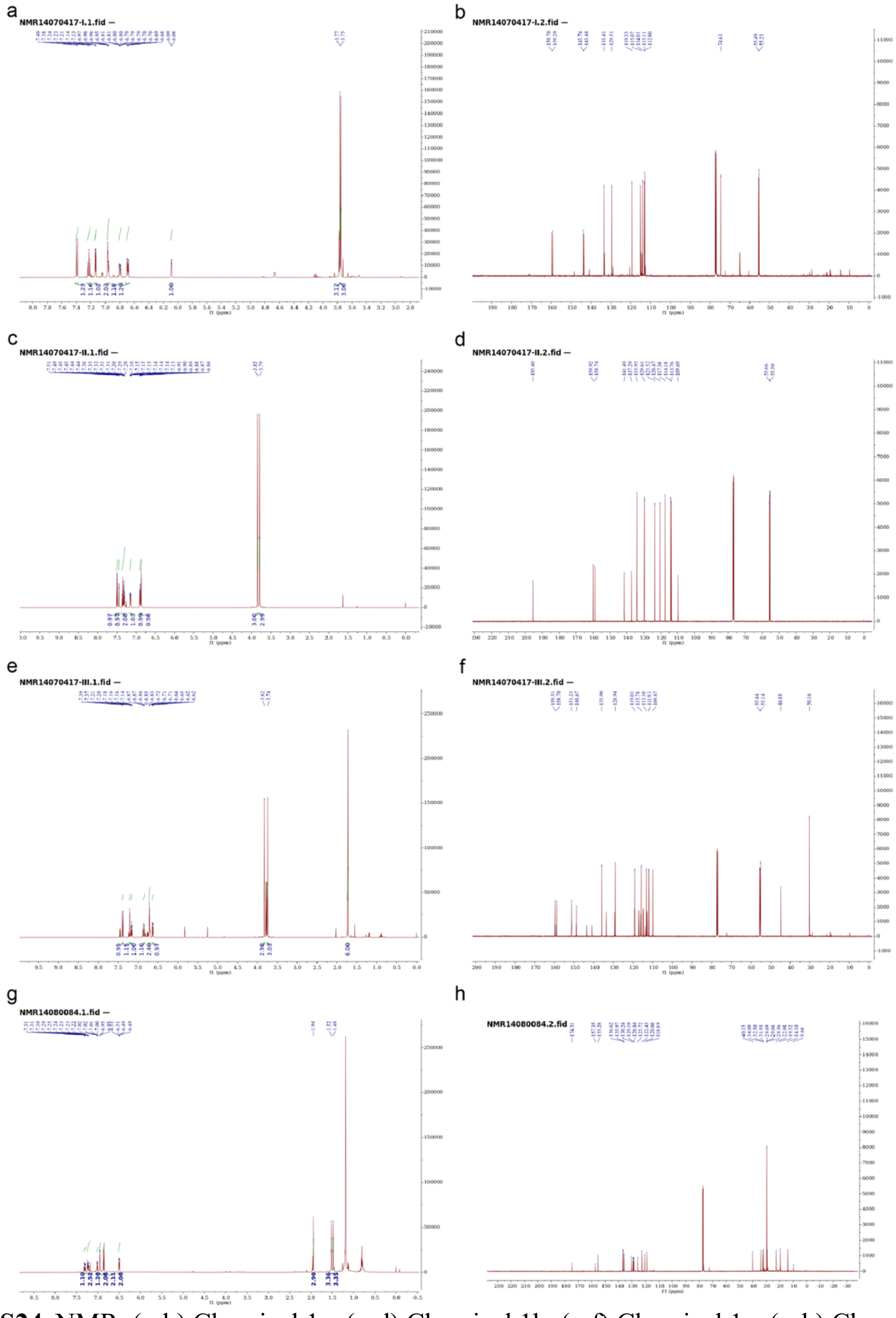


**Figure S24.** NMR. (a-b) Chemical 1a. (c-d) Chemical 1b. (e-f) Chemical 1c. (g-h) Chemical PO. (aceg) $^{1}$H-NMR. (bdfh) $^{13}$C-NMR.

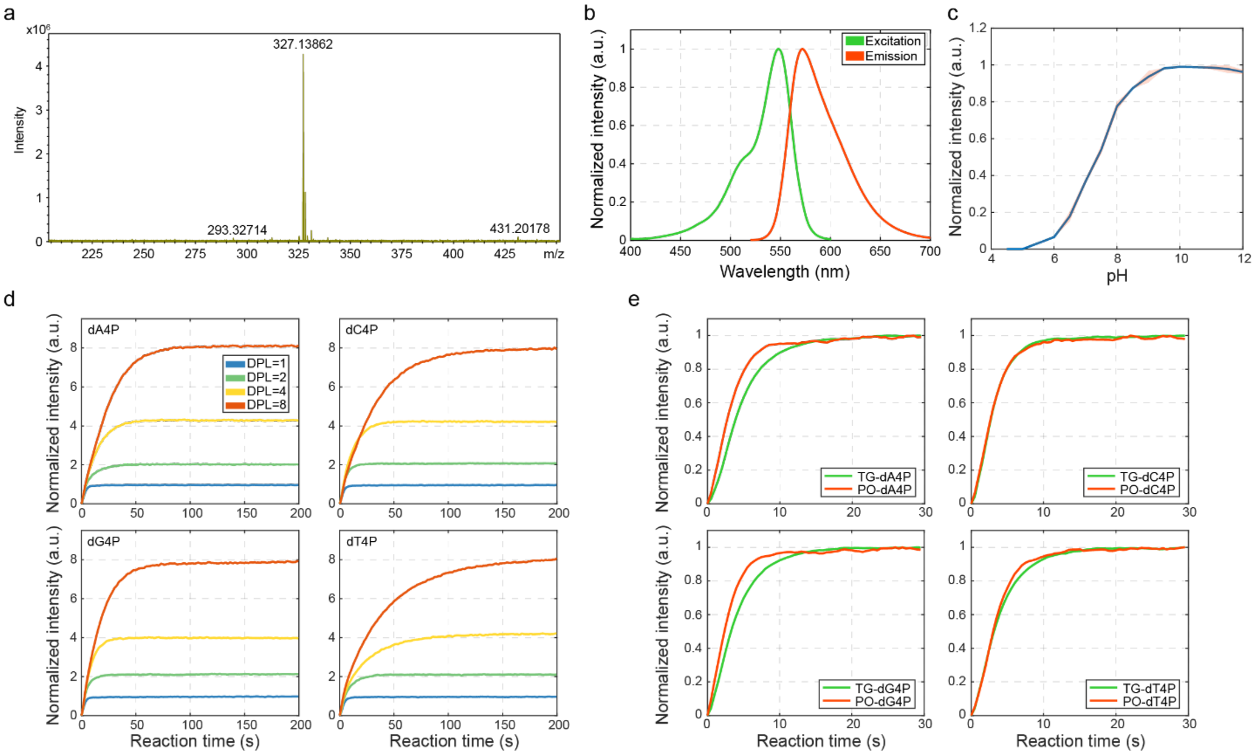


**Figure S25.** PO properties. (a) HRMS of PO. (b) Excitation and emission spectra of PO. (c) The fluorescence emission of PO is pH sensitive. The pink shade indicates the standard deviation. (d) Fluorogenic kinetics of PO-dN4P extending multiple bases. (e) The sequencing reaction kinetics using PO-dN4P and TG-dN4P as substrates.

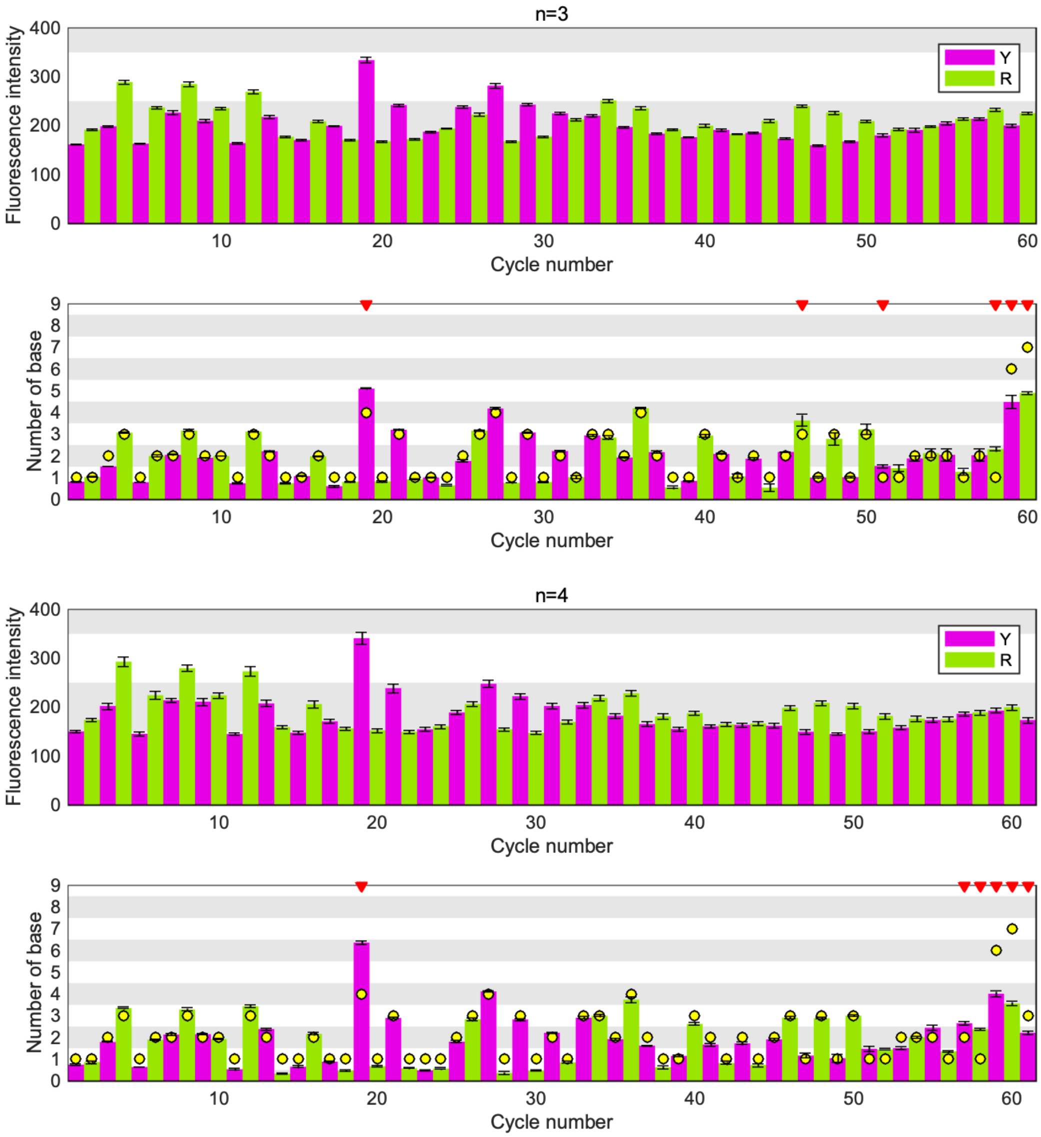
n=3
Fluorescence intensity
Cycle number
Y
R
Number of base
n=4

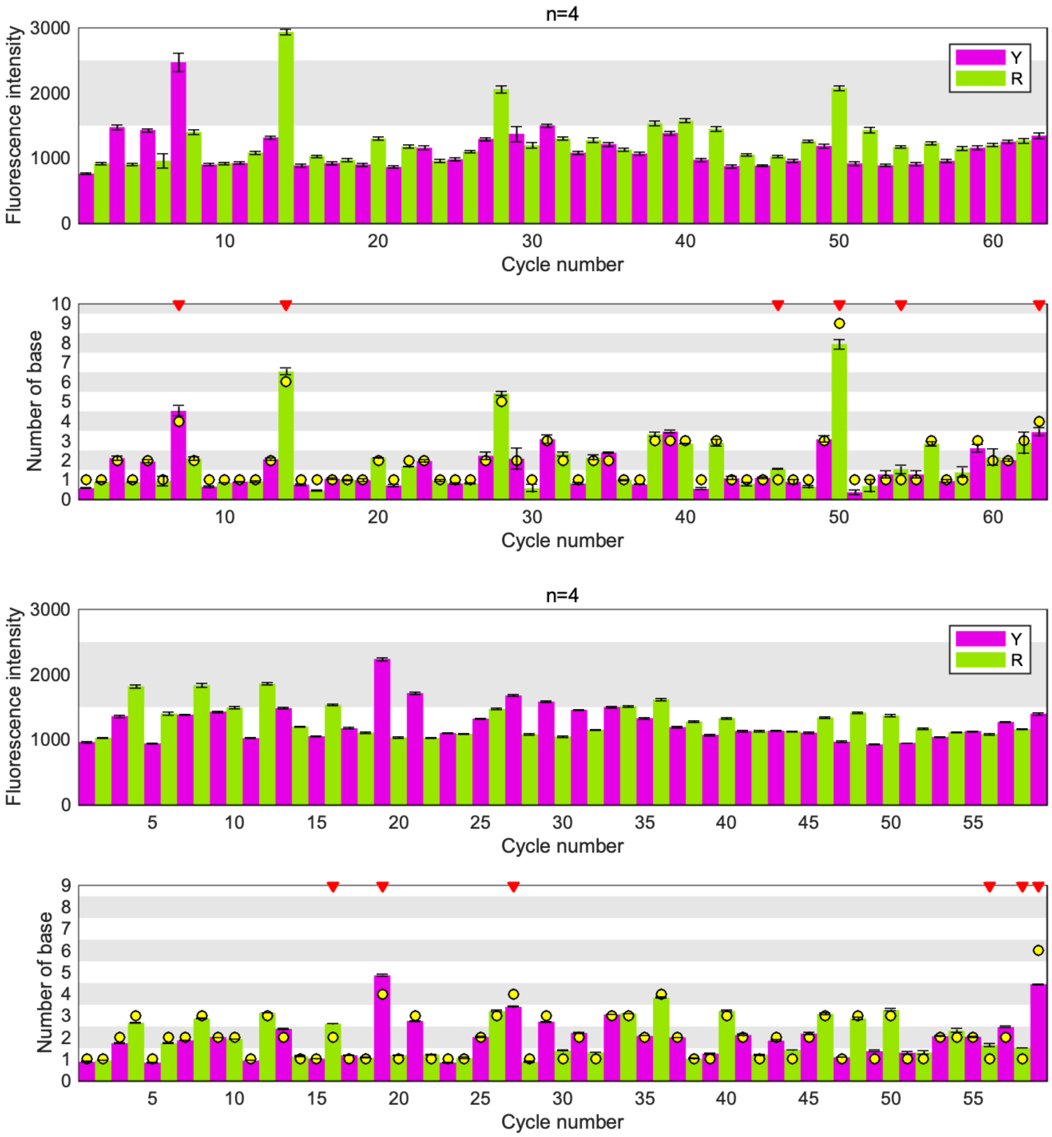


**Figure S26.** The fluorescence intensities (top) and their dephasing corrected signals (bottom) of a single-template BitSeq experiment using PO.

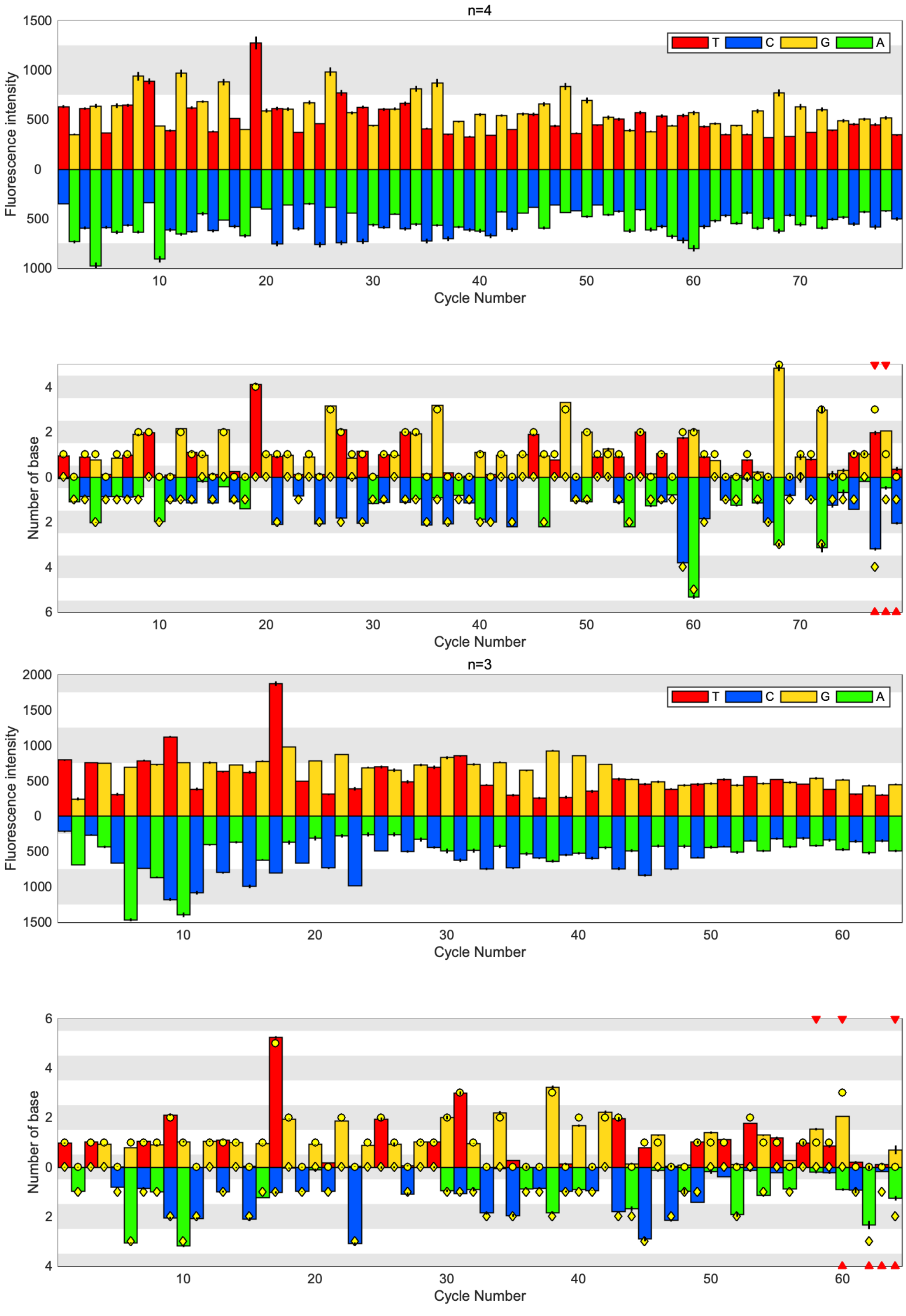


**Figure S27.** The fluorescence intensities (top) and their dephasing corrected signals (bottom) of a single-template SuperBitSeq experiment.

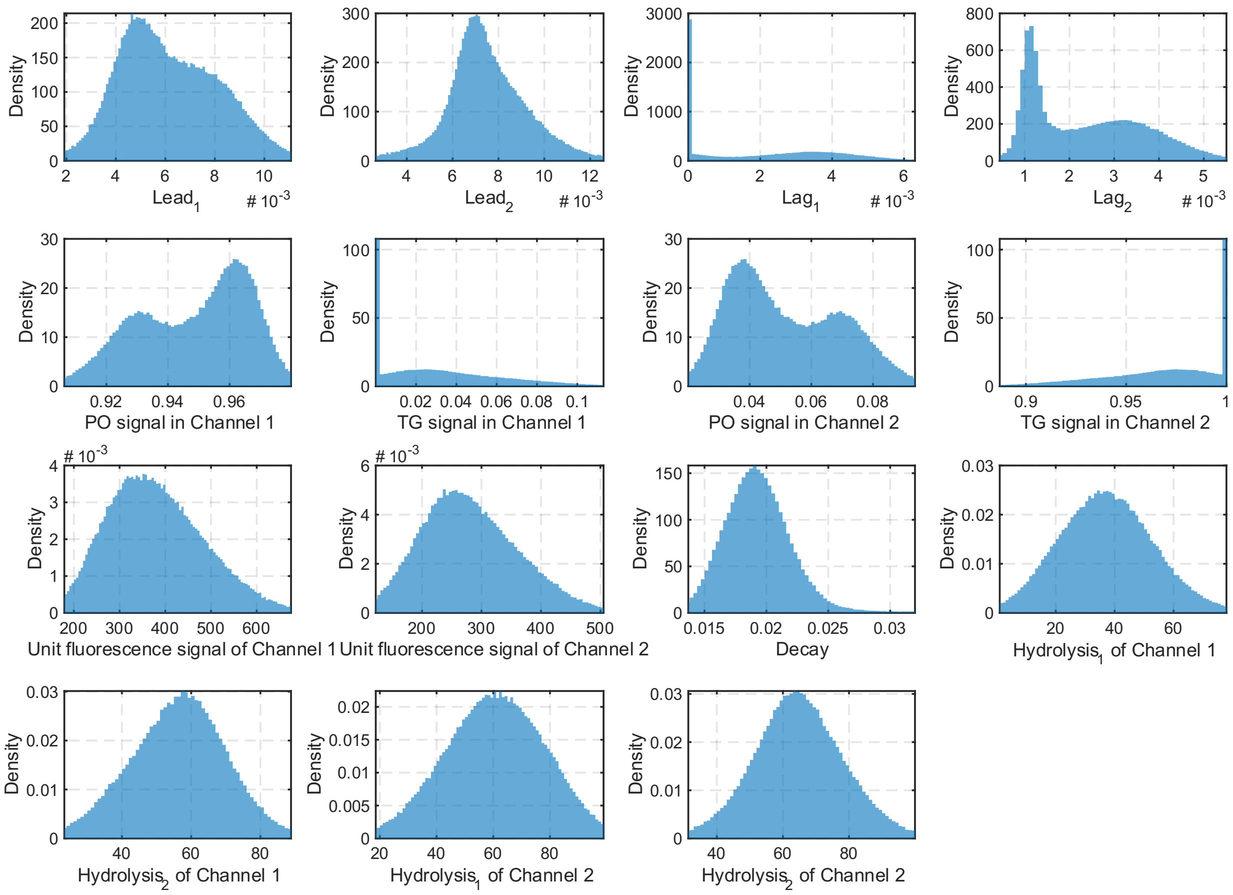


**Figure S28.** Parameter distributions of dephasing correction in SuperBitSeq.

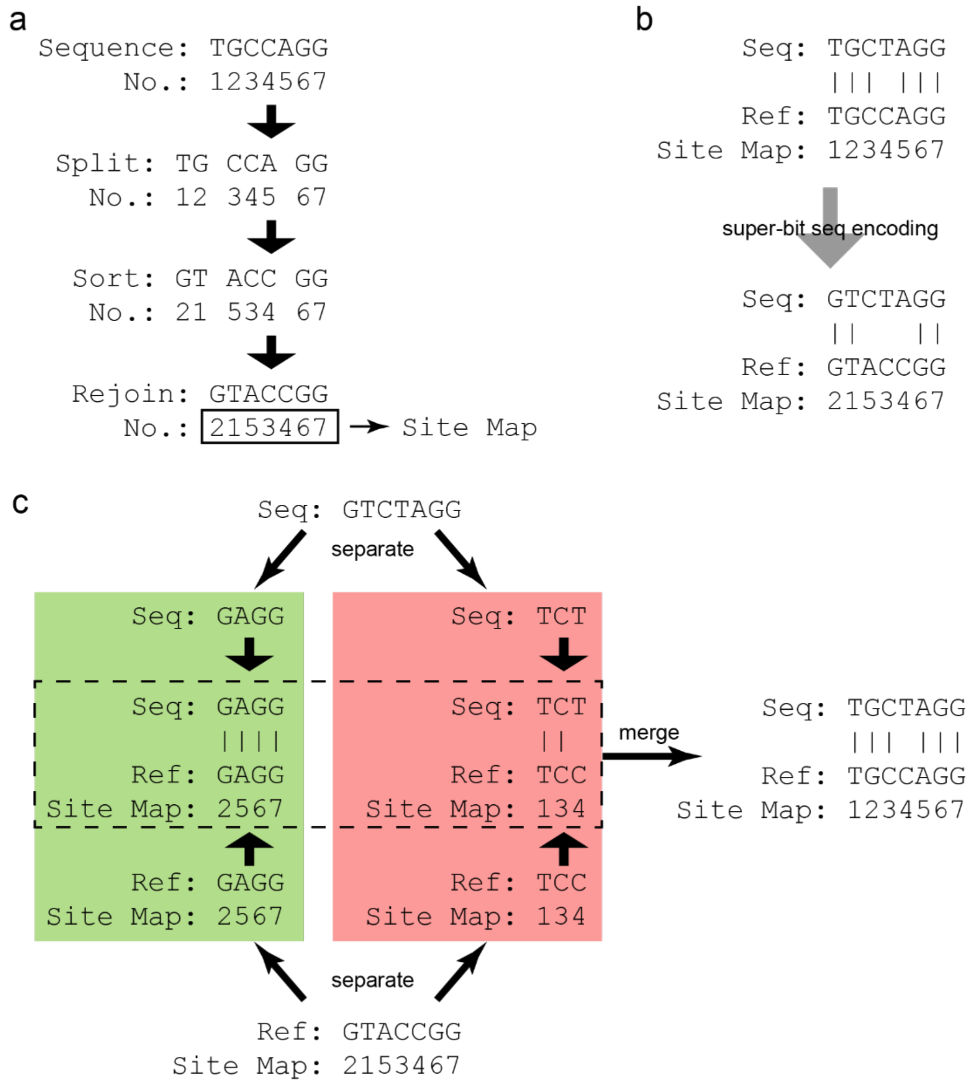


**Figure S29.** SNV reconstruction in SuperBitSeq. (a) Site map is defined as the positions of encoded bases in the original genome and is recorded during SuperBitSeq encoding. (b) In some cases SNV information may be altered after SuperBitSeq encoding. (c) Schematic of SNV reconstruction. The sequencing read and its matched reference sequence are separated into two semi-sequences, respectively. One semi-sequence is composed of A and G, which are both labeled with the green fluorophore Tokyo Green. The other semi-sequence is composed of C and T, which are both labeled with the red fluorophore Peking Orange. The semi-sequences are locally aligned individually, then the two alignments are merged together by sorting the site map in ascending order.

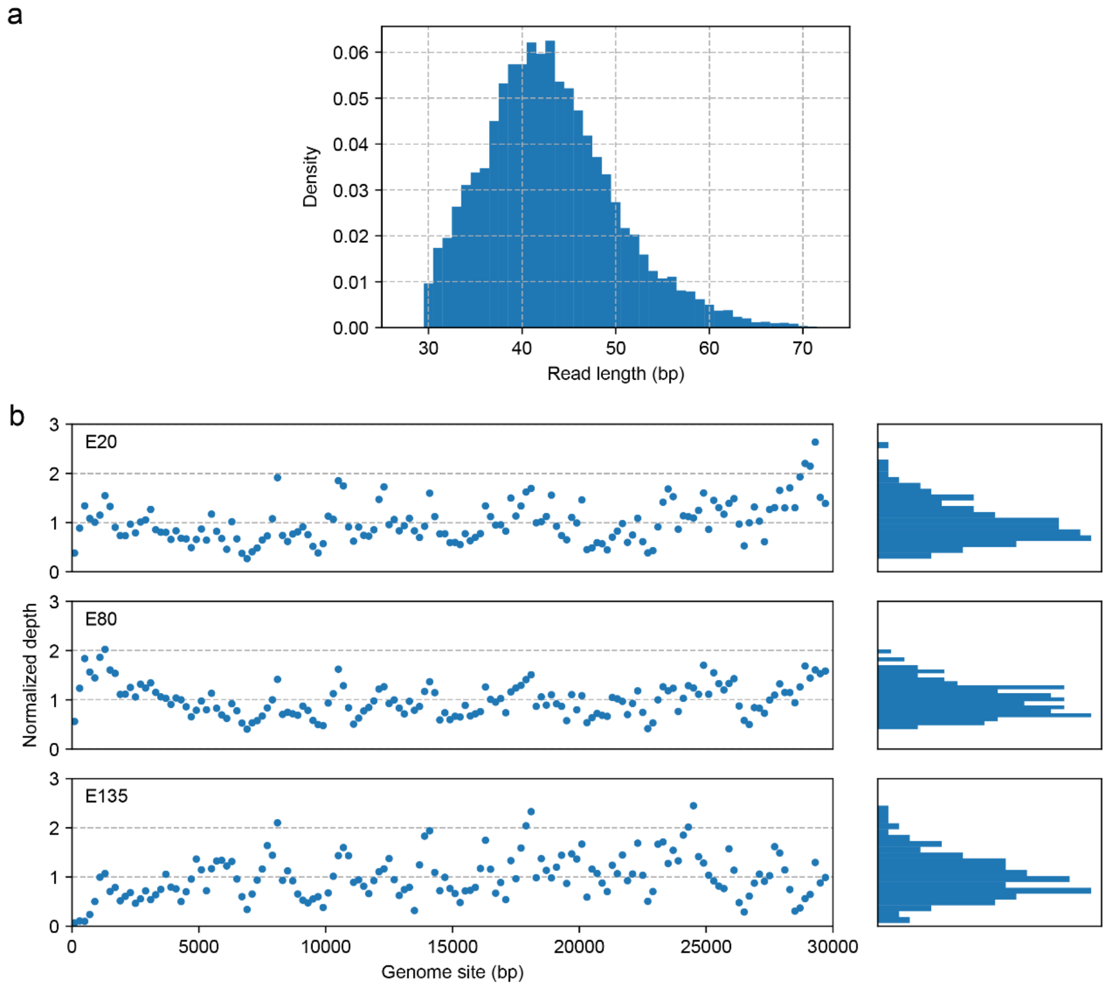


**Figure S30.** SuperBitSeq of SARS-CoV-2. (a) The read length distribution of superbit sequences. (b) Normalized depth of the 3 SARS-CoV-2 samples. Bin size: 200 bp.

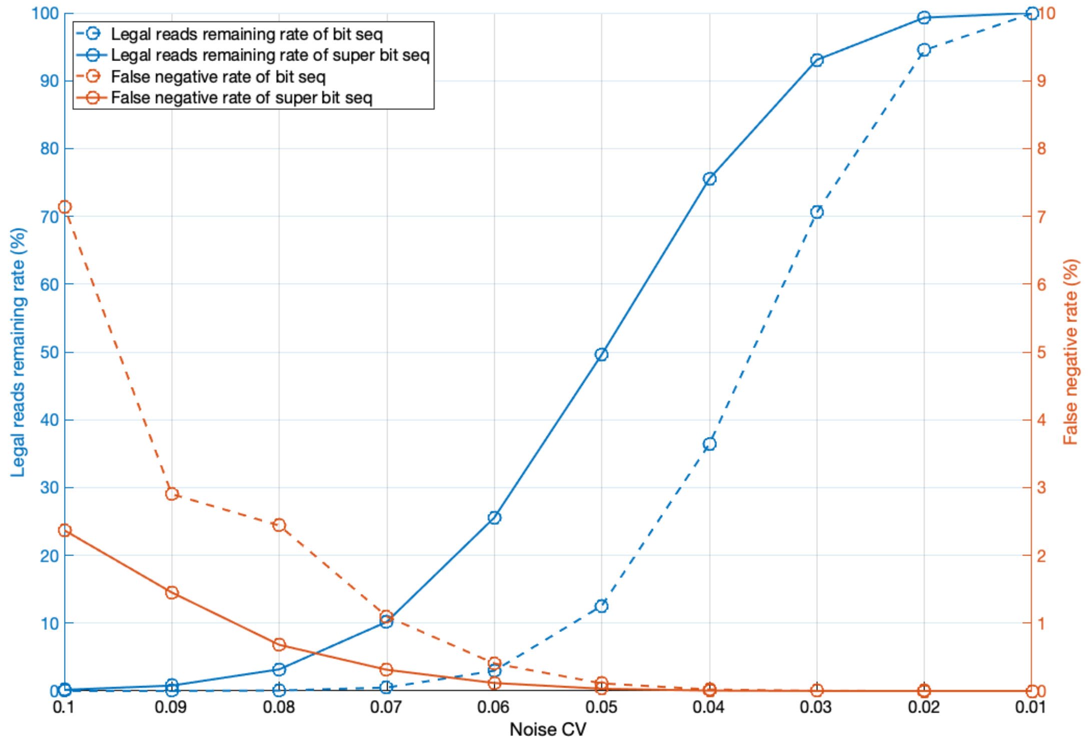


**Figure S31.** Error-correction capability of three rounds of bit sequences or superbit sequences. One round bit sequence or superbit sequence encode 50% or 84% information of the 4-base DNA sequence, and three orthogonal rounds (MK, RY, WS) of them will provide information redundancy that can be used to correct potential sequencing errors. To compare their error-correction capability, we did the following simulation:

1. Simulate $1 \times 10^7$ 100-bp random DNA sequence reads (denoted as $\boldsymbol{A}$);
2. Calculate their theoretical three-round sequencing signals $\boldsymbol{S}$;
3. Add white noise with standard deviation $\boldsymbol{\sigma} = \boldsymbol{CV} \times \boldsymbol{S} + \mathbf{0.05}$ to $\boldsymbol{S}$, then round to integers $\boldsymbol{D}$; The 0.05 in $\boldsymbol{\sigma}$ is to disturb the zero signals in SuperBitSeq;
4. Use the error-correction principle to check potential sequencing errors in $\boldsymbol{D}$;
5. If no error is found, mark $\boldsymbol{D}$ as legal;
6. For legal signals, decode the signal as a new sequence read $\boldsymbol{B}$;
7. If $\boldsymbol{B}$ is not a substring of $\boldsymbol{A}$ starting from the beginning, mark $\boldsymbol{B}$ as false negative (sequencing error not detected);
8. Calculate the ratio of legal signals and false negative sequences of BitSeq and SuperBitSeq under different $\boldsymbol{CV}$.

Thanks to reduced dynamic range and greater information entropy, error-correction of SuperBitSeq has higher legal signal rates and lower false-negative rate than BitSeq under the same noise $\boldsymbol{CV}$.

**Table S1**. Reaction time cost comparison between the fuzzy sequencer and CRT using BitSeq and NovaSeq as examples.

| Item | Fuzzy Sequencer | CRT |
|---|---|---|
| Incorporation time per cycle (s) | 30 | 23.7 |
| Deblock time per cycle (s) | 0 | 16.4 |
| Total reaction time per cycle (s) | 30 | 40.1 |
| Cycle number to read 100 bp | 50 | 100 |
| Total reaction time to read 100bp (min) | 25 | 66.8 |

**Table S2.** Sequences of DNA templates used in sequencing. All sequences are written from 5’ to 3’. The inserted part, namely, those to be sequenced instead of library adaptors, is underlined. The mutation site is marked as red.

| Name | Sequence |
|---|---|
| SD | CCATCTCATCCCTGCGTGTCTCCGACTCAGCAACACTAGA<br>TCAGCTGTGACAGCAGAGCTGCGTAATCTCCCGCATATTG<br>CCAGCATGGCCTTTAATGAGCCGCTGATGCTTGAACCCGC<br>CTATGCGCGGGTTTTCTTTTGTGCGCTTGCAGGCCAGCTTG<br>GGATCAGCAGCCTGACGGATGCGGTGTCCGGCGACATCA<br>CCGACTGCCCATAGAGAGG |
| EGFR G719S wild type (2155G) | CCTTACCTTATACACCGTGCCGAACGCACCGGAGCCCAGC<br>AAGATCGGAAGAGCGTCGTGTAGGGAAAGAGTGT |
| EGFR G719S mutant (2155A) | CCTTACCTTATACACCGTGCCGAACGCACCGGAGCTCAGC<br>AAGATCGGAAGAGCGTCGTGTAGGGAAAGAGTGT |
| EGFR T790M wild type (2369C) | CGGACATAGTCCAGGAGGCAGCCGAAGGGCATGAGCTGC<br>GTGATGAGATCGGAAGAGCGTCGTGTAGGGAAAGAGTGT |
| EGFR T790M mutant (2369T) | CGGACATAGTCCAGGAGGCAGCCGAAGGGCATGAGCTGC<br>ATGATGAGATCGGAAGAGCGTCGTGTAGGGAAAGAGTGT |

**Table S3.** Different encoding strategies for the MK bit sequences. rc: reverse-complement.

| No. | Encoding | Operation to ACAGGTG |
|---|---|---|
| 1 | Encode every M to A and every K to T | $ACAGGTG \xrightarrow{encode} AAATTTT \xrightarrow{rc} AAAATTT$<br>$ACAGGTG \xrightarrow{rc} CACCTGT \xrightarrow{encode} AAAATTT$ |
| 2 | Encode every M to A and every K to C | $ACAGGTG \xrightarrow{encode} AAACCCC \xrightarrow{rc} GGGGTTT$<br>$ACAGGTG \xrightarrow{rc} CACCTGT \xrightarrow{encode} AAAACCC$ |

**Table S4.** Different encoding strategies for the WS bit sequences.

| No. | Encoding | Operation to ACAGGTG |
|---|---|---|
| 1 | Encode every W to A and every S to T | $ACAGGTG \xrightarrow{encode} ATATTAT \xrightarrow{rc} ATAATAT$<br>$ACAGGTG \xrightarrow{rc} CACCTGT \xrightarrow{encode} TATTATA$ |
| 2 | Encode every W to A and every S to C | $ACAGGTG \xrightarrow{encode} ACACCAC \xrightarrow{rc} GTGGTGT$<br>$ACAGGTG \xrightarrow{rc} CACCTGT \xrightarrow{encode} CACCACA$ |

| | | |
|---|---|---|
| 3 | Encode every W to AT and every S to CG | $ACAGGTG \xrightarrow{encode} ATCGATCGCGATCG \xrightarrow{rc}$ CGATCGCGATCGAT<br>$ACAGGTG \xrightarrow{rc} CACCTGT \xrightarrow{encode} CGATCGCGATCGAT$ |

**Table S5.** An example of SuperBitSeq encoding.

| Flow | Encoding process |
|---|---|
| MK | $AGATGC \xrightarrow{split} (A,G,A,TG,C) \xrightarrow{sort} (A,G,A,GT,C) \xrightarrow{rejoin} AGAGTC$ |
| RY | $AGATGC \xrightarrow{split} (AGA,T,G,C) \xrightarrow{sort} (AAG,T,G,C) \xrightarrow{rejoin} AAGTGC$ |
| WS | $AGATGC \xrightarrow{split} (A,G,AT,GC) \xrightarrow{sort} (A,G,AT,CG) \xrightarrow{rejoin} AGATCG$ |

**Table S6.** The encoding strategy for SuperBitSeq follows the commutative criterion.

| Flow | Operation to AGATGC |
|---|---|
| MK | $AGATGC \xrightarrow{encode} AGAGTC \xrightarrow{rc} GACTCT$<br>$AGATGC \xrightarrow{rc} GCATCT \xrightarrow{encode} GACTCT$ |
| RY | $AGATGC \xrightarrow{encode} AAGTGC \xrightarrow{rc} GCACTT$<br>$AGATGC \xrightarrow{rc} GCATCT \xrightarrow{encode} GCACTT$ |
| WS | $AGATGC \xrightarrow{encode} AGATCG \xrightarrow{rc} CGATCT$<br>$AGATGC \xrightarrow{rc} GCATCT \xrightarrow{encode} CGATCT$ |

**Table S7.** Total mapping rate of simulated error-free sequences using BWA-MEM. (*Homo sapiens*, %)

| Cycle | CRT | bMK | bRY | bWS | sMK | sRY | sWS |
|---|---|---|---|---|---|---|---|
| 5 | 0.00 | 0.35 | 0.73 | 0.53 | 0.06 | 0.14 | 0.07 |
| 10 | 0.00 | 9.25 | 14.06 | 7.04 | 2.13 | 3.84 | 2.25 |
| 15 | 0.00 | 68.46 | 71.62 | 46.21 | 39.72 | 46.39 | 23.53 |
| 20 | 0.00 | 98.78 | 97.95 | 93.00 | 94.04 | 93.11 | 78.84 |
| 25 | 0.00 | 99.94 | 99.70 | 99.86 | 99.85 | 99.41 | 99.30 |
| 30 | 100 | 100 | 100 | 100 | 99.98 | 99.89 | 99.96 |
| 35 | 100 | 100 | 100 | 100 | 100 | 100 | 100 |
| 40 | 100 | 100 | 100 | 100 | 100 | 100 | 100 |
| 45 | 100 | 100 | 100 | 100 | 100 | 100 | 100 |
| 50 | 100 | 100 | 100 | 100 | 100 | 100 | 100 |
| 75 | 100 | 100 | 100 | 100 | 100 | 100 | 100 |
| 100 | 100 | 100 | 100 | 100 | 100 | 100 | 100 |
| 125 | 100 | 100 | 100 | 100 | 100 | 100 | 100 |
| 150 | 100 | 100 | 100 | 100 | 100 | 100 | 100 |

**Table S8.** Unique mapping rate of simulated error-free sequences using BWA-MEM. (*Homo sapiens*, %)

| Cycle | CRT | bMK | bRY | bWS | sMK | sRY | sWS |
|---|---|---|---|---|---|---|---|
| 5 | 0.00 | 0.01 | 0.02 | 0.00 | 0.00 | 0.02 | 0.00 |
| 10 | 0.00 | 1.93 | 2.82 | 0.92 | 1.73 | 3.29 | 1.93 |
| 15 | 0.00 | 35.92 | 37.20 | 18.93 | 33.12 | 37.83 | 20.66 |
| 20 | 0.00 | 79.54 | 70.31 | 62.82 | 80.91 | 79.42 | 67.79 |
| 25 | 0.00 | 87.06 | 77.49 | 82.66 | 89.09 | 88.23 | 86.48 |
| 30 | 81.30 | 89.73 | 80.96 | 87.23 | 91.27 | 91.00 | 89.52 |
| 35 | 84.06 | 91.42 | 83.61 | 89.53 | 92.57 | 92.48 | 91.29 |
| 40 | 86.23 | 92.51 | 85.75 | 91.12 | 93.39 | 93.36 | 92.48 |
| 45 | 87.95 | 93.23 | 87.43 | 92.24 | 93.95 | 93.91 | 93.28 |
| 50 | 89.33 | 93.74 | 88.75 | 93.04 | 94.35 | 94.32 | 93.84 |
| 75 | 92.92 | 95.07 | 92.57 | 94.71 | 95.49 | 95.50 | 95.17 |
| 100 | 94.19 | 95.74 | 94.19 | 95.45 | 96.08 | 96.13 | 95.81 |
| 125 | 94.86 | 96.17 | 95.02 | 95.91 | 96.47 | 96.54 | 96.22 |
| 150 | 95.32 | 96.49 | 95.56 | 96.24 | 96.77 | 96.85 | 96.53 |

**Table S9.** Q20 mapping rate of simulated error-free sequences using BWA-MEM. (*Homo sapiens*, %)

| Cycle | CRT | bMK | bRY | bWS | sMK | sRY | sWS |
|---|---|---|---|---|---|---|---|
| 5 | 0.00 | 0.00 | 0.00 | 0.00 | 0.00 | 0.00 | 0.00 |
| 10 | 0.00 | 0.27 | 0.47 | 0.09 | 1.28 | 2.30 | 0.86 |
| 15 | 0.00 | 12.58 | 15.51 | 5.23 | 31.30 | 35.61 | 19.46 |
| 20 | 0.00 | 60.59 | 55.50 | 36.08 | 77.62 | 75.70 | 65.02 |
| 25 | 0.00 | 81.19 | 70.67 | 71.44 | 85.76 | 83.96 | 83.15 |
| 30 | 75.89 | 84.64 | 74.48 | 81.53 | 88.06 | 86.73 | 86.05 |
| 35 | 79.77 | 87.28 | 77.66 | 84.42 | 90.07 | 89.14 | 87.79 |
| 40 | 82.71 | 89.22 | 80.32 | 86.72 | 91.53 | 90.84 | 89.57 |
| 45 | 85.06 | 90.54 | 82.49 | 88.60 | 92.47 | 91.87 | 90.97 |
| 50 | 83.54 | 91.42 | 84.16 | 90.01 | 93.10 | 92.55 | 91.99 |
| 75 | 89.69 | 93.43 | 89.30 | 92.97 | 94.61 | 94.19 | 94.03 |
| 100 | 92.12 | 93.80 | 91.11 | 93.72 | 94.96 | 94.65 | 94.74 |
| 125 | 93.10 | 94.42 | 92.44 | 94.08 | 95.43 | 95.18 | 95.02 |
| 150 | 93.73 | 94.86 | 93.20 | 94.58 | 95.79 | 95.58 | 95.42 |

**Table S10.** Q30 mapping rate of simulated error-free sequences using BWA-MEM. (*Homo sapiens*, %)

| Cycle | CRT | bMK | bRY | bWS | sMK | sRY | sWS |
|---|---|---|---|---|---|---|---|
| 5 | 0.00 | 0.00 | 0.00 | 0.00 | 0.00 | 0.00 | 0.00 |
| 10 | 0.00 | 0.14 | 0.23 | 0.04 | 0.96 | 1.64 | 0.56 |
| 15 | 0.00 | 8.44 | 10.96 | 3.13 | 30.03 | 33.91 | 18.18 |
| 20 | 0.00 | 51.97 | 48.78 | 27.73 | 75.59 | 73.29 | 63.23 |
| 25 | 0.00 | 78.65 | 68.04 | 65.34 | 84.21 | 82.12 | 81.14 |
| 30 | 72.67 | 83.15 | 72.50 | 79.35 | 87.00 | 85.44 | 84.55 |
| 35 | 76.63 | 85.55 | 75.29 | 82.89 | 89.02 | 87.76 | 86.71 |
| 40 | 79.62 | 87.36 | 77.67 | 85.02 | 90.36 | 89.42 | 88.47 |
| 45 | 82.10 | 88.83 | 79.80 | 86.69 | 91.36 | 90.58 | 89.78 |
| 50 | 82.28 | 89.92 | 81.59 | 88.15 | 92.10 | 91.41 | 90.81 |
| 75 | 87.81 | 92.39 | 87.23 | 91.85 | 93.89 | 93.36 | 93.29 |
| 100 | 90.76 | 93.09 | 89.85 | 92.83 | 94.46 | 94.04 | 94.11 |
| 125 | 92.05 | 93.78 | 91.42 | 93.41 | 94.99 | 94.64 | 94.54 |
| 150 | 92.78 | 94.27 | 92.30 | 93.96 | 95.35 | 95.08 | 94.97 |

**Table S11.** Correct mapping rate of simulated error-free sequences using BWA-MEM. (*Homo sapiens*, %)

| Cycle | CRT | bMK | bRY | bWS | sMK | sRY | sWS |
|---|---|---|---|---|---|---|---|
| 5 | 0.00 | 0.01 | 0.03 | 0.01 | 0.01 | 0.04 | 0.01 |
| 10 | 0.00 | 3.23 | 4.49 | 1.50 | 1.79 | 3.42 | 2.02 |
| 15 | 0.00 | 43.93 | 43.94 | 24.47 | 34.23 | 39.12 | 21.21 |
| 20 | 0.00 | 83.46 | 74.25 | 69.41 | 83.29 | 81.83 | 69.74 |
| 25 | 0.00 | 89.54 | 80.47 | 85.84 | 91.30 | 90.53 | 88.84 |
| 30 | 84.18 | 91.89 | 83.71 | 89.65 | 93.23 | 93.02 | 91.68 |
| 35 | 86.75 | 93.33 | 86.19 | 91.69 | 94.33 | 94.29 | 93.24 |
| 40 | 88.74 | 94.26 | 88.21 | 93.08 | 95.03 | 95.01 | 94.26 |
| 45 | 90.30 | 94.87 | 89.73 | 94.03 | 95.50 | 95.48 | 94.94 |
| 50 | 91.53 | 95.30 | 90.94 | 94.69 | 95.85 | 95.83 | 95.41 |
| 75 | 94.63 | 96.44 | 94.36 | 96.09 | 96.82 | 96.83 | 96.52 |
| 100 | 95.70 | 97.00 | 95.78 | 96.73 | 97.31 | 97.36 | 97.06 |
| 125 | 96.28 | 97.37 | 96.49 | 97.12 | 97.62 | 97.69 | 97.41 |
| 150 | 96.68 | 97.63 | 96.93 | 97.40 | 97.87 | 97.94 | 97.66 |

**Table S12.** Correct and unique mapping rate of simulated error-free sequences using BWA-MEM. (*Homo sapiens*, %)

| Cycle | CRT | bMK | bRY | bWS | sMK | sRY | sWS |
|---|---|---|---|---|---|---|---|
| 5 | 0.00 | 0.00 | 0.01 | 0.00 | 0.01 | 0.03 | 0.01 |
| 10 | 0.00 | 1.94 | 2.81 | 0.92 | 1.72 | 3.30 | 1.94 |
| 15 | 0.00 | 35.93 | 37.21 | 18.93 | 33.12 | 37.83 | 20.66 |
| 20 | 0.00 | 79.54 | 70.32 | 62.82 | 80.91 | 79.42 | 67.79 |
| 25 | 0.00 | 87.06 | 77.49 | 82.66 | 89.09 | 88.23 | 86.48 |
| 30 | 81.29 | 89.73 | 80.95 | 87.23 | 91.27 | 91.00 | 89.53 |
| 35 | 84.06 | 91.41 | 83.60 | 89.53 | 92.56 | 92.49 | 91.30 |
| 40 | 86.23 | 92.51 | 85.76 | 91.12 | 93.39 | 93.35 | 92.48 |
| 45 | 87.94 | 93.24 | 87.42 | 92.24 | 93.95 | 93.91 | 93.29 |
| 50 | 89.32 | 93.75 | 88.74 | 93.04 | 94.36 | 94.33 | 93.85 |
| 75 | 92.91 | 95.08 | 92.56 | 94.71 | 95.50 | 95.49 | 95.17 |
| 100 | 94.18 | 95.74 | 94.19 | 95.45 | 96.08 | 96.13 | 95.80 |
| 125 | 94.86 | 96.18 | 95.02 | 95.91 | 96.46 | 96.54 | 96.22 |
| 150 | 95.33 | 96.50 | 95.55 | 96.24 | 96.77 | 96.85 | 96.53 |

**Table S13.** Total mapping rate of simulated error-free sequences using Bowtie2 and BWA-SW. (*Homo sapiens*, %)

| Software | Cycle | CRT | bMK | bRY | sMK | sRY | sWS |
|---|---|---|---|---|---|---|---|
| Bowtie2 | 5 | 100 | 100 | 100 | 100 | 100 | 100 |
| | 10 | 100 | 100 | 100 | 100 | 100 | 100 |
| | 15 | 100 | 100 | 100 | 100 | 100 | 100 |
| | 20 | 100 | 100 | 100 | 100 | 100 | 100 |
| | 25 | 100 | 100 | 100 | 100 | 100 | 100 |
| | 30 | 100 | 100 | 100 | 100 | 100 | 100 |
| | 35 | 100 | 100 | 100 | 100 | 100 | 100 |
| | 40 | 100 | 100 | 100 | 100 | 100 | 100 |
| | 45 | 100 | 100 | 100 | 100 | 100 | 100 |
| | 50 | 100 | 100 | 100 | 100 | 100 | 100 |
| | 75 | 100 | 100 | 100 | 100 | 100 | 100 |
| | 100 | 100 | 100 | 100 | 100 | 100 | 100 |
| | 125 | 100 | 100 | 100 | 100 | 100 | 100 |
| | 150 | 100 | 100 | 100 | 100 | 100 | 100 |
| BWA-SW | 5 | 0.00 | 0.35 | 0.73 | 0.06 | 0.14 | 0.07 |
| | 10 | 0.00 | 9.25 | 14.06 | 2.13 | 3.84 | 2.25 |
| | 15 | 0.00 | 68.46 | 71.62 | 39.72 | 46.39 | 23.53 |
| | 20 | 0.00 | 98.78 | 97.95 | 94.04 | 93.11 | 78.84 |
| | 25 | 0.00 | 99.94 | 99.70 | 99.85 | 99.41 | 99.30 |
| | 30 | 100 | 100 | 100 | 99.98 | 99.89 | 99.96 |
| | 35 | 100 | 100 | 100 | 100 | 100 | 100 |
| | 40 | 100 | 100 | 100 | 100 | 100 | 100 |
| | 45 | 100 | 100 | 100 | 100 | 100 | 100 |
| | 50 | 100 | 100 | 100 | 100 | 100 | 100 |
| | 75 | 100 | 100 | 100 | 100 | 100 | 100 |
| | 100 | 100 | 100 | 100 | 100 | 100 | 100 |
| | 125 | 100 | 100 | 100 | 100 | 100 | 100 |
| | 150 | 100 | 100 | 100 | 100 | 100 | 100 |

**Table S14.** Unique mapping rate of simulated error-free sequences using Bowtie2 and BWA-SW. (*Homo sapiens*, %)

| Software | Cycle | CRT | bMK | bRY | sMK | sRY | sWS |
|---|---|---|---|---|---|---|---|
| Bowtie2 | 5 | 0.00 | 0.01 | 0.02 | 0.01 | 0.02 | 0.01 |
| | 10 | 0.00 | 1.93 | 2.82 | 17.44 | 20.07 | 11.90 |
| | 15 | 2.90 | 35.93 | 37.23 | 77.94 | 76.35 | 72.28 |
| | 20 | 69.44 | 79.55 | 70.33 | 85.62 | 84.58 | 83.10 |
| | 25 | 77.62 | 87.06 | 77.49 | 89.16 | 88.53 | 86.92 |
| | 30 | 81.30 | 89.73 | 80.96 | 91.27 | 91.02 | 89.53 |
| | 35 | 84.06 | 91.42 | 83.61 | 92.57 | 92.48 | 91.29 |
| | 40 | 86.23 | 92.51 | 85.75 | 93.39 | 93.36 | 92.48 |
| | 45 | 87.95 | 93.23 | 87.42 | 93.95 | 93.91 | 93.28 |
| | 50 | 89.33 | 93.74 | 88.75 | 94.35 | 94.32 | 93.84 |
| | 75 | 92.92 | 95.07 | 92.57 | 95.49 | 95.50 | 95.16 |
| | 100 | 94.19 | 95.74 | 94.19 | 96.08 | 96.13 | 95.81 |
| | 125 | 94.86 | 96.17 | 95.02 | 96.47 | 96.54 | 96.22 |
| | 150 | 95.32 | 96.49 | 95.56 | 96.77 | 96.85 | 96.53 |
| BWA-SW | 5 | 0.00 | 0.01 | 0.02 | 0.00 | 0.02 | 0.00 |
| | 10 | 0.00 | 1.93 | 2.82 | 1.73 | 3.29 | 1.93 |
| | 15 | 0.00 | 35.92 | 37.20 | 33.12 | 37.83 | 20.66 |
| | 20 | 0.00 | 79.54 | 70.31 | 80.91 | 79.42 | 67.79 |
| | 25 | 0.00 | 87.06 | 77.49 | 89.09 | 88.23 | 86.48 |
| | 30 | 81.30 | 89.72 | 80.96 | 91.27 | 90.99 | 89.52 |
| | 35 | 84.06 | 91.41 | 83.60 | 92.56 | 92.48 | 91.29 |
| | 40 | 86.23 | 92.50 | 85.75 | 93.39 | 93.35 | 92.48 |
| | 45 | 87.95 | 93.23 | 87.42 | 93.94 | 93.91 | 93.28 |
| | 50 | 89.32 | 93.74 | 88.74 | 94.35 | 94.32 | 93.84 |
| | 75 | 92.92 | 95.07 | 92.56 | 95.48 | 95.50 | 95.16 |
| | 100 | 94.19 | 95.73 | 94.18 | 96.07 | 96.12 | 95.80 |
| | 125 | 94.86 | 96.16 | 95.01 | 96.46 | 96.53 | 96.22 |
| | 150 | 95.31 | 96.48 | 95.54 | 96.76 | 96.84 | 96.52 |

**Table S15.** Q20 mapping rate of simulated error-free sequences using Bowtie2 and BWA-SW. (*Homo sapiens*, %)

| Software | Cycle | CRT | bMK | bRY | sMK | sRY | sWS |
|---|---|---|---|---|---|---|---|
| Bowtie2 | 5 | 0.00 | 0.00 | 0.01 | 0.01 | 0.01 | 0.00 |
| | 10 | 0.00 | 1.93 | 2.80 | 17.44 | 20.07 | 11.90 |
| | 15 | 2.90 | 35.93 | 37.22 | 77.94 | 76.35 | 72.28 |
| | 20 | 69.44 | 79.55 | 70.32 | 85.62 | 84.58 | 83.10 |
| | 25 | 77.62 | 87.06 | 77.48 | 89.16 | 88.52 | 86.92 |
| | 30 | 81.30 | 89.72 | 80.92 | 91.27 | 91.01 | 89.52 |
| | 35 | 84.06 | 91.35 | 83.39 | 92.55 | 92.44 | 91.28 |
| | 40 | 86.23 | 92.14 | 84.81 | 93.30 | 93.13 | 92.45 |
| | 45 | 87.95 | 92.14 | 84.94 | 93.59 | 93.21 | 93.17 |
| | 50 | 89.33 | 92.07 | 85.34 | 93.64 | 93.17 | 93.49 |
| | 75 | 92.92 | 93.66 | 89.91 | 94.76 | 94.35 | 94.19 |
| | 100 | 92.30 | 94.54 | 92.28 | 95.18 | 94.87 | 94.90 |
| | 125 | 93.31 | 95.10 | 93.45 | 95.56 | 95.29 | 95.20 |
| | 150 | 93.90 | 95.48 | 94.12 | 95.73 | 95.49 | 95.48 |
| BWA-SW | 5 | 0.00 | 0.00 | 0.00 | 0.00 | 0.00 | 0.00 |
| | 10 | 0.00 | 0.00 | 0.00 | 0.01 | 0.07 | 0.03 |
| | 15 | 0.00 | 0.13 | 0.26 | 0.47 | 1.18 | 0.59 |
| | 20 | 0.00 | 4.17 | 5.96 | 8.74 | 13.79 | 6.49 |
| | 25 | 0.00 | 31.20 | 32.10 | 45.01 | 48.48 | 27.02 |
| | 30 | 0.00 | 66.17 | 57.76 | 76.49 | 74.37 | 60.56 |
| | 35 | 0.00 | 78.31 | 67.22 | 84.22 | 82.29 | 79.12 |
| | 40 | 0.00 | 82.26 | 71.34 | 87.27 | 85.92 | 83.91 |
| | 45 | 0.03 | 84.83 | 74.22 | 89.24 | 88.24 | 86.44 |
| | 50 | 77.01 | 86.70 | 76.55 | 90.52 | 89.72 | 88.35 |
| | 75 | 84.88 | 91.24 | 83.96 | 93.13 | 92.61 | 92.51 |
| | 100 | 89.77 | 92.31 | 87.65 | 93.86 | 93.37 | 93.45 |
| | 125 | 91.32 | 92.76 | 89.60 | 94.26 | 93.77 | 93.89 |
| | 150 | 92.34 | 93.05 | 90.56 | 94.52 | 94.04 | 94.19 |

**Table S16.** Q30 mapping rate of simulated error-free sequences using Bowtie2 and BWA-SW. (*Homo sapiens*, %)

| Software | Cycle | CRT | bMK | bRY | sMK | sRY | sWS |
|---|---|---|---|---|---|---|---|
| Bowtie2 | 5 | 0.00 | 0.00 | 0.01 | 0.01 | 0.01 | 0.00 |
| | 10 | 0.00 | 1.93 | 2.80 | 17.44 | 20.07 | 11.90 |
| | 15 | 2.90 | 35.93 | 37.22 | 77.94 | 76.35 | 72.28 |
| | 20 | 69.44 | 79.55 | 70.32 | 85.62 | 84.58 | 83.10 |
| | 25 | 77.62 | 87.06 | 77.48 | 89.16 | 88.52 | 86.92 |
| | 30 | 81.30 | 89.72 | 80.92 | 91.27 | 91.01 | 89.52 |
| | 35 | 84.06 | 91.35 | 83.39 | 92.55 | 92.44 | 91.28 |
| | 40 | 86.23 | 92.14 | 84.81 | 93.30 | 93.13 | 92.45 |
| | 45 | 87.95 | 92.14 | 84.94 | 93.59 | 93.21 | 93.17 |
| | 50 | 89.33 | 92.07 | 85.34 | 93.64 | 93.17 | 93.49 |
| | 75 | 92.92 | 93.66 | 89.91 | 94.76 | 94.35 | 94.19 |
| | 100 | 92.30 | 94.54 | 92.28 | 95.18 | 94.87 | 94.90 |
| | 125 | 93.31 | 95.10 | 93.45 | 95.56 | 95.29 | 95.20 |
| | 150 | 93.90 | 95.48 | 94.12 | 95.73 | 95.49 | 95.48 |
| BWA-SW | 5 | 0.00 | 0.00 | 0.00 | 0.00 | 0.00 | 0.00 |
| | 10 | 0.00 | 0.00 | 0.00 | 0.01 | 0.05 | 0.02 |
| | 15 | 0.00 | 0.07 | 0.14 | 0.09 | 0.23 | 0.18 |
| | 20 | 0.00 | 1.14 | 2.23 | 0.47 | 1.01 | 0.82 |
| | 25 | 0.00 | 8.13 | 11.79 | 3.62 | 6.61 | 3.14 |
| | 30 | 0.00 | 28.65 | 31.55 | 23.89 | 30.43 | 13.71 |
| | 35 | 0.00 | 58.58 | 54.16 | 63.34 | 64.62 | 38.39 |
| | 40 | 0.00 | 77.39 | 67.27 | 84.11 | 82.30 | 68.88 |
| | 45 | 0.03 | 83.47 | 72.41 | 88.40 | 87.07 | 83.83 |
| | 50 | 0.13 | 85.79 | 75.13 | 89.88 | 88.89 | 87.53 |
| | 75 | 83.71 | 90.30 | 82.41 | 92.48 | 91.80 | 91.82 |
| | 100 | 88.90 | 91.26 | 85.89 | 93.16 | 92.48 | 92.71 |
| | 125 | 90.53 | 91.67 | 87.89 | 93.52 | 92.85 | 93.12 |
| | 150 | 91.17 | 91.94 | 89.13 | 93.77 | 93.12 | 93.40 |

**Table S17.** Correct mapping rate of simulated error-free sequences using Bowtie2 and BWA-SW. (*Homo sapiens*, %)

| Software | Cycle | CRT | bMK | bRY | sMK | sRY | sWS |
|---|---|---|---|---|---|---|---|
| Bowtie2 | 5 | 0.01 | 0.01 | 0.03 | 0.02 | 0.04 | 0.02 |
| | 10 | 0.02 | 3.53 | 4.79 | 24.07 | 26.70 | 17.04 |
| | 15 | 11.36 | 45.11 | 45.13 | 81.57 | 80.08 | 76.84 |
| | 20 | 74.22 | 83.60 | 74.40 | 88.19 | 87.25 | 85.77 |
| | 25 | 80.77 | 89.55 | 80.48 | 91.38 | 90.86 | 89.31 |
| | 30 | 84.20 | 91.88 | 83.72 | 93.22 | 93.03 | 91.68 |
| | 35 | 86.75 | 93.34 | 86.20 | 94.33 | 94.29 | 93.23 |
| | 40 | 88.74 | 94.27 | 88.19 | 95.03 | 95.00 | 94.26 |
| | 45 | 90.30 | 94.87 | 89.73 | 95.49 | 95.48 | 94.94 |
| | 50 | 91.53 | 95.31 | 90.94 | 95.84 | 95.82 | 95.40 |
| | 75 | 94.63 | 96.43 | 94.35 | 96.82 | 96.83 | 96.51 |
| | 100 | 95.71 | 96.99 | 95.77 | 97.30 | 97.36 | 97.06 |
| | 125 | 96.29 | 97.37 | 96.49 | 97.62 | 97.68 | 97.41 |
| | 150 | 96.66 | 97.62 | 96.93 | 97.85 | 97.94 | 97.66 |
| BWA-SW | 5 | 0.00 | 0.01 | 0.04 | 0.01 | 0.04 | 0.01 |
| | 10 | 0.00 | 3.21 | 4.43 | 1.80 | 3.41 | 2.02 |
| | 15 | 0.00 | 43.82 | 43.78 | 34.19 | 39.09 | 21.22 |
| | 20 | 0.00 | 83.25 | 74.01 | 83.15 | 81.71 | 69.69 |
| | 25 | 0.00 | 89.39 | 80.26 | 91.19 | 90.40 | 88.71 |
| | 30 | 84.20 | 91.77 | 83.51 | 93.15 | 92.93 | 91.56 |
| | 35 | 86.48 | 93.26 | 86.04 | 94.27 | 94.23 | 93.15 |
| | 40 | 88.59 | 94.19 | 88.04 | 94.99 | 94.97 | 94.20 |
| | 45 | 90.17 | 94.81 | 89.61 | 95.46 | 95.44 | 94.88 |
| | 50 | 91.41 | 95.25 | 90.83 | 95.82 | 95.80 | 95.36 |
| | 75 | 94.58 | 96.41 | 94.31 | 96.79 | 96.81 | 96.49 |
| | 100 | 95.67 | 96.97 | 95.73 | 97.28 | 97.33 | 97.05 |
| | 125 | 96.25 | 97.34 | 96.44 | 97.61 | 97.67 | 97.38 |
| | 150 | 96.64 | 97.60 | 96.89 | 97.84 | 97.91 | 97.63 |

**Table S18.** Unique and correct mapping rate of simulated error-free sequences using Bowtie2 and BWA-SW. (*Homo sapiens*, %)

| Software | Cycle | CRT | bMK | bRY | sMK | sRY | sWS |
|---|---|---|---|---|---|---|---|
| Bowtie2 | 5 | 0.01 | 0.00 | 0.01 | 0.01 | 0.02 | 0.01 |
| | 10 | 0.00 | 1.94 | 2.82 | 17.44 | 20.08 | 11.89 |
| | 15 | 2.91 | 35.92 | 37.24 | 77.94 | 76.35 | 72.28 |
| | 20 | 69.44 | 79.54 | 70.33 | 85.63 | 84.58 | 83.10 |
| | 25 | 77.62 | 87.06 | 77.49 | 89.16 | 88.53 | 86.92 |
| | 30 | 81.30 | 89.72 | 80.96 | 91.26 | 91.01 | 89.53 |
| | 35 | 84.05 | 91.41 | 83.61 | 92.56 | 92.49 | 91.29 |
| | 40 | 86.22 | 92.51 | 85.75 | 93.39 | 93.35 | 92.49 |
| | 45 | 87.94 | 93.24 | 87.42 | 93.94 | 93.92 | 93.29 |
| | 50 | 89.32 | 93.75 | 88.75 | 94.35 | 94.32 | 93.85 |
| | 75 | 92.91 | 95.07 | 92.56 | 95.49 | 95.50 | 95.16 |
| | 100 | 94.19 | 95.74 | 94.18 | 96.08 | 96.13 | 95.80 |
| | 125 | 94.87 | 96.18 | 95.03 | 96.47 | 96.53 | 96.23 |
| | 150 | 95.31 | 96.49 | 95.56 | 96.76 | 96.85 | 96.53 |
| BWA-SW | 5 | 0.00 | 0.00 | 0.01 | 0.01 | 0.02 | 0.00 |
| | 10 | 0.00 | 1.94 | 2.81 | 1.73 | 3.29 | 1.93 |
| | 15 | 0.00 | 35.93 | 37.20 | 33.11 | 37.83 | 20.66 |
| | 20 | 0.00 | 79.53 | 70.32 | 80.89 | 79.42 | 67.79 |
| | 25 | 0.00 | 87.05 | 77.49 | 89.10 | 88.22 | 86.48 |
| | 30 | 81.30 | 89.72 | 80.95 | 91.27 | 90.99 | 89.52 |
| | 35 | 84.06 | 91.42 | 83.61 | 92.56 | 92.49 | 91.29 |
| | 40 | 86.24 | 92.50 | 85.74 | 93.39 | 93.36 | 92.49 |
| | 45 | 87.95 | 93.23 | 87.42 | 93.94 | 93.91 | 93.28 |
| | 50 | 89.32 | 93.74 | 88.75 | 94.36 | 94.33 | 93.84 |
| | 75 | 92.92 | 95.07 | 92.56 | 95.48 | 95.49 | 95.16 |
| | 100 | 94.19 | 95.73 | 94.18 | 96.07 | 96.11 | 95.80 |
| | 125 | 94.86 | 96.17 | 95.00 | 96.46 | 96.53 | 96.21 |
| | 150 | 95.31 | 96.48 | 95.53 | 96.76 | 96.83 | 96.51 |

**Table S19.** Total mapping rate of simulated error-free sequences. (*Arabidopsis thaliana*, %)

| Software | Cycle | CRT | bMK | bRY | bWS | sMK | sRY | sWS |
|---|---|---|---|---|---|---|---|---|
| Bowtie2 | 5 | 100 | 100 | 100 | 100 | 100 | 100 | 100 |
| | 10 | 100 | 100 | 100 | 100 | 100 | 100 | 100 |
| | 15 | 100 | 100 | 100 | 100 | 100 | 100 | 100 |
| | 20 | 100 | 100 | 100 | 100 | 100 | 100 | 100 |
| | 25 | 100 | 100 | 100 | 100 | 100 | 100 | 100 |
| | 30 | 100 | 100 | 100 | 100 | 100 | 100 | 100 |
| | 35 | 100 | 100 | 100 | 100 | 100 | 100 | 100 |
| | 40 | 100 | 100 | 100 | 100 | 100 | 100 | 100 |
| | 45 | 100 | 100 | 100 | 100 | 100 | 100 | 100 |
| | 50 | 100 | 100 | 100 | 100 | 100 | 100 | 100 |
| | 75 | 100 | 100 | 100 | 100 | 100 | 100 | 100 |
| | 100 | 100 | 100 | 100 | 100 | 100 | 100 | 100 |
| | 125 | 100 | 100 | 100 | 100 | 100 | 100 | 100 |
| | 150 | 100 | 100 | 100 | 100 | 100 | 100 | 100 |
| BWA-MEM | 5 | 0.00 | 0.15 | 0.36 | 0.77 | 0.02 | 0.04 | 0.08 |
| | 10 | 0.00 | 8.54 | 11.47 | 10.12 | 2.02 | 3.08 | 3.66 |
| | 15 | 0.00 | 61.60 | 67.06 | 55.35 | 34.74 | 40.67 | 30.92 |
| | 20 | 0.00 | 97.51 | 97.93 | 96.19 | 90.02 | 92.00 | 86.21 |
| | 25 | 0.00 | 99.95 | 99.93 | 99.97 | 99.80 | 99.76 | 99.77 |
| | 30 | 100 | 100 | 100 | 100 | 99.98 | 99.98 | 99.99 |
| | 35 | 100 | 100 | 100 | 100 | 100 | 100 | 100 |
| | 40 | 100 | 100 | 100 | 100 | 100 | 100 | 100 |
| | 45 | 100 | 100 | 100 | 100 | 100 | 100 | 100 |
| | 50 | 100 | 100 | 100 | 100 | 100 | 100 | 100 |
| | 75 | 100 | 100 | 100 | 100 | 100 | 100 | 100 |
| | 100 | 100 | 100 | 100 | 100 | 100 | 100 | 100 |
| | 125 | 100 | 100 | 100 | 100 | 100 | 100 | 100 |
| | 150 | 100 | 100 | 100 | 100 | 100 | 100 | 100 |
| BWA-SW | 5 | 0.00 | 0.15 | 0.36 | 0.77 | 0.02 | 0.04 | 0.08 |
| | 10 | 0.00 | 8.54 | 11.47 | 10.12 | 2.02 | 3.08 | 3.66 |
| | 15 | 0.00 | 61.60 | 67.06 | 55.35 | 34.74 | 40.67 | 30.92 |
| | 20 | 0.00 | 97.51 | 97.93 | 96.19 | 90.02 | 92.00 | 86.21 |
| | 25 | 0.00 | 99.95 | 99.93 | 99.97 | 99.80 | 99.76 | 99.77 |
| | 30 | 100 | 100 | 100 | 100 | 99.98 | 99.98 | 99.99 |
| | 35 | 100 | 100 | 100 | 100 | 100 | 100 | 100 |
| | 40 | 100 | 100 | 100 | 100 | 100 | 100 | 100 |
| | 45 | 100 | 100 | 100 | 100 | 100 | 100 | 100 |
| | 50 | 100 | 100 | 100 | 100 | 100 | 100 | 100 |
| | 75 | 100 | 100 | 100 | 100 | 100 | 100 | 100 |
| | 100 | 100 | 100 | 100 | 100 | 100 | 100 | 100 |
| | 125 | 100 | 100 | 100 | 100 | 100 | 100 | 100 |
| | 150 | 100 | 100 | 100 | 100 | 100 | 100 | 100 |

**Table S20.** Unique mapping rate of simulated error-free sequences. (*Arabidopsis thaliana*, %)

| Software | Cycle | CRT | bMK | bRY | bWS | sMK | sRY | sWS |
|---|---|---|---|---|---|---|---|---|
| Bowtie2 | 5 | 0.00 | 0.03 | 0.05 | 0.02 | 0.04 | 0.05 | 0.03 |
| | 10 | 0.00 | 8.01 | 9.44 | 5.26 | 36.98 | 41.29 | 35.17 |
| | 15 | 46.77 | 61.38 | 62.99 | 47.98 | 89.05 | 89.16 | 89.01 |
| | 20 | 86.67 | 89.71 | 86.43 | 86.81 | 92.16 | 92.21 | 92.07 |
| | 25 | 89.26 | 92.78 | 89.43 | 92.61 | 93.56 | 93.60 | 93.46 |
| | 30 | 90.71 | 93.83 | 90.81 | 93.75 | 94.50 | 94.55 | 94.41 |
| | 35 | 91.76 | 94.59 | 91.83 | 94.51 | 95.17 | 95.23 | 95.08 |
| | 40 | 92.58 | 95.15 | 92.63 | 95.07 | 95.67 | 95.72 | 95.58 |
| | 45 | 93.23 | 95.58 | 93.27 | 95.50 | 96.04 | 96.09 | 95.96 |
| | 50 | 93.77 | 95.93 | 93.80 | 95.85 | 96.33 | 96.39 | 96.27 |
| | 75 | 95.47 | 96.93 | 95.47 | 96.88 | 97.19 | 97.23 | 97.14 |
| | 100 | 96.32 | 97.42 | 96.34 | 97.37 | 97.61 | 97.65 | 97.56 |
| | 125 | 96.82 | 97.72 | 96.86 | 97.68 | 97.88 | 97.91 | 97.83 |
| | 150 | 97.15 | 97.92 | 97.21 | 97.88 | 98.05 | 98.09 | 98.01 |
| BWA-MEM | 5 | 0.00 | 0.03 | 0.05 | 0.02 | 0.01 | 0.02 | 0.01 |
| | 10 | 0.00 | 6.29 | 7.84 | 4.05 | 1.86 | 2.83 | 3.34 |
| | 15 | 0.00 | 53.50 | 55.61 | 42.33 | 32.00 | 37.19 | 28.61 |
| | 20 | 0.00 | 88.70 | 85.58 | 85.82 | 83.28 | 85.02 | 79.87 |
| | 25 | 0.00 | 92.78 | 89.42 | 92.61 | 93.41 | 93.40 | 93.26 |
| | 30 | 90.71 | 93.83 | 90.81 | 93.75 | 94.50 | 94.55 | 94.41 |
| | 35 | 91.76 | 94.59 | 91.83 | 94.51 | 95.17 | 95.23 | 95.08 |
| | 40 | 92.58 | 95.15 | 92.63 | 95.07 | 95.67 | 95.72 | 95.58 |
| | 45 | 93.23 | 95.58 | 93.27 | 95.50 | 96.04 | 96.09 | 95.96 |
| | 50 | 93.77 | 95.93 | 93.80 | 95.85 | 96.33 | 96.39 | 96.27 |
| | 75 | 95.47 | 96.93 | 95.47 | 96.88 | 97.19 | 97.23 | 97.14 |
| | 100 | 96.32 | 97.42 | 96.34 | 97.37 | 97.61 | 97.65 | 97.56 |
| | 125 | 96.82 | 97.72 | 96.86 | 97.68 | 97.88 | 97.91 | 97.83 |
| | 150 | 97.15 | 97.92 | 97.21 | 97.88 | 98.05 | 98.09 | 98.01 |
| BWA-SW | 5 | 0.00 | 0.03 | 0.05 | 0.02 | 0.01 | 0.02 | 0.01 |
| | 10 | 0.00 | 6.29 | 7.84 | 4.05 | 1.86 | 2.83 | 3.34 |
| | 15 | 0.00 | 53.50 | 55.61 | 42.33 | 32.00 | 37.19 | 28.61 |
| | 20 | 0.00 | 88.70 | 85.58 | 85.81 | 83.28 | 85.02 | 79.87 |
| | 25 | 0.00 | 92.78 | 89.42 | 92.61 | 93.41 | 93.40 | 93.26 |
| | 30 | 90.71 | 93.83 | 90.81 | 93.75 | 94.50 | 94.55 | 94.41 |
| | 35 | 91.76 | 94.59 | 91.82 | 94.50 | 95.17 | 95.23 | 95.08 |
| | 40 | 92.58 | 95.14 | 92.62 | 95.07 | 95.67 | 95.72 | 95.58 |
| | 45 | 93.23 | 95.57 | 93.26 | 95.49 | 96.04 | 96.09 | 95.96 |
| | 50 | 93.77 | 95.92 | 93.79 | 95.84 | 96.33 | 96.38 | 96.26 |
| | 75 | 95.47 | 96.93 | 95.46 | 96.87 | 97.19 | 97.22 | 97.13 |
| | 100 | 96.31 | 97.42 | 96.33 | 97.36 | 97.61 | 97.65 | 97.56 |
| | 125 | 96.82 | 97.72 | 96.85 | 97.67 | 97.87 | 97.91 | 97.82 |
| | 150 | 97.14 | 97.92 | 97.20 | 97.88 | 98.05 | 98.08 | 98.01 |

**Table S21.** Q20 mapping rate of simulated error-free sequences. (*Arabidopsis thaliana*, %)

| Software | Cycle | CRT | bMK | bRY | bWS | sMK | sRY | sWS |
|---|---|---|---|---|---|---|---|---|
| Bowtie2 | 5 | 0.00 | 0.03 | 0.05 | 0.02 | 0.04 | 0.05 | 0.03 |
| | 10 | 0.00 | 8.01 | 9.44 | 5.26 | 36.98 | 41.29 | 35.17 |
| | 15 | 46.77 | 61.38 | 62.99 | 47.98 | 89.05 | 89.16 | 89.01 |
| | 20 | 86.67 | 89.71 | 86.43 | 86.80 | 92.16 | 92.21 | 92.07 |
| | 25 | 89.26 | 92.78 | 89.42 | 92.57 | 93.56 | 93.60 | 93.46 |
| | 30 | 90.71 | 93.82 | 90.78 | 93.66 | 94.50 | 94.54 | 94.39 |
| | 35 | 91.76 | 94.53 | 91.67 | 94.32 | 95.16 | 95.19 | 95.02 |
| | 40 | 92.58 | 94.89 | 92.02 | 94.77 | 95.61 | 95.57 | 95.47 |
| | 45 | 93.23 | 94.81 | 91.78 | 94.95 | 95.82 | 95.63 | 95.77 |
| | 50 | 93.77 | 94.54 | 91.48 | 94.89 | 95.86 | 95.51 | 95.90 |
| | 75 | 95.47 | 95.55 | 93.28 | 95.45 | 96.58 | 96.26 | 96.41 |
| | 100 | 94.52 | 96.28 | 94.46 | 96.17 | 96.96 | 96.65 | 96.90 |
| | 125 | 95.41 | 96.76 | 95.24 | 96.53 | 97.23 | 96.95 | 97.11 |
| | 150 | 95.96 | 97.01 | 95.70 | 96.80 | 97.35 | 97.09 | 97.29 |
| BWA-MEM | 5 | 0.00 | 0.00 | 0.00 | 0.00 | 0.00 | 0.00 | 0.00 |
| | 10 | 0.00 | 1.47 | 1.93 | 0.74 | 1.76 | 2.65 | 2.65 |
| | 15 | 0.00 | 29.91 | 33.14 | 20.09 | 31.47 | 36.50 | 28.13 |
| | 20 | 0.00 | 78.47 | 76.87 | 67.89 | 82.00 | 83.52 | 78.64 |
| | 25 | 0.00 | 89.24 | 84.65 | 88.63 | 91.78 | 91.22 | 91.73 |
| | 30 | 88.57 | 90.05 | 85.84 | 90.20 | 92.47 | 91.81 | 92.43 |
| | 35 | 90.07 | 91.10 | 87.24 | 91.06 | 93.24 | 92.66 | 92.98 |
| | 40 | 91.18 | 92.08 | 88.48 | 92.00 | 94.01 | 93.52 | 93.75 |
| | 45 | 92.07 | 92.86 | 89.46 | 92.78 | 94.64 | 94.18 | 94.38 |
| | 50 | 89.68 | 93.47 | 90.24 | 93.40 | 95.11 | 94.69 | 94.88 |
| | 75 | 92.63 | 95.30 | 92.77 | 95.18 | 96.41 | 96.10 | 96.24 |
| | 100 | 94.28 | 95.68 | 93.46 | 95.80 | 96.78 | 96.44 | 96.74 |
| | 125 | 95.17 | 96.22 | 94.45 | 96.14 | 97.12 | 96.85 | 97.02 |
| | 150 | 95.78 | 96.67 | 95.19 | 96.57 | 97.42 | 97.19 | 97.31 |
| BWA-SW | 5 | 0.00 | 0.00 | 0.00 | 0.00 | 0.00 | 0.00 | 0.00 |
| | 10 | 0.00 | 0.02 | 0.04 | 0.01 | 0.01 | 0.05 | 0.08 |
| | 15 | 0.00 | 1.13 | 1.61 | 0.54 | 0.55 | 1.13 | 1.64 |
| | 20 | 0.00 | 15.79 | 18.71 | 7.62 | 8.91 | 12.62 | 11.52 |
| | 25 | 0.00 | 55.93 | 57.05 | 38.18 | 43.77 | 49.83 | 37.86 |
| | 30 | 0.00 | 82.69 | 79.10 | 74.47 | 79.96 | 81.79 | 74.62 |
| | 35 | 0.00 | 89.14 | 84.93 | 87.73 | 90.60 | 90.45 | 89.71 |
| | 40 | 0.01 | 90.98 | 86.94 | 90.67 | 92.75 | 92.47 | 92.39 |
| | 45 | 0.04 | 91.79 | 87.81 | 91.71 | 93.66 | 93.30 | 93.45 |
| | 50 | 87.86 | 92.17 | 88.28 | 92.24 | 94.15 | 93.71 | 94.04 |
| | 75 | 91.90 | 93.47 | 90.07 | 93.49 | 95.39 | 94.85 | 95.27 |
| | 100 | 93.78 | 94.12 | 91.07 | 94.11 | 95.91 | 95.38 | 95.79 |
| | 125 | 93.85 | 94.53 | 91.75 | 94.52 | 96.22 | 95.70 | 96.11 |
| | 150 | 94.58 | 94.85 | 92.31 | 94.85 | 96.45 | 95.95 | 96.33 |

**Table S22.** Q30 mapping rate of simulated error-free sequences. (*Arabidopsis thaliana*, %)

| Software | Cycle | CRT | bMK | bRY | bWS | sMK | sRY | sWS |
|---|---|---|---|---|---|---|---|---|
| Bowtie2 | 5 | 0.00 | 0.03 | 0.05 | 0.02 | 0.04 | 0.05 | 0.03 |
| | 10 | 0.00 | 8.01 | 9.44 | 5.26 | 36.98 | 41.29 | 35.17 |
| | 15 | 46.77 | 61.38 | 62.99 | 47.98 | 89.05 | 89.16 | 89.01 |
| | 20 | 86.67 | 89.71 | 86.43 | 86.80 | 92.16 | 92.21 | 92.07 |
| | 25 | 89.26 | 92.78 | 89.42 | 92.57 | 93.56 | 93.60 | 93.46 |
| | 30 | 90.71 | 93.82 | 90.78 | 93.66 | 94.50 | 94.54 | 94.39 |
| | 35 | 91.76 | 94.53 | 91.67 | 94.32 | 95.16 | 95.19 | 95.02 |
| | 40 | 92.58 | 94.89 | 92.02 | 94.77 | 95.61 | 95.57 | 95.47 |
| | 45 | 93.23 | 94.81 | 91.78 | 94.95 | 95.82 | 95.63 | 95.77 |
| | 50 | 93.77 | 94.54 | 91.48 | 94.89 | 95.86 | 95.51 | 95.90 |
| | 75 | 95.47 | 95.55 | 93.28 | 95.45 | 96.58 | 96.26 | 96.41 |
| | 100 | 94.52 | 96.28 | 94.46 | 96.17 | 96.96 | 96.65 | 96.90 |
| | 125 | 95.41 | 96.76 | 95.24 | 96.53 | 97.23 | 96.95 | 97.11 |
| | 150 | 95.96 | 97.01 | 95.70 | 96.80 | 97.35 | 97.09 | 97.29 |
| BWA-MEM | 5 | 0.00 | 0.00 | 0.00 | 0.00 | 0.00 | 0.00 | 0.00 |
| | 10 | 0.00 | 0.86 | 1.15 | 0.39 | 1.61 | 2.38 | 2.10 |
| | 15 | 0.00 | 22.06 | 24.95 | 13.96 | 30.81 | 35.47 | 27.52 |
| | 20 | 0.00 | 71.14 | 70.23 | 58.64 | 80.43 | 81.32 | 77.07 |
| | 25 | 0.00 | 87.21 | 82.53 | 85.46 | 90.51 | 89.68 | 90.17 |
| | 30 | 85.77 | 89.05 | 84.46 | 89.03 | 91.72 | 90.98 | 91.50 |
| | 35 | 86.93 | 89.84 | 85.48 | 89.97 | 92.51 | 91.75 | 92.32 |
| | 40 | 88.00 | 90.43 | 86.34 | 90.58 | 93.04 | 92.30 | 92.94 |
| | 45 | 89.08 | 91.11 | 87.25 | 91.13 | 93.56 | 92.89 | 93.39 |
| | 50 | 88.99 | 91.77 | 88.08 | 91.72 | 94.07 | 93.46 | 93.87 |
| | 75 | 90.82 | 93.96 | 90.87 | 93.86 | 95.69 | 95.21 | 95.51 |
| | 100 | 92.68 | 94.75 | 92.12 | 94.82 | 96.26 | 95.83 | 96.20 |
| | 125 | 93.85 | 95.50 | 93.32 | 95.41 | 96.73 | 96.36 | 96.58 |
| | 150 | 94.64 | 96.04 | 94.19 | 95.95 | 97.08 | 96.77 | 96.95 |
| BWA-SW | 5 | 0.00 | 0.00 | 0.00 | 0.00 | 0.00 | 0.00 | 0.00 |
| | 10 | 0.00 | 0.01 | 0.02 | 0.01 | 0.01 | 0.02 | 0.04 |
| | 15 | 0.00 | 0.28 | 0.53 | 0.33 | 0.04 | 0.13 | 0.34 |
| | 20 | 0.00 | 2.82 | 4.17 | 3.06 | 0.27 | 0.66 | 1.52 |
| | 25 | 0.00 | 13.79 | 16.76 | 12.81 | 3.04 | 5.23 | 5.95 |
| | 30 | 0.00 | 41.56 | 44.37 | 34.35 | 21.33 | 27.50 | 21.60 |
| | 35 | 0.00 | 75.02 | 73.54 | 66.19 | 60.48 | 65.55 | 51.57 |
| | 40 | 0.01 | 88.00 | 83.92 | 85.37 | 86.72 | 87.23 | 81.88 |
| | 45 | 0.04 | 90.01 | 85.68 | 89.68 | 92.45 | 91.77 | 91.83 |
| | 50 | 0.11 | 90.46 | 86.27 | 90.50 | 93.25 | 92.50 | 93.09 |
| | 75 | 90.51 | 91.70 | 87.99 | 91.73 | 94.44 | 93.66 | 94.30 |
| | 100 | 92.23 | 92.43 | 89.02 | 92.48 | 95.01 | 94.24 | 94.88 |
| | 125 | 92.53 | 92.97 | 89.76 | 92.98 | 95.37 | 94.64 | 95.26 |
| | 150 | 92.72 | 93.37 | 90.37 | 93.39 | 95.62 | 94.92 | 95.52 |

**Table S23.** Correct mapping rate of simulated error-free sequences. (*Arabidopsis thaliana*, %)

| Software | Cycle | CRT | bMK | bRY | bWS | sMK | sRY | sWS |
|---|---|---|---|---|---|---|---|---|
| Bowtie2 | 5 | 0.00 | 0.10 | 0.13 | 0.07 | 0.12 | 0.14 | 0.10 |
| | 10 | 0.44 | 12.91 | 14.92 | 8.81 | 46.87 | 51.39 | 45.09 |
| | 15 | 63.49 | 70.50 | 71.96 | 57.97 | 92.39 | 92.47 | 92.40 |
| | 20 | 90.60 | 93.02 | 90.29 | 91.09 | 94.68 | 94.70 | 94.58 |
| | 25 | 92.48 | 95.16 | 92.49 | 95.04 | 95.73 | 95.77 | 95.67 |
| | 30 | 93.56 | 95.94 | 93.55 | 95.88 | 96.45 | 96.48 | 96.38 |
| | 35 | 94.34 | 96.51 | 94.33 | 96.46 | 96.93 | 96.97 | 96.87 |
| | 40 | 95.01 | 96.91 | 95.00 | 96.86 | 97.31 | 97.33 | 97.24 |
| | 45 | 95.50 | 97.22 | 95.45 | 97.16 | 97.57 | 97.60 | 97.49 |
| | 50 | 95.90 | 97.49 | 95.87 | 97.43 | 97.79 | 97.82 | 97.73 |
| | 75 | 97.15 | 98.18 | 97.15 | 98.13 | 98.37 | 98.39 | 98.33 |
| | 100 | 97.76 | 98.51 | 97.76 | 98.48 | 98.64 | 98.66 | 98.60 |
| | 125 | 98.12 | 98.70 | 98.15 | 98.68 | 98.80 | 98.84 | 98.78 |
| | 150 | 98.33 | 98.83 | 98.38 | 98.81 | 98.91 | 98.94 | 98.89 |
| BWA-MEM | 5 | 0.00 | 0.06 | 0.09 | 0.04 | 0.01 | 0.02 | 0.02 |
| | 10 | 0.00 | 7.16 | 9.13 | 5.45 | 1.91 | 2.91 | 3.44 |
| | 15 | 0.00 | 56.35 | 59.16 | 46.88 | 32.88 | 38.29 | 29.33 |
| | 20 | 0.00 | 91.57 | 89.13 | 89.34 | 85.45 | 87.29 | 81.93 |
| | 25 | 0.00 | 95.15 | 92.46 | 95.04 | 95.56 | 95.56 | 95.45 |
| | 30 | 93.55 | 95.94 | 93.55 | 95.90 | 96.44 | 96.47 | 96.38 |
| | 35 | 94.36 | 96.50 | 94.35 | 96.45 | 96.93 | 96.97 | 96.87 |
| | 40 | 94.99 | 96.92 | 94.97 | 96.87 | 97.30 | 97.33 | 97.23 |
| | 45 | 95.49 | 97.23 | 95.46 | 97.17 | 97.56 | 97.61 | 97.50 |
| | 50 | 95.90 | 97.48 | 95.87 | 97.43 | 97.77 | 97.81 | 97.73 |
| | 75 | 97.16 | 98.19 | 97.12 | 98.15 | 98.36 | 98.38 | 98.33 |
| | 100 | 97.76 | 98.51 | 97.75 | 98.48 | 98.63 | 98.66 | 98.60 |
| | 125 | 98.11 | 98.70 | 98.13 | 98.68 | 98.80 | 98.82 | 98.77 |
| | 150 | 98.33 | 98.83 | 98.37 | 98.81 | 98.91 | 98.93 | 98.89 |
| BWA-SW | 5 | 0.00 | 0.06 | 0.09 | 0.03 | 0.01 | 0.03 | 0.03 |
| | 10 | 0.00 | 7.13 | 9.12 | 5.40 | 1.90 | 2.92 | 3.44 |
| | 15 | 0.00 | 56.30 | 59.10 | 46.64 | 32.85 | 38.28 | 29.32 |
| | 20 | 0.00 | 91.48 | 89.01 | 89.29 | 85.41 | 87.24 | 81.92 |
| | 25 | 0.00 | 95.08 | 92.35 | 94.97 | 95.53 | 95.50 | 95.40 |
| | 30 | 93.57 | 95.87 | 93.43 | 95.84 | 96.40 | 96.42 | 96.33 |
| | 35 | 94.29 | 96.46 | 94.25 | 96.40 | 96.89 | 96.93 | 96.84 |
| | 40 | 94.90 | 96.86 | 94.89 | 96.82 | 97.28 | 97.31 | 97.21 |
| | 45 | 95.42 | 97.19 | 95.37 | 97.13 | 97.55 | 97.57 | 97.48 |
| | 50 | 95.84 | 97.45 | 95.80 | 97.39 | 97.75 | 97.79 | 97.70 |
| | 75 | 97.10 | 98.17 | 97.09 | 98.13 | 98.36 | 98.39 | 98.33 |
| | 100 | 97.72 | 98.52 | 97.73 | 98.47 | 98.64 | 98.67 | 98.61 |
| | 125 | 98.10 | 98.71 | 98.11 | 98.67 | 98.81 | 98.81 | 98.77 |
| | 150 | 98.31 | 98.83 | 98.35 | 98.80 | 98.91 | 98.93 | 98.88 |

**Table S24.** Unique and correct mapping rate of simulated error-free sequences. (*Arabidopsis thaliana*, %)

| Software | Cycle | CRT | bMK | bRY | bWS | sMK | sRY | sWS |
|---|---|---|---|---|---|---|---|---|
| Bowtie2 | 5 | 0.00 | 0.03 | 0.05 | 0.02 | 0.04 | 0.05 | 0.03 |
| | 10 | 0.00 | 8.01 | 9.44 | 5.26 | 36.98 | 41.29 | 35.17 |
| | 15 | 46.77 | 61.38 | 62.99 | 47.98 | 89.05 | 89.16 | 89.01 |
| | 20 | 86.67 | 89.71 | 86.43 | 86.81 | 92.16 | 92.21 | 92.07 |
| | 25 | 89.26 | 92.78 | 89.43 | 92.61 | 93.56 | 93.60 | 93.46 |
| | 30 | 90.71 | 93.83 | 90.81 | 93.75 | 94.50 | 94.55 | 94.41 |
| | 35 | 91.76 | 94.59 | 91.83 | 94.51 | 95.17 | 95.23 | 95.08 |
| | 40 | 92.58 | 95.15 | 92.63 | 95.07 | 95.67 | 95.72 | 95.58 |
| | 45 | 93.23 | 95.58 | 93.27 | 95.50 | 96.04 | 96.09 | 95.96 |
| | 50 | 93.77 | 95.93 | 93.80 | 95.85 | 96.33 | 96.39 | 96.27 |
| | 75 | 95.47 | 96.93 | 95.47 | 96.88 | 97.19 | 97.23 | 97.14 |
| | 100 | 96.32 | 97.42 | 96.34 | 97.37 | 97.61 | 97.65 | 97.56 |
| | 125 | 96.82 | 97.72 | 96.86 | 97.68 | 97.88 | 97.91 | 97.83 |
| | 150 | 97.15 | 97.92 | 97.21 | 97.88 | 98.05 | 98.09 | 98.01 |
| BWA-MEM | 5 | 0.00 | 0.03 | 0.05 | 0.02 | 0.01 | 0.02 | 0.01 |
| | 10 | 0.00 | 6.29 | 7.84 | 4.05 | 1.86 | 2.83 | 3.34 |
| | 15 | 0.00 | 53.50 | 55.61 | 42.33 | 32.00 | 37.19 | 28.61 |
| | 20 | 0.00 | 88.70 | 85.58 | 85.82 | 83.28 | 85.02 | 79.87 |
| | 25 | 0.00 | 92.78 | 89.42 | 92.61 | 93.41 | 93.40 | 93.26 |
| | 30 | 90.71 | 93.83 | 90.81 | 93.75 | 94.50 | 94.55 | 94.41 |
| | 35 | 91.76 | 94.59 | 91.83 | 94.51 | 95.17 | 95.23 | 95.08 |
| | 40 | 92.58 | 95.15 | 92.63 | 95.07 | 95.67 | 95.72 | 95.58 |
| | 45 | 93.23 | 95.58 | 93.27 | 95.50 | 96.04 | 96.09 | 95.96 |
| | 50 | 93.77 | 95.93 | 93.80 | 95.85 | 96.33 | 96.39 | 96.27 |
| | 75 | 95.47 | 96.93 | 95.47 | 96.88 | 97.19 | 97.23 | 97.14 |
| | 100 | 96.32 | 97.42 | 96.34 | 97.37 | 97.61 | 97.65 | 97.56 |
| | 125 | 96.82 | 97.72 | 96.86 | 97.68 | 97.88 | 97.91 | 97.83 |
| | 150 | 97.15 | 97.92 | 97.21 | 97.88 | 98.05 | 98.09 | 98.01 |
| BWA-SW | 5 | 0.00 | 0.03 | 0.05 | 0.02 | 0.01 | 0.02 | 0.01 |
| | 10 | 0.00 | 6.29 | 7.84 | 4.05 | 1.86 | 2.83 | 3.34 |
| | 15 | 0.00 | 53.50 | 55.61 | 42.33 | 32.00 | 37.19 | 28.61 |
| | 20 | 0.00 | 88.70 | 85.58 | 85.81 | 83.28 | 85.02 | 79.87 |
| | 25 | 0.00 | 92.78 | 89.42 | 92.61 | 93.41 | 93.40 | 93.26 |
| | 30 | 90.71 | 93.83 | 90.81 | 93.75 | 94.50 | 94.55 | 94.41 |
| | 35 | 91.76 | 94.59 | 91.82 | 94.50 | 95.17 | 95.23 | 95.08 |
| | 40 | 92.58 | 95.14 | 92.62 | 95.07 | 95.67 | 95.72 | 95.58 |
| | 45 | 93.23 | 95.57 | 93.26 | 95.49 | 96.04 | 96.09 | 95.96 |
| | 50 | 93.77 | 95.92 | 93.79 | 95.84 | 96.33 | 96.38 | 96.26 |
| | 75 | 95.47 | 96.93 | 95.46 | 96.87 | 97.19 | 97.22 | 97.13 |
| | 100 | 96.31 | 97.42 | 96.32 | 97.36 | 97.61 | 97.64 | 97.56 |
| | 125 | 96.82 | 97.72 | 96.85 | 97.67 | 97.87 | 97.91 | 97.82 |
| | 150 | 97.14 | 97.92 | 97.20 | 97.88 | 98.05 | 98.08 | 98.01 |

**Table S25.** Total mapping rate of simulated erroneous sequences using Bowtie2. (%)

| Cycle | bMK | | | | | bRY | | | | |
|---|---|---|---|---|---|---|---|---|---|---|
| | 0.25 | 0.50 | 1 | 2 | 4 | 0.25 | 0.50 | 1 | 2 | 4 |
| 15 | 99.66 | 99.31 | 98.68 | 97.57 | 95.71 | 99.55 | 99.11 | 98.30 | 96.87 | 94.52 |
| 20 | 97.67 | 95.42 | 91.30 | 84.06 | 72.93 | 97.68 | 95.43 | 91.36 | 84.21 | 73.16 |
| 25 | 95.28 | 90.75 | 82.62 | 68.79 | 48.74 | 95.70 | 91.66 | 84.18 | 71.74 | 53.54 |
| 30 | 93.79 | 87.93 | 77.53 | 60.56 | 37.83 | 94.41 | 89.22 | 79.80 | 64.70 | 44.21 |
| 35 | 92.58 | 85.68 | 73.68 | 54.86 | 31.48 | 93.33 | 87.22 | 76.31 | 59.54 | 38.38 |
| 40 | 91.41 | 83.60 | 70.13 | 49.93 | 26.56 | 92.27 | 85.31 | 73.08 | 55.03 | 33.82 |
| 45 | 90.26 | 81.56 | 66.82 | 45.57 | 22.56 | 91.21 | 83.39 | 70.03 | 50.95 | 30.04 |
| 50 | 89.12 | 79.56 | 63.64 | 41.57 | 19.34 | 90.13 | 81.55 | 67.05 | 47.22 | 26.87 |
| 75 | 83.67 | 70.28 | 49.99 | 26.89 | 10.36 | 84.98 | 72.74 | 54.24 | 33.17 | 17.42 |
| 100 | 78.57 | 62.13 | 39.70 | 18.38 | 7.26 | 80.02 | 64.86 | 44.12 | 24.39 | 13.22 |
| 125 | 73.84 | 55.12 | 31.87 | 13.44 | 6.09 | 75.28 | 57.71 | 36.09 | 18.67 | 10.82 |

**Table S26.** Total mapping rate of simulated erroneous sequences using BWA-MEM. (%)

| Cycle | bMK | | | | | bRY | | | | |
|---|---|---|---|---|---|---|---|---|---|---|
| | 0.25 | 0.50 | 1 | 2 | 4 | 0.25 | 0.50 | 1 | 2 | 4 |
| 15 | 68.51 | 68.56 | 68.70 | 69.00 | 69.61 | 71.62 | 71.68 | 71.77 | 71.97 | 72.43 |
| 20 | 98.79 | 98.79 | 98.79 | 98.80 | 98.82 | 97.92 | 97.91 | 97.91 | 97.92 | 97.94 |
| 25 | 99.94 | 99.94 | 99.94 | 99.94 | 99.95 | 99.70 | 99.70 | 99.71 | 99.71 | 99.73 |
| 30 | 100 | 100 | 100 | 100 | 100 | 100 | 100 | 100 | 100 | 100 |
| 35 | 100 | 100 | 100 | 100 | 100 | 100 | 100 | 100 | 100 | 100 |
| 40 | 100 | 100 | 100 | 100 | 100 | 100 | 100 | 100 | 100 | 100 |
| 45 | 100 | 100 | 100 | 100 | 100 | 100 | 100 | 100 | 100 | 100 |
| 50 | 100 | 100 | 100 | 100 | 100 | 100 | 100 | 100 | 100 | 100 |
| 75 | 100 | 100 | 100 | 100 | 100 | 100 | 100 | 100 | 100 | 100 |
| 100 | 100 | 100 | 100 | 100 | 100 | 100 | 100 | 100 | 100 | 100 |
| 125 | 100 | 100 | 100 | 100 | 100 | 100 | 100 | 100 | 100 | 100 |

**Table S27.** Total mapping rate of simulated erroneous sequences using BWA-SW. (%)

| Cycle | bMK | | | | | bRY | | | | |
|---|---|---|---|---|---|---|---|---|---|---|
| | 0.25 | 0.50 | 1 | 2 | 4 | 0.25 | 0.50 | 1 | 2 | 4 |
| 15 | 68.50 | 68.55 | 68.67 | 68.94 | 69.52 | 71.60 | 71.63 | 71.67 | 71.79 | 72.14 |
| 20 | 98.74 | 98.69 | 98.59 | 98.43 | 98.19 | 97.81 | 97.72 | 97.53 | 97.18 | 96.63 |
| 25 | 99.89 | 99.84 | 99.73 | 99.54 | 99.30 | 99.59 | 99.49 | 99.30 | 98.96 | 98.41 |
| 30 | 99.95 | 99.90 | 99.81 | 99.65 | 99.43 | 99.90 | 99.83 | 99.65 | 99.36 | 98.91 |
| 35 | 99.96 | 99.93 | 99.87 | 99.77 | 99.63 | 99.94 | 99.90 | 99.79 | 99.61 | 99.37 |
| 40 | 99.98 | 99.97 | 99.94 | 99.90 | 99.85 | 99.98 | 99.96 | 99.92 | 99.85 | 99.74 |
| 45 | 100 | 99.99 | 99.98 | 99.97 | 99.96 | 99.99 | 99.99 | 99.98 | 99.95 | 99.91 |
| 50 | 100 | 100 | 100 | 99.99 | 99.99 | 100 | 100 | 99.99 | 99.98 | 99.96 |
| 75 | 100 | 100 | 100 | 100 | 100 | 100 | 100 | 100 | 100 | 100 |
| 100 | 100 | 100 | 100 | 100 | 100 | 100 | 100 | 100 | 100 | 100 |
| 125 | 100 | 100 | 100 | 100 | 100 | 100 | 100 | 100 | 100 | 100 |

**Table S28.** Unique mapping rate of simulated erroneous sequences using Bowtie2. (%)

| Cycle | bMK | | | | | bRY | | | | |
|---|---|---|---|---|---|---|---|---|---|---|
| | 0.25 | 0.50 | 1 | 2 | 4 | 0.25 | 0.50 | 1 | 2 | 4 |
| 15 | 35.36 | 34.72 | 33.56 | 31.65 | 28.77 | 36.35 | 35.70 | 34.49 | 32.48 | 29.60 |
| 20 | 77.24 | 75.02 | 70.93 | 63.92 | 53.41 | 68.10 | 66.05 | 62.35 | 56.12 | 46.86 |
| 25 | 82.74 | 78.64 | 71.29 | 58.92 | 41.20 | 73.62 | 70.09 | 63.62 | 53.09 | 38.27 |
| 30 | 83.94 | 78.52 | 68.98 | 53.48 | 32.98 | 75.91 | 71.30 | 63.04 | 50.03 | 33.02 |
| 35 | 84.44 | 78.01 | 66.86 | 49.45 | 28.03 | 77.54 | 72.06 | 62.39 | 47.74 | 29.92 |
| 40 | 84.40 | 77.06 | 64.42 | 45.58 | 23.97 | 78.65 | 72.35 | 61.39 | 45.45 | 27.34 |
| 45 | 84.00 | 75.78 | 61.87 | 41.95 | 20.54 | 79.30 | 72.17 | 60.10 | 43.09 | 25.04 |
| 50 | 83.39 | 74.33 | 59.26 | 38.48 | 17.70 | 79.58 | 71.70 | 58.49 | 40.67 | 22.92 |
| 75 | 79.44 | 66.62 | 47.20 | 25.25 | 9.64 | 78.42 | 66.93 | 49.71 | 30.25 | 15.94 |
| 100 | 75.12 | 59.32 | 37.74 | 17.38 | 6.84 | 75.25 | 60.85 | 41.31 | 22.79 | 12.46 |
| 125 | 70.92 | 52.87 | 30.46 | 12.77 | 5.80 | 71.43 | 54.68 | 34.15 | 17.67 | 10.36 |

**Table S29.** Unique mapping rate of simulated erroneous sequences using BWA-MEM. (%)

| Cycle | bMK | | | | | bRY | | | | |
|---|---|---|---|---|---|---|---|---|---|---|
| | 0.25 | 0.50 | 1 | 2 | 4 | 0.25 | 0.50 | 1 | 2 | 4 |
| 15 | 35.95 | 35.91 | 35.84 | 35.83 | 35.93 | 36.98 | 36.94 | 36.93 | 36.97 | 37.27 |
| 20 | 79.24 | 78.93 | 78.29 | 77.14 | 74.94 | 70.08 | 69.98 | 69.68 | 69.13 | 68.22 |
| 25 | 87.00 | 86.94 | 86.78 | 86.31 | 84.72 | 77.49 | 77.58 | 77.70 | 77.79 | 77.53 |
| 30 | 89.72 | 89.77 | 89.81 | 89.71 | 88.96 | 81.06 | 81.22 | 81.52 | 81.92 | 82.12 |
| 35 | 91.42 | 91.47 | 91.54 | 91.58 | 91.17 | 83.73 | 83.89 | 84.22 | 84.70 | 85.10 |
| 40 | 92.51 | 92.55 | 92.63 | 92.69 | 92.50 | 85.85 | 86.02 | 86.32 | 86.78 | 87.22 |
| 45 | 93.24 | 93.27 | 93.33 | 93.41 | 93.36 | 87.51 | 87.68 | 87.93 | 88.34 | 88.73 |
| 50 | 93.74 | 93.77 | 93.84 | 93.92 | 93.93 | 88.83 | 88.98 | 89.22 | 89.56 | 89.91 |
| 75 | 95.07 | 95.09 | 95.14 | 95.20 | 95.29 | 92.61 | 92.69 | 92.83 | 93.00 | 93.12 |
| 100 | 95.75 | 95.76 | 95.79 | 95.84 | 95.90 | 94.23 | 94.28 | 94.36 | 94.48 | 94.55 |
| 125 | 96.17 | 96.19 | 96.21 | 96.25 | 96.31 | 95.05 | 95.09 | 95.16 | 95.24 | 95.30 |

**Table S30.** Unique mapping rate of simulated erroneous sequences using BWA-SW. (%)

| Cycle | bMK | | | | | bRY | | | | |
|---|---|---|---|---|---|---|---|---|---|---|
| | 0.25 | 0.50 | 1 | 2 | 4 | 0.25 | 0.50 | 1 | 2 | 4 |
| 15 | 36.01 | 36.03 | 36.05 | 36.24 | 36.63 | 37.03 | 37.02 | 37.12 | 37.32 | 37.85 |
| 20 | 79.29 | 79.04 | 78.48 | 77.49 | 75.52 | 70.08 | 69.96 | 69.70 | 69.17 | 68.21 |
| 25 | 86.95 | 86.81 | 86.43 | 85.35 | 82.63 | 77.43 | 77.45 | 77.37 | 77.10 | 75.86 |
| 30 | 89.69 | 89.64 | 89.41 | 88.36 | 85.35 | 81.03 | 81.13 | 81.24 | 81.04 | 79.81 |
| 35 | 91.43 | 91.43 | 91.27 | 90.48 | 87.67 | 83.74 | 83.90 | 84.12 | 84.14 | 83.00 |
| 40 | 92.56 | 92.60 | 92.57 | 92.09 | 89.70 | 85.91 | 86.13 | 86.43 | 86.61 | 85.62 |
| 45 | 93.31 | 93.38 | 93.44 | 93.18 | 91.08 | 87.61 | 87.84 | 88.15 | 88.41 | 87.50 |
| 50 | 93.83 | 93.91 | 94.03 | 93.89 | 91.96 | 88.94 | 89.17 | 89.50 | 89.77 | 88.84 |
| 75 | 95.18 | 95.29 | 95.47 | 95.63 | 94.83 | 92.76 | 92.95 | 93.25 | 93.49 | 92.92 |
| 100 | 95.88 | 95.98 | 96.14 | 96.33 | 96.09 | 94.39 | 94.57 | 94.83 | 95.02 | 94.67 |
| 125 | 96.31 | 96.43 | 96.58 | 96.74 | 96.70 | 95.23 | 95.40 | 95.63 | 95.81 | 95.66 |

**Table S31.** Q20 mapping rate of simulated erroneous sequences using Bowtie2. (%)

| Cycle | bMK | | | | | bRY | | | | |
|---|---|---|---|---|---|---|---|---|---|---|
| | 0.25 | 0.50 | 1 | 2 | 4 | 0.25 | 0.50 | 1 | 2 | 4 |
| 15 | 35.00 | 34.02 | 32.24 | 29.13 | 24.35 | 35.93 | 34.89 | 32.93 | 29.58 | 24.44 |
| 20 | 76.17 | 72.90 | 66.86 | 56.41 | 40.57 | 67.15 | 64.21 | 58.79 | 49.46 | 35.42 |
| 25 | 82.10 | 77.43 | 68.98 | 54.76 | 34.57 | 72.97 | 68.83 | 61.20 | 48.70 | 30.97 |
| 30 | 83.46 | 77.66 | 67.37 | 50.74 | 28.96 | 75.25 | 70.05 | 60.69 | 45.90 | 26.48 |
| 35 | 83.88 | 77.02 | 65.12 | 46.59 | 24.15 | 76.54 | 70.32 | 59.35 | 42.71 | 22.45 |
| 40 | 83.50 | 75.72 | 62.31 | 42.45 | 20.02 | 76.80 | 69.66 | 57.28 | 39.24 | 18.87 |
| 45 | 82.39 | 73.78 | 59.19 | 38.42 | 16.56 | 75.88 | 67.92 | 54.42 | 35.46 | 15.71 |
| 50 | 81.25 | 71.83 | 56.19 | 34.72 | 13.76 | 75.21 | 66.46 | 51.88 | 32.22 | 13.20 |
| 75 | 77.45 | 64.22 | 44.20 | 21.75 | 6.35 | 74.12 | 61.35 | 42.32 | 21.09 | 6.68 |
| 100 | 73.22 | 56.96 | 34.78 | 14.06 | 3.85 | 71.25 | 55.35 | 33.93 | 14.16 | 4.43 |
| 125 | 69.04 | 50.45 | 27.47 | 9.56 | 3.02 | 67.61 | 49.33 | 27.15 | 9.97 | 3.69 |

**Table S32.** Q20 mapping rate of simulated erroneous sequences using BWA-MEM. (%)

| Cycle | bMK | | | | | bRY | | | | |
|---|---|---|---|---|---|---|---|---|---|---|
| | 0.25 | 0.50 | 1 | 2 | 4 | 0.25 | 0.50 | 1 | 2 | 4 |
| 15 | 12.10 | 11.63 | 10.75 | 9.19 | 6.72 | 14.75 | 14.19 | 13.12 | 11.24 | 8.25 |
| 20 | 58.61 | 56.66 | 52.92 | 46.02 | 34.57 | 53.70 | 52.07 | 48.95 | 43.12 | 33.22 |
| 25 | 79.96 | 78.70 | 76.08 | 70.44 | 58.67 | 69.67 | 68.77 | 66.78 | 62.52 | 53.29 |
| 30 | 84.33 | 83.99 | 83.14 | 80.75 | 73.48 | 74.09 | 73.76 | 72.99 | 70.93 | 65.11 |
| 35 | 87.11 | 86.98 | 86.64 | 85.49 | 81.26 | 77.39 | 77.19 | 76.76 | 75.56 | 71.80 |
| 40 | 89.08 | 89.00 | 88.77 | 88.11 | 85.44 | 80.09 | 79.91 | 79.56 | 78.66 | 75.89 |
| 45 | 90.41 | 90.34 | 90.17 | 89.72 | 87.93 | 82.24 | 82.06 | 81.72 | 80.93 | 78.72 |
| 50 | 91.30 | 91.25 | 91.11 | 90.79 | 89.52 | 83.94 | 83.77 | 83.46 | 82.75 | 80.87 |
| 75 | 93.33 | 93.26 | 93.13 | 92.89 | 92.35 | 89.07 | 88.91 | 88.58 | 87.94 | 86.71 |
| 100 | 93.74 | 93.70 | 93.66 | 93.57 | 93.35 | 90.97 | 90.89 | 90.76 | 90.45 | 89.73 |
| 125 | 94.35 | 94.33 | 94.29 | 94.19 | 94.01 | 92.33 | 92.27 | 92.18 | 91.95 | 91.44 |

**Table S33.** Q20 mapping rate of simulated erroneous sequences using BWA-SW. (%)

| Cycle | bMK | | | | | bRY | | | | |
|---|---|---|---|---|---|---|---|---|---|---|
| | 0.25 | 0.50 | 1 | 2 | 4 | 0.25 | 0.50 | 1 | 2 | 4 |
| 15 | 0.12 | 0.11 | 0.10 | 0.07 | 0.05 | 0.20 | 0.19 | 0.16 | 0.13 | 0.08 |
| 20 | 3.90 | 3.64 | 3.19 | 2.48 | 1.47 | 5.49 | 5.16 | 4.54 | 3.52 | 2.09 |
| 25 | 29.23 | 27.36 | 24.00 | 18.20 | 10.28 | 30.11 | 28.34 | 24.99 | 19.33 | 11.26 |
| 30 | 62.72 | 59.36 | 52.97 | 41.44 | 24.48 | 55.04 | 52.52 | 47.50 | 38.39 | 23.97 |
| 35 | 75.95 | 73.53 | 68.39 | 57.70 | 38.25 | 65.41 | 63.63 | 59.70 | 51.48 | 35.45 |
| 40 | 81.15 | 79.87 | 76.56 | 68.18 | 48.66 | 70.30 | 69.21 | 66.55 | 59.72 | 43.64 |
| 45 | 84.16 | 83.34 | 80.98 | 73.96 | 54.69 | 73.50 | 72.69 | 70.60 | 64.76 | 49.09 |
| 50 | 86.18 | 85.56 | 83.73 | 77.78 | 59.79 | 75.94 | 75.26 | 73.51 | 68.48 | 53.76 |
| 75 | 91.03 | 90.85 | 90.34 | 88.51 | 79.12 | 83.51 | 83.08 | 82.16 | 79.65 | 70.74 |
| 100 | 92.20 | 92.14 | 91.97 | 91.35 | 87.51 | 87.28 | 86.97 | 86.28 | 84.60 | 79.19 |
| 125 | 92.68 | 92.65 | 92.58 | 92.29 | 90.55 | 89.33 | 89.12 | 88.65 | 87.41 | 83.61 |

**Table S34.** Q30 mapping rate of simulated erroneous sequences using Bowtie2. (%)

| Cycle | bMK | | | | | bRY | | | | |
|---|---|---|---|---|---|---|---|---|---|---|
| | 0.25 | 0.50 | 1 | 2 | 4 | 0.25 | 0.50 | 1 | 2 | 4 |
| 15 | 34.87 | 33.77 | 31.76 | 28.26 | 22.86 | 35.79 | 34.61 | 32.41 | 28.64 | 22.86 |
| 20 | 75.92 | 72.42 | 65.98 | 54.79 | 37.96 | 66.96 | 63.82 | 58.06 | 48.16 | 33.30 |
| 25 | 82.00 | 77.24 | 68.62 | 54.12 | 33.61 | 72.86 | 68.62 | 60.80 | 47.99 | 29.88 |
| 30 | 83.38 | 77.51 | 67.08 | 50.21 | 28.17 | 75.13 | 69.83 | 60.27 | 45.14 | 25.35 |
| 35 | 83.79 | 76.86 | 64.78 | 46.03 | 23.30 | 76.40 | 70.06 | 58.85 | 41.86 | 21.22 |
| 40 | 83.41 | 75.56 | 62.00 | 41.91 | 19.18 | 76.65 | 69.38 | 56.75 | 38.33 | 17.64 |
| 45 | 82.30 | 73.61 | 58.90 | 37.90 | 15.78 | 75.72 | 67.62 | 53.88 | 34.59 | 14.50 |
| 50 | 81.16 | 71.67 | 55.89 | 34.23 | 12.97 | 75.05 | 66.15 | 51.35 | 31.37 | 12.02 |
| 75 | 77.36 | 64.06 | 43.93 | 21.30 | 5.55 | 73.93 | 61.00 | 41.77 | 20.31 | 5.53 |
| 100 | 73.11 | 56.79 | 34.51 | 13.62 | 3.01 | 71.03 | 54.97 | 33.36 | 13.43 | 3.30 |
| 125 | 68.93 | 50.26 | 27.17 | 9.16 | 2.16 | 67.38 | 48.95 | 26.58 | 9.30 | 2.55 |

**Table S35.** Q30 mapping rate of simulated erroneous sequences using BWA-MEM. (%)

| Cycle | bMK | | | | | bRY | | | | |
|---|---|---|---|---|---|---|---|---|---|---|
| | 0.25 | 0.50 | 1 | 2 | 4 | 0.25 | 0.50 | 1 | 2 | 4 |
| 15 | 8.08 | 7.75 | 7.10 | 6.01 | 4.28 | 10.38 | 9.95 | 9.14 | 7.72 | 5.49 |
| 20 | 50.03 | 48.12 | 44.48 | 37.89 | 27.35 | 46.92 | 45.23 | 42.04 | 36.22 | 26.64 |
| 25 | 76.93 | 75.20 | 71.73 | 64.65 | 51.10 | 66.70 | 65.47 | 62.80 | 57.39 | 46.61 |
| 30 | 82.62 | 82.07 | 80.69 | 77.18 | 67.56 | 71.96 | 71.48 | 70.31 | 67.40 | 59.85 |
| 35 | 85.30 | 85.09 | 84.56 | 82.95 | 77.29 | 74.95 | 74.68 | 74.09 | 72.48 | 67.73 |
| 40 | 87.17 | 87.05 | 86.75 | 85.92 | 82.64 | 77.38 | 77.15 | 76.72 | 75.64 | 72.47 |
| 45 | 88.66 | 88.57 | 88.36 | 87.82 | 85.74 | 79.51 | 79.31 | 78.93 | 78.03 | 75.63 |
| 50 | 89.78 | 89.71 | 89.55 | 89.15 | 87.70 | 81.33 | 81.13 | 80.77 | 79.98 | 78.01 |
| 75 | 92.27 | 92.21 | 92.09 | 91.87 | 91.37 | 87.00 | 86.85 | 86.56 | 85.97 | 84.69 |
| 100 | 93.01 | 92.98 | 92.93 | 92.82 | 92.55 | 89.69 | 89.60 | 89.43 | 89.05 | 88.19 |
| 125 | 93.70 | 93.67 | 93.63 | 93.51 | 93.25 | 91.30 | 91.23 | 91.10 | 90.84 | 90.22 |

**Table S36.** Q30 mapping rate of simulated erroneous sequences using BWA-SW. (%)

| Cycle | bMK | | | | | bRY | | | | |
|---|---|---|---|---|---|---|---|---|---|---|
| | 0.25 | 0.50 | 1 | 2 | 4 | 0.25 | 0.50 | 1 | 2 | 4 |
| 15 | 0.06 | 0.05 | 0.05 | 0.04 | 0.03 | 0.11 | 0.11 | 0.09 | 0.07 | 0.04 |
| 20 | 1.06 | 1.00 | 0.89 | 0.69 | 0.42 | 2.04 | 1.91 | 1.68 | 1.31 | 0.78 |
| 25 | 7.63 | 7.11 | 6.23 | 4.71 | 2.65 | 10.95 | 10.26 | 9.00 | 6.87 | 3.91 |
| 30 | 26.88 | 25.15 | 22.06 | 16.62 | 9.10 | 29.57 | 27.87 | 24.59 | 18.91 | 10.74 |
| 35 | 55.49 | 52.40 | 46.58 | 35.97 | 20.23 | 51.45 | 48.92 | 43.91 | 34.72 | 20.42 |
| 40 | 74.40 | 71.34 | 65.03 | 52.58 | 31.55 | 64.88 | 62.55 | 57.67 | 47.78 | 29.95 |
| 45 | 81.65 | 79.70 | 75.09 | 64.34 | 41.83 | 70.91 | 69.34 | 65.74 | 57.27 | 38.95 |
| 50 | 84.85 | 83.77 | 80.90 | 72.80 | 51.75 | 74.19 | 73.17 | 70.66 | 64.11 | 47.09 |
| 75 | 90.07 | 89.88 | 89.30 | 87.24 | 77.34 | 81.95 | 81.51 | 80.55 | 77.84 | 68.48 |
| 100 | 91.15 | 91.11 | 90.95 | 90.28 | 85.89 | 85.56 | 85.29 | 84.63 | 82.95 | 77.09 |
| 125 | 91.60 | 91.59 | 91.54 | 91.28 | 89.19 | 87.66 | 87.49 | 87.06 | 85.86 | 81.76 |

**Table S37.** Correct mapping rate of simulated erroneous sequences using Bowtie2. (%)

| Cycle | bMK | | | | | bRY | | | | |
|---|---|---|---|---|---|---|---|---|---|---|
| | 0.25 | 0.50 | 1 | 2 | 4 | 0.25 | 0.50 | 1 | 2 | 4 |
| 15 | 43.40 | 41.65 | 38.47 | 32.72 | 23.68 | 43.15 | 41.43 | 38.19 | 32.44 | 23.39 |
| 20 | 79.73 | 76.03 | 69.18 | 57.27 | 39.21 | 70.75 | 67.35 | 61.13 | 50.37 | 34.29 |
| 25 | 84.32 | 79.42 | 70.57 | 55.69 | 34.70 | 75.59 | 71.09 | 62.91 | 49.46 | 30.62 |
| 30 | 85.39 | 79.35 | 68.66 | 51.43 | 29.08 | 77.62 | 72.06 | 62.09 | 46.40 | 26.05 |
| 35 | 85.59 | 78.50 | 66.20 | 47.11 | 24.16 | 78.90 | 72.30 | 60.62 | 43.05 | 21.99 |
| 40 | 85.29 | 77.23 | 63.39 | 42.93 | 19.99 | 79.64 | 72.01 | 58.82 | 39.71 | 18.50 |
| 45 | 84.73 | 75.74 | 60.59 | 39.03 | 16.53 | 79.96 | 71.34 | 56.81 | 36.45 | 15.59 |
| 50 | 83.98 | 74.11 | 57.78 | 35.40 | 13.74 | 79.91 | 70.43 | 54.61 | 33.37 | 13.13 |
| 75 | 79.59 | 65.83 | 45.10 | 21.77 | 6.01 | 77.57 | 63.98 | 43.72 | 21.20 | 6.18 |
| 100 | 74.93 | 58.11 | 35.18 | 13.73 | 3.39 | 73.67 | 56.99 | 34.43 | 13.73 | 3.79 |
| 125 | 70.44 | 51.24 | 27.53 | 9.07 | 2.51 | 69.47 | 50.39 | 27.19 | 9.31 | 2.99 |

**Table S38.** Correct mapping rate of simulated erroneous sequences using BWA-MEM. (%)

| Cycle | bMK | | | | | bRY | | | | |
|---|---|---|---|---|---|---|---|---|---|---|
| | 0.25 | 0.50 | 1 | 2 | 4 | 0.25 | 0.50 | 1 | 2 | 4 |
| 15 | 42.55 | 41.18 | 38.58 | 33.80 | 25.85 | 42.39 | 41.07 | 38.59 | 33.97 | 26.21 |
| 20 | 81.67 | 79.91 | 76.37 | 69.46 | 56.60 | 72.74 | 71.33 | 68.54 | 62.85 | 52.16 |
| 25 | 88.87 | 88.16 | 86.54 | 82.72 | 73.33 | 79.81 | 79.19 | 77.82 | 74.53 | 66.66 |
| 30 | 91.64 | 91.39 | 90.69 | 88.66 | 82.43 | 83.36 | 83.04 | 82.30 | 80.33 | 74.76 |
| 35 | 93.17 | 93.03 | 92.66 | 91.48 | 87.23 | 85.90 | 85.67 | 85.16 | 83.80 | 79.69 |
| 40 | 94.13 | 94.06 | 93.81 | 93.08 | 90.18 | 87.92 | 87.73 | 87.28 | 86.22 | 83.05 |
| 45 | 94.77 | 94.70 | 94.54 | 94.05 | 92.01 | 89.48 | 89.31 | 88.92 | 88.04 | 85.45 |
| 50 | 95.21 | 95.16 | 95.05 | 94.68 | 93.17 | 90.70 | 90.54 | 90.22 | 89.44 | 87.25 |
| 75 | 96.39 | 96.36 | 96.31 | 96.13 | 95.51 | 94.23 | 94.13 | 93.93 | 93.39 | 92.09 |
| 100 | 96.97 | 96.95 | 96.91 | 96.75 | 96.28 | 95.70 | 95.63 | 95.51 | 95.18 | 94.26 |
| 125 | 97.33 | 97.31 | 97.27 | 97.14 | 96.72 | 96.42 | 96.39 | 96.29 | 96.07 | 95.37 |

**Table S39.** Correct mapping rate of simulated erroneous sequences using BWA-SW. (%)

| Cycle | bMK | | | | | bRY | | | | |
|---|---|---|---|---|---|---|---|---|---|---|
| | 0.25 | 0.50 | 1 | 2 | 4 | 0.25 | 0.50 | 1 | 2 | 4 |
| 15 | 42.41 | 41.07 | 38.47 | 33.66 | 25.62 | 42.22 | 40.93 | 38.40 | 33.75 | 25.88 |
| 20 | 81.22 | 79.19 | 75.05 | 67.03 | 52.29 | 72.21 | 70.55 | 67.12 | 60.25 | 47.36 |
| 25 | 88.03 | 86.60 | 83.20 | 75.44 | 58.71 | 79.01 | 77.78 | 74.85 | 68.15 | 53.60 |
| 30 | 90.95 | 89.89 | 87.07 | 79.59 | 61.84 | 82.65 | 81.66 | 79.17 | 72.78 | 57.54 |
| 35 | 92.62 | 91.84 | 89.53 | 83.02 | 66.06 | 85.32 | 84.52 | 82.40 | 76.73 | 62.17 |
| 40 | 93.67 | 93.09 | 91.34 | 86.08 | 70.70 | 87.41 | 86.72 | 84.99 | 80.16 | 66.39 |
| 45 | 94.38 | 93.91 | 92.55 | 88.29 | 74.00 | 89.02 | 88.43 | 86.93 | 82.70 | 69.63 |
| 50 | 94.84 | 94.45 | 93.36 | 89.70 | 76.37 | 90.28 | 89.73 | 88.35 | 84.59 | 72.29 |
| 75 | 96.05 | 95.74 | 95.03 | 93.17 | 85.54 | 93.83 | 93.39 | 92.37 | 89.85 | 81.72 |
| 100 | 96.62 | 96.32 | 95.65 | 94.18 | 89.49 | 95.33 | 94.95 | 94.04 | 92.06 | 86.18 |
| 125 | 97.00 | 96.69 | 96.04 | 94.64 | 91.02 | 96.09 | 95.71 | 94.94 | 93.23 | 88.54 |

**Table S40.** Unique and correct mapping rate of simulated erroneous sequences using Bowtie2. (%)

| Cycle | bMK | | | | | bRY | | | | |
|---|---|---|---|---|---|---|---|---|---|---|
| | 0.25 | 0.50 | 1 | 2 | 4 | 0.25 | 0.50 | 1 | 2 | 4 |
| 15 | 40.35 | 38.61 | 35.46 | 29.77 | 20.84 | 39.94 | 38.25 | 35.05 | 29.36 | 20.47 |
| 20 | 77.25 | 73.57 | 66.77 | 54.97 | 37.06 | 67.93 | 64.57 | 58.42 | 47.80 | 31.99 |
| 25 | 82.02 | 77.18 | 68.42 | 53.76 | 33.10 | 72.89 | 68.44 | 60.40 | 47.19 | 28.74 |
| 30 | 83.19 | 77.22 | 66.64 | 49.67 | 27.71 | 75.03 | 69.54 | 59.73 | 44.31 | 24.40 |
| 35 | 83.47 | 76.46 | 64.30 | 45.46 | 22.92 | 76.45 | 69.93 | 58.41 | 41.11 | 20.50 |
| 40 | 83.26 | 75.28 | 61.57 | 41.38 | 18.83 | 77.29 | 69.74 | 56.71 | 37.86 | 17.11 |
| 45 | 82.77 | 73.86 | 58.85 | 37.56 | 15.47 | 77.71 | 69.16 | 54.80 | 34.71 | 14.27 |
| 50 | 82.09 | 72.30 | 56.12 | 34.00 | 12.71 | 77.74 | 68.34 | 52.68 | 31.71 | 11.87 |
| 75 | 77.98 | 64.30 | 43.71 | 20.59 | 5.13 | 75.71 | 62.20 | 42.09 | 19.80 | 5.11 |
| 100 | 73.52 | 56.76 | 33.93 | 12.65 | 2.55 | 72.06 | 55.42 | 32.98 | 12.45 | 2.80 |
| 125 | 69.18 | 50.01 | 26.36 | 8.04 | 1.68 | 68.02 | 48.97 | 25.83 | 8.12 | 2.04 |

**Table S41.** Unique and correct mapping rate of simulated erroneous sequences using BWA-MEM. (%)

| Cycle | bMK | | | | | bRY | | | | |
|---|---|---|---|---|---|---|---|---|---|---|
| | 0.25 | 0.50 | 1 | 2 | 4 | 0.25 | 0.50 | 1 | 2 | 4 |
| 15 | 40.58 | 39.20 | 36.62 | 31.84 | 23.88 | 40.07 | 38.77 | 36.29 | 31.70 | 23.99 |
| 20 | 79.21 | 77.45 | 73.88 | 66.94 | 53.97 | 69.95 | 68.54 | 65.74 | 60.04 | 49.31 |
| 25 | 86.51 | 85.80 | 84.16 | 80.27 | 70.68 | 77.05 | 76.44 | 75.06 | 71.73 | 63.72 |
| 30 | 89.36 | 89.12 | 88.39 | 86.29 | 79.80 | 80.70 | 80.38 | 79.62 | 77.59 | 71.82 |
| 35 | 90.98 | 90.83 | 90.44 | 89.17 | 84.62 | 83.35 | 83.11 | 82.59 | 81.16 | 76.80 |
| 40 | 92.01 | 91.94 | 91.65 | 90.84 | 87.64 | 85.47 | 85.27 | 84.81 | 83.67 | 80.23 |
| 45 | 92.73 | 92.65 | 92.45 | 91.88 | 89.52 | 87.13 | 86.95 | 86.54 | 85.58 | 82.70 |
| 50 | 93.23 | 93.17 | 93.03 | 92.57 | 90.73 | 88.44 | 88.27 | 87.93 | 87.06 | 84.57 |
| 75 | 94.70 | 94.66 | 94.58 | 94.31 | 93.35 | 92.32 | 92.21 | 91.98 | 91.33 | 89.74 |
| 100 | 95.50 | 95.46 | 95.40 | 95.14 | 94.33 | 94.04 | 93.95 | 93.81 | 93.37 | 92.15 |
| 125 | 96.04 | 96.01 | 95.94 | 95.72 | 94.95 | 94.95 | 94.91 | 94.78 | 94.46 | 93.43 |

**Table S42.** Unique and correct mapping rate of simulated erroneous sequences using BWA-SW. (%)

| Cycle | bMK | | | | | bRY | | | | |
|---|---|---|---|---|---|---|---|---|---|---|
| | 0.25 | 0.50 | 1 | 2 | 4 | 0.25 | 0.50 | 1 | 2 | 4 |
| 15 | 40.80 | 39.44 | 36.79 | 31.91 | 23.75 | 40.28 | 38.97 | 36.40 | 31.68 | 23.72 |
| 20 | 78.88 | 76.73 | 72.32 | 63.91 | 48.73 | 69.51 | 67.74 | 64.09 | 56.91 | 43.63 |
| 25 | 85.55 | 83.92 | 80.14 | 71.80 | 54.38 | 76.19 | 74.79 | 71.55 | 64.36 | 49.20 |
| 30 | 88.48 | 87.19 | 83.96 | 75.82 | 57.30 | 79.86 | 78.67 | 75.81 | 68.84 | 52.89 |
| 35 | 90.19 | 89.19 | 86.40 | 79.13 | 61.13 | 82.58 | 81.57 | 79.05 | 72.67 | 57.17 |
| 40 | 91.30 | 90.48 | 88.22 | 82.07 | 65.37 | 84.75 | 83.84 | 81.65 | 76.04 | 61.05 |
| 45 | 92.07 | 91.35 | 89.45 | 84.22 | 68.43 | 86.44 | 85.62 | 83.63 | 78.54 | 64.08 |
| 50 | 92.58 | 91.93 | 90.30 | 85.64 | 70.65 | 87.76 | 86.98 | 85.09 | 80.42 | 66.60 |
| 75 | 94.06 | 93.49 | 92.20 | 89.20 | 79.40 | 91.63 | 90.93 | 89.36 | 85.78 | 75.64 |
| 100 | 94.84 | 94.26 | 93.01 | 90.36 | 83.27 | 93.37 | 92.71 | 91.23 | 88.15 | 80.04 |
| 125 | 95.40 | 94.81 | 93.57 | 90.95 | 84.87 | 94.32 | 93.65 | 92.30 | 89.46 | 82.42 |

**Table S43.** Total mapping rate of simulated random sequences. (%)

| Software | Bowtie2 | | | BWA-MEM | | | BWA-SW | | |
|---|---|---|---|---|---|---|---|---|---|
| Cycle | CRT | bMK | bRY | CRT | bMK | bRY | CRT | bMK | bRY |
| 5 | 100 | 100 | 100 | 0.00 | 0.01 | 0.00 | 0.00 | 0.01 | 0.00 |
| 10 | 100 | 99.98 | 99.99 | 0.00 | 3.07 | 3.02 | 0.00 | 3.07 | 3.02 |
| 15 | 99.77 | 97.25 | 97.42 | 0.00 | 49.80 | 49.31 | 0.00 | 49.53 | 48.20 |
| 20 | 16.09 | 66.88 | 65.73 | 0.00 | 96.45 | 95.88 | 0.00 | 93.56 | 88.51 |
| 25 | 0.05 | 18.44 | 16.66 | 0.00 | 99.98 | 99.97 | 0.00 | 96.31 | 91.43 |
| 30 | 0.00 | 2.03 | 1.85 | 0.00 | 100 | 100 | 0.00 | 96.49 | 91.99 |
| 35 | 0.00 | 0.15 | 0.15 | 0.00 | 100 | 100 | 0.00 | 97.23 | 93.97 |
| 40 | 0.00 | 0.01 | 0.01 | 0.00 | 100 | 100 | 0.00 | 98.52 | 96.82 |
| 45 | 0.00 | 0.00 | 0.00 | 0.00 | 100 | 100 | 0.00 | 99.42 | 98.54 |
| 50 | 0.00 | 0.00 | 0.00 | 0.00 | 100 | 100 | 0.00 | 99.76 | 99.15 |

**Table S44.** Unique mapping rate of simulated random sequences. (%)

| Software | Bowtie2 | | | BWA-MEM | | | BWA-SW | | |
|---|---|---|---|---|---|---|---|---|---|
| Cycle | CRT | bMK | bRY | CRT | bMK | bRY | CRT | bMK | bRY |
| 5 | 0.00 | 0.00 | 0.00 | 0.00 | 0.00 | 0.00 | 0.00 | 0.00 | 0.00 |
| 10 | 0.00 | 0.41 | 0.33 | 0.00 | 0.59 | 0.49 | 0.00 | 0.61 | 0.50 |
| 15 | 16.38 | 14.67 | 16.06 | 0.00 | 22.79 | 24.43 | 0.00 | 23.58 | 24.80 |
| 20 | 11.41 | 38.39 | 39.17 | 0.00 | 62.53 | 62.16 | 0.00 | 64.86 | 61.29 |
| 25 | 0.04 | 16.23 | 14.60 | 0.00 | 69.66 | 69.00 | 0.00 | 74.03 | 69.94 |
| 30 | 0.00 | 1.96 | 1.77 | 0.00 | 72.14 | 71.75 | 0.00 | 78.75 | 74.36 |
| 35 | 0.00 | 0.15 | 0.14 | 0.00 | 74.52 | 74.04 | 0.00 | 80.48 | 77.06 |
| 40 | 0.00 | 0.01 | 0.01 | 0.00 | 76.77 | 76.11 | 0.00 | 82.58 | 80.40 |
| 45 | 0.00 | 0.00 | 0.00 | 0.00 | 78.60 | 77.86 | 0.00 | 84.64 | 83.25 |
| 50 | 0.00 | 0.00 | 0.00 | 0.00 | 80.14 | 79.38 | 0.00 | 86.02 | 84.79 |

**Table S45.** Q20 mapping rate of simulated random sequences. (%)

| Software | Bowtie2 | | | BWA-MEM | | | BWA-SW | | |
|---|---|---|---|---|---|---|---|---|---|
| Cycle | CRT | bMK | bRY | CRT | bMK | bRY | CRT | bMK | bRY |
| 5 | 0.00 | 0.00 | 0.00 | 0.00 | 0.00 | 0.00 | 0.00 | 0.00 | 0.00 |
| 10 | 0.00 | 0.38 | 0.32 | 0.00 | 0.00 | 0.00 | 0.00 | 0.00 | 0.00 |
| 15 | 15.94 | 10.29 | 11.14 | 0.00 | 0.12 | 0.12 | 0.00 | 0.00 | 0.00 |
| 20 | 11.41 | 11.60 | 10.44 | 0.00 | 1.01 | 1.02 | 0.00 | 0.00 | 0.00 |
| 25 | 0.03 | 1.31 | 1.16 | 0.00 | 1.97 | 1.90 | 0.00 | 0.00 | 0.00 |
| 30 | 0.00 | 0.02 | 0.03 | 0.00 | 2.43 | 2.36 | 0.00 | 0.00 | 0.00 |
| 35 | 0.00 | 0.00 | 0.00 | 0.00 | 2.91 | 2.81 | 0.00 | 0.00 | 0.00 |
| 40 | 0.00 | 0.00 | 0.00 | 0.00 | 3.41 | 3.29 | 0.00 | 0.00 | 0.00 |
| 45 | 0.00 | 0.00 | 0.00 | 0.00 | 3.93 | 3.77 | 0.00 | 0.00 | 0.00 |
| 50 | 0.00 | 0.00 | 0.00 | 0.00 | 4.44 | 4.28 | 0.00 | 0.00 | 0.00 |

**Table S46.** Q30 mapping rate of simulated random sequences. (%)

| Software | Bowtie2 | | | BWA-MEM | | | BWA-SW | | |
|---|---|---|---|---|---|---|---|---|---|
| Cycle | CRT | bMK | bRY | CRT | bMK | bRY | CRT | bMK | bRY |
| 5 | 0.00 | 0.00 | 0.00 | 0.00 | 0.00 | 0.00 | 0.00 | 0.00 | 0.00 |
| 10 | 0.00 | 0.36 | 0.31 | 0.00 | 0.00 | 0.00 | 0.00 | 0.00 | 0.00 |
| 15 | 15.94 | 8.80 | 9.54 | 0.00 | 0.02 | 0.03 | 0.00 | 0.00 | 0.00 |
| 20 | 0.38 | 6.43 | 5.59 | 0.00 | 0.31 | 0.31 | 0.00 | 0.00 | 0.00 |
| 25 | 0.00 | 0.48 | 0.47 | 0.00 | 0.69 | 0.68 | 0.00 | 0.00 | 0.00 |
| 30 | 0.00 | 0.01 | 0.01 | 0.00 | 0.88 | 0.85 | 0.00 | 0.00 | 0.00 |
| 35 | 0.00 | 0.00 | 0.00 | 0.00 | 1.03 | 1.00 | 0.00 | 0.00 | 0.00 |
| 40 | 0.00 | 0.00 | 0.00 | 0.00 | 1.20 | 1.17 | 0.00 | 0.00 | 0.00 |
| 45 | 0.00 | 0.00 | 0.00 | 0.00 | 1.37 | 1.34 | 0.00 | 0.00 | 0.00 |
| 50 | 0.00 | 0.00 | 0.00 | 0.00 | 1.55 | 1.53 | 0.00 | 0.00 | 0.00 |

**Table S47.** Total mapping rate of error-free reads from a random genome. (%)

| Software | Bowtie2 | | | BWA-MEM | | | BWA-SW | | |
|---|---|---|---|---|---|---|---|---|---|
| Cycle | CRT | bMK | bRY | CRT | bMK | bRY | CRT | bMK | bRY |
| 5 | 100 | 100 | 100 | 0.00 | 0.01 | 0.00 | 0.00 | 0.01 | 0.00 |
| 10 | 100 | 100 | 100 | 0.00 | 3.07 | 3.07 | 0.00 | 0.00 | 0.00 |
| 15 | 100 | 100 | 100 | 0.00 | 50.00 | 50.01 | 0.00 | 0.00 | 0.00 |
| 20 | 100 | 100 | 100 | 0.00 | 96.92 | 96.93 | 0.00 | 96.92 | 96.93 |
| 25 | 100 | 100 | 100 | 0.00 | 99.99 | 100 | 0.00 | 99.99 | 100 |
| 30 | 100 | 100 | 100 | 100 | 100 | 100 | 100 | 100 | 100 |
| 35 | 100 | 100 | 100 | 100 | 100 | 100 | 100 | 100 | 100 |
| 40 | 100 | 100 | 100 | 100 | 100 | 100 | 100 | 100 | 100 |
| 45 | 100 | 100 | 100 | 100 | 100 | 100 | 100 | 100 | 100 |
| 50 | 100 | 100 | 100 | 100 | 100 | 100 | 100 | 100 | 100 |
| 75 | 100 | 100 | 100 | 100 | 100 | 100 | 100 | 100 | 100 |
| 100 | 100 | 100 | 100 | 100 | 100 | 100 | 100 | 100 | 100 |
| 125 | 100 | 100 | 100 | 100 | 100 | 100 | 100 | 100 | 100 |
| 150 | 100 | 100 | 100 | 100 | 100 | 100 | 100 | 100 | 100 |

**Table S48.** Unique mapping rate of error-free reads from a random genome. (%)

| Software | Bowtie2 | | | BWA-MEM | | | BWA-SW | | |
|---|---|---|---|---|---|---|---|---|---|
| Cycle | CRT | bMK | bRY | CRT | bMK | bRY | CRT | bMK | bRY |
| 5 | 0.00 | 0.00 | 0.00 | 0.00 | 0.00 | 0.00 | 0.00 | 0.00 | 0.00 |
| 10 | 0.00 | 0.89 | 0.90 | 0.00 | 0.89 | 0.90 | 0.00 | 0.00 | 0.00 |
| 15 | 0.37 | 26.49 | 26.50 | 0.00 | 26.49 | 26.50 | 0.00 | 0.00 | 0.00 |
| 20 | 99.46 | 84.87 | 84.87 | 0.00 | 84.87 | 84.87 | 0.00 | 84.87 | 84.87 |
| 25 | 100 | 99.56 | 99.55 | 0.00 | 99.56 | 99.55 | 0.00 | 99.56 | 99.55 |
| 30 | 100 | 100 | 100 | 100 | 100 | 100 | 100 | 100 | 100 |
| 35 | 100 | 100 | 100 | 100 | 100 | 100 | 100 | 100 | 100 |
| 40 | 100 | 100 | 100 | 100 | 100 | 100 | 100 | 100 | 100 |
| 45 | 100 | 100 | 100 | 100 | 100 | 100 | 100 | 100 | 100 |
| 50 | 100 | 100 | 100 | 100 | 100 | 100 | 100 | 100 | 100 |
| 75 | 100 | 100 | 100 | 100 | 100 | 100 | 100 | 100 | 100 |
| 100 | 100 | 100 | 100 | 100 | 100 | 100 | 100 | 100 | 100 |
| 125 | 100 | 100 | 100 | 100 | 100 | 100 | 100 | 100 | 100 |
| 150 | 100 | 100 | 100 | 100 | 100 | 100 | 100 | 100 | 100 |

**Table S49.** Q20 mapping rate of error-free reads from a random genome. (%)

| Software | Bowtie2 | | | BWA-MEM | | | BWA-SW | | |
|---|---|---|---|---|---|---|---|---|---|
| Cycle | CRT | bMK | bRY | CRT | bMK | bRY | CRT | bMK | bRY |
| 5 | 0.00 | 0.00 | 0.00 | 0.00 | 0.00 | 0.00 | 0.00 | 0.00 | 0.00 |
| 10 | 0.00 | 0.89 | 0.90 | 0.00 | 0.00 | 0.00 | 0.00 | 0.00 | 0.00 |
| 15 | 0.37 | 26.49 | 26.50 | 0.00 | 0.03 | 0.03 | 0.00 | 0.00 | 0.00 |
| 20 | 99.46 | 84.87 | 84.87 | 0.00 | 1.80 | 1.81 | 0.00 | 54.13 | 54.13 |
| 25 | 100 | 99.56 | 99.55 | 0.00 | 21.05 | 21.05 | 0.00 | 95.07 | 95.06 |
| 30 | 100 | 100 | 100 | 0.00 | 66.71 | 66.72 | 100 | 99.91 | 99.91 |
| 35 | 100 | 100 | 100 | 0.00 | 94.59 | 94.60 | 100 | 100 | 100 |
| 40 | 100 | 100 | 100 | 0.00 | 99.66 | 99.66 | 100 | 100 | 100 |
| 45 | 100 | 100 | 100 | 0.00 | 99.99 | 99.99 | 100 | 100 | 100 |
| 50 | 100 | 100 | 100 | 100 | 100 | 100 | 100 | 100 | 100 |
| 75 | 100 | 100 | 100 | 100 | 100 | 100 | 100 | 100 | 100 |
| 100 | 100 | 100 | 100 | 100 | 100 | 100 | 100 | 100 | 100 |
| 125 | 100 | 100 | 100 | 100 | 100 | 100 | 100 | 100 | 100 |
| 150 | 100 | 100 | 100 | 100 | 100 | 100 | 100 | 100 | 100 |

**Table S50.** Q30 mapping rate of error-free reads from a random genome. (%)

| Software | Bowtie2 | | | BWA-MEM | | | BWA-SW | | |
|---|---|---|---|---|---|---|---|---|---|
| Cycle | CRT | bMK | bRY | CRT | bMK | bRY | CRT | bMK | bRY |
| 5 | 0.00 | 0.00 | 0.00 | 0.00 | 0.00 | 0.00 | 0.00 | 0.00 | 0.00 |
| 10 | 0.00 | 0.89 | 0.90 | 0.00 | 0.00 | 0.00 | 0.00 | 0.00 | 0.00 |
| 15 | 0.37 | 26.49 | 26.50 | 0.00 | 0.00 | 0.00 | 0.00 | 0.00 | 0.00 |
| 20 | 99.46 | 84.87 | 84.87 | 0.00 | 0.14 | 0.14 | 0.00 | 43.81 | 43.79 |
| 25 | 100 | 99.56 | 99.55 | 0.00 | 2.48 | 2.48 | 0.00 | 91.26 | 91.25 |
| 30 | 100 | 100 | 100 | 0.00 | 16.50 | 16.51 | 100 | 99.72 | 99.72 |
| 35 | 100 | 100 | 100 | 0.00 | 49.87 | 49.88 | 100 | 99.99 | 99.99 |
| 40 | 100 | 100 | 100 | 0.00 | 83.45 | 83.45 | 100 | 100 | 100 |
| 45 | 100 | 100 | 100 | 0.00 | 97.43 | 97.43 | 100 | 100 | 100 |
| 50 | 100 | 100 | 100 | 0.00 | 99.82 | 99.82 | 100 | 100 | 100 |
| 75 | 100 | 100 | 100 | 100 | 100 | 100 | 100 | 100 | 100 |
| 100 | 100 | 100 | 100 | 100 | 100 | 100 | 100 | 100 | 100 |
| 125 | 100 | 100 | 100 | 100 | 100 | 100 | 100 | 100 | 100 |
| 150 | 100 | 100 | 100 | 100 | 100 | 100 | 100 | 100 | 100 |

**Table S51.** Correct mapping rate of error-free reads from a random genome. (%)

| Software | Bowtie2 | | | BWA-MEM | | | BWA-SW | | |
|---|---|---|---|---|---|---|---|---|---|
| Cycle | CRT | bMK | bRY | CRT | bMK | bRY | CRT | bMK | bRY |
| 5 | 0.00 | 0.01 | 0.00 | 0.00 | 0.01 | 0.00 | 0.00 | 0.01 | 0.00 |
| 10 | 0.01 | 1.83 | 1.84 | 0.00 | 100 | 100 | 0.00 | 1.50 | 1.50 |
| 15 | 17.84 | 35.14 | 35.15 | 0.00 | 100 | 100 | 0.00 | 33.84 | 33.86 |
| 20 | 99.72 | 89.51 | 89.53 | 0.00 | 89.35 | 89.35 | 0.00 | 89.34 | 89.34 |
| 25 | 100 | 99.75 | 99.75 | 0.00 | 99.74 | 99.75 | 0.00 | 99.74 | 99.75 |
| 30 | 100 | 100 | 100 | 100 | 100 | 100 | 100 | 100 | 100 |
| 35 | 100 | 100 | 100 | 100 | 100 | 100 | 100 | 100 | 100 |
| 40 | 100 | 100 | 100 | 100 | 100 | 100 | 100 | 100 | 100 |
| 45 | 100 | 100 | 100 | 100 | 100 | 100 | 100 | 100 | 100 |
| 50 | 100 | 100 | 100 | 100 | 100 | 100 | 100 | 100 | 100 |
| 75 | 100 | 100 | 100 | 100 | 100 | 100 | 100 | 100 | 100 |
| 100 | 100 | 100 | 100 | 100 | 100 | 100 | 100 | 100 | 100 |
| 125 | 100 | 100 | 100 | 100 | 100 | 100 | 100 | 100 | 100 |
| 150 | 100 | 100 | 100 | 100 | 100 | 100 | 100 | 100 | 100 |

**Table S52.** Correct and unique mapping rate of error-free reads from a random genome. (%)

| Software | Bowtie2 | | | BWA-MEM | | | BWA-SW | | |
|---|---|---|---|---|---|---|---|---|---|
| Cycle | CRT | bMK | bRY | CRT | bMK | bRY | CRT | bMK | bRY |
| 5 | 0.00 | 0.01 | 0.00 | 0.00 | 0.01 | 0.00 | 0.00 | 0.01 | 0.00 |
| 10 | 0.00 | 0.89 | 0.90 | 0.00 | 100 | 100 | 0.00 | 0.90 | 0.90 |
| 15 | 0.38 | 26.48 | 26.50 | 0.00 | 100 | 100 | 0.00 | 26.48 | 26.50 |
| 20 | 99.45 | 84.87 | 84.88 | 0.00 | 84.87 | 84.87 | 0.00 | 84.87 | 84.87 |
| 25 | 100 | 99.55 | 99.55 | 0.00 | 99.54 | 99.55 | 0.00 | 99.54 | 99.55 |
| 30 | 100 | 100 | 100 | 100 | 100 | 100 | 100 | 100 | 100 |
| 35 | 100 | 100 | 100 | 100 | 100 | 100 | 100 | 100 | 100 |
| 40 | 100 | 100 | 100 | 100 | 100 | 100 | 100 | 100 | 100 |
| 45 | 100 | 100 | 100 | 100 | 100 | 100 | 100 | 100 | 100 |
| 50 | 100 | 100 | 100 | 100 | 100 | 100 | 100 | 100 | 100 |
| 75 | 100 | 100 | 100 | 100 | 100 | 100 | 100 | 100 | 100 |
| 100 | 100 | 100 | 100 | 100 | 100 | 100 | 100 | 100 | 100 |
| 125 | 100 | 100 | 100 | 100 | 100 | 100 | 100 | 100 | 100 |
| 150 | 100 | 100 | 100 | 100 | 100 | 100 | 100 | 100 | 100 |

**Table S53.** Number and percentage of faithful mapping sites of 50 bp sequences mapped by Bowtie2.

| Chromosome | Number | | | Percentage (%) | | |
|---|---|---|---|---|---|---|
| | DNA | bMK | bRY | DNA | bMK | bRY |
| chr1 | 181,705,146 | 171,058,644 | 143,965,993 | 73 | 69 | 58 |
| chr2 | 197,852,765 | 186,442,919 | 159,025,035 | 82 | 77 | 66 |
| chr3 | 164,643,233 | 155,048,603 | 131,319,786 | 83 | 78 | 66 |
| chr4 | 156,629,212 | 147,887,392 | 124,921,862 | 82 | 78 | 66 |
| chr5 | 147,220,639 | 138,627,664 | 117,698,174 | 81 | 76 | 65 |
| chr6 | 140,714,566 | 132,698,360 | 112,525,624 | 82 | 78 | 66 |
| chr7 | 122,747,413 | 116,236,445 | 97,205,812 | 77 | 73 | 61 |
| chr8 | 120,117,573 | 112,875,381 | 95,744,009 | 83 | 78 | 66 |
| chr9 | 91,200,811 | 86,000,545 | 72,841,619 | 66 | 62 | 53 |
| chr10 | 107,341,251 | 101,145,618 | 85,417,526 | 80 | 76 | 64 |
| chr11 | 107,576,499 | 101,628,965 | 85,388,190 | 80 | 75 | 63 |
| chr12 | 107,826,095 | 101,251,528 | 84,813,031 | 81 | 76 | 64 |
| chr13 | 81,657,693 | 77,176,239 | 65,800,450 | 71 | 67 | 58 |
| chr14 | 72,906,277 | 68,634,357 | 57,838,337 | 68 | 64 | 54 |
| chr15 | 63,585,178 | 60,116,868 | 50,613,762 | 62 | 59 | 50 |
| chr16 | 60,346,626 | 56,488,631 | 47,201,859 | 67 | 63 | 52 |
| chr17 | 60,041,166 | 56,400,290 | 47,051,728 | 72 | 68 | 57 |
| chr18 | 63,990,476 | 60,285,947 | 51,882,503 | 80 | 75 | 65 |
| chr19 | 39,686,130 | 37,098,080 | 28,443,330 | 68 | 63 | 49 |
| chr20 | 50,701,550 | 47,635,805 | 40,258,316 | 79 | 74 | 62 |
| chr21 | 27,805,184 | 26,338,452 | 22,265,003 | 60 | 56 | 48 |
| chr22 | 26,506,604 | 25,033,193 | 20,844,310 | 52 | 49 | 41 |
| chrX | 112,316,720 | 106,074,709 | 82,558,058 | 72 | 68 | 53 |
| chrY | 8,180,587 | 8,473,337 | 5,331,627 | 14 | 15 | 9 |
| Sum | 2,313,304,912 | 2,180,665,789 | 1,830,957,106 | | | |

**Table S54.** Number and percentage of faithful mapping sites of 70bp sequences mapped by Bowtie2.

| Chromosome | Number | | | Percentage (%) | | |
|---|---|---|---|---|---|---|
| | DNA | bMK | bRY | DNA | bMK | bRY |
| chr1 | 200,914,700 | 198,392,685 | 173,716,632 | 81 | 80 | 70 |
| chr2 | 215,931,598 | 213,566,184 | 189,877,745 | 89 | 88 | 78 |
| chr3 | 180,512,867 | 178,045,677 | 157,541,578 | 91 | 90 | 79 |
| chr4 | 171,638,088 | 169,687,361 | 149,879,481 | 90 | 89 | 79 |
| chr5 | 161,208,233 | 159,102,213 | 141,071,656 | 89 | 88 | 78 |
| chr6 | 154,563,383 | 152,668,223 | 134,726,299 | 90 | 89 | 79 |
| chr7 | 136,234,711 | 135,735,813 | 117,631,676 | 85 | 85 | 74 |
| chr8 | 131,318,301 | 129,393,850 | 114,915,316 | 90 | 89 | 79 |
| chr9 | 100,288,854 | 99,410,964 | 87,520,898 | 72 | 72 | 63 |
| chr10 | 118,349,113 | 117,248,482 | 102,951,007 | 88 | 88 | 77 |
| chr11 | 118,488,387 | 117,551,335 | 103,229,117 | 88 | 87 | 76 |
| chr12 | 119,902,151 | 118,014,318 | 102,814,860 | 90 | 89 | 77 |
| chr13 | 88,706,555 | 87,981,451 | 78,464,198 | 78 | 77 | 69 |
| chr14 | 80,491,392 | 79,400,971 | 69,640,483 | 75 | 74 | 65 |
| chr15 | 70,248,096 | 69,888,853 | 61,074,458 | 69 | 69 | 60 |
| chr16 | 67,604,105 | 66,928,423 | 57,504,694 | 75 | 74 | 64 |
| chr17 | 68,295,547 | 67,596,368 | 57,344,709 | 82 | 81 | 69 |
| chr18 | 69,253,865 | 68,553,669 | 61,573,582 | 86 | 85 | 77 |
| chr19 | 48,242,382 | 47,219,308 | 37,018,457 | 82 | 81 | 63 |
| chr20 | 55,914,844 | 55,333,673 | 48,672,972 | 87 | 86 | 76 |
| chr21 | 30,500,761 | 30,544,229 | 26,792,693 | 65 | 65 | 57 |
| chr22 | 30,034,596 | 29,928,638 | 25,467,134 | 59 | 59 | 50 |
| chrX | 127,505,433 | 125,972,433 | 105,670,198 | 82 | 81 | 68 |
| chrY | 10,067,762 | 11,050,930 | 7,992,739 | 18 | 19 | 14 |
| Sum | 2,556,222,337 | 2,529,226,446 | 2,213,094,385 | | | |

**Table S55.** Number and percentage of faithful mapping sites of 100 bp sequences mapped by Bowtie2.

| Chromosome | Number | | | Percentage (%) | | |
|---|---|---|---|---|---|---|
| | DNA | bMK | bRY | DNA | bMK | bRY |
| chr1 | 210,027,652 | 210,301,316 | 189,658,069 | 84 | 84 | 76 |
| chr2 | 224,391,064 | 224,598,852 | 205,565,383 | 93 | 93 | 85 |
| chr3 | 188,090,261 | 187,462,272 | 171,218,967 | 95 | 95 | 86 |
| chr4 | 178,916,363 | 178,592,179 | 163,215,464 | 94 | 94 | 86 |
| chr5 | 167,759,539 | 167,446,977 | 153,213,173 | 92 | 92 | 84 |
| chr6 | 161,145,010 | 160,984,633 | 146,550,787 | 94 | 94 | 86 |
| chr7 | 142,992,769 | 144,557,327 | 129,388,693 | 90 | 91 | 81 |
| chr8 | 136,361,732 | 135,756,910 | 124,657,529 | 94 | 94 | 86 |
| chr9 | 104,732,528 | 105,403,606 | 95,289,933 | 76 | 76 | 69 |
| chr10 | 123,665,944 | 123,994,875 | 112,346,158 | 92 | 93 | 84 |
| chr11 | 124,041,187 | 124,469,815 | 112,737,068 | 92 | 92 | 83 |
| chr12 | 125,570,232 | 125,256,831 | 112,793,755 | 94 | 94 | 85 |
| chr13 | 92,001,075 | 92,065,151 | 84,885,244 | 80 | 81 | 74 |
| chr14 | 84,016,077 | 83,889,713 | 76,020,107 | 78 | 78 | 71 |
| chr15 | 73,617,319 | 74,250,175 | 66,736,582 | 72 | 73 | 65 |
| chr16 | 70,876,891 | 71,686,080 | 63,543,190 | 78 | 79 | 70 |
| chr17 | 72,486,798 | 73,310,272 | 63,695,548 | 87 | 88 | 77 |
| chr18 | 71,645,848 | 71,694,396 | 66,276,060 | 89 | 89 | 82 |
| chr19 | 52,423,833 | 52,618,005 | 43,394,139 | 89 | 90 | 74 |
| chr20 | 58,514,505 | 58,762,564 | 53,053,487 | 91 | 91 | 82 |
| chr21 | 31,856,480 | 32,452,300 | 29,293,706 | 68 | 69 | 63 |
| chr22 | 31,969,115 | 32,547,093 | 28,253,443 | 63 | 64 | 56 |
| chrX | 134,703,269 | 134,802,682 | 119,757,630 | 86 | 86 | 77 |
| chrY | 11,330,415 | 12,618,760 | 10,146,742 | 20 | 22 | 18 |
| Sum | 2,673,144,155 | 2,679,535,004 | 2,421,693,568 | | | |

**Table S56.** Number and percentage of faithful mapping sites of 50 bp sequences mapped by BWA-MEM.

| Chromosome | Number | | | Percentage (%) | | |
|---|---|---|---|---|---|---|
| | DNA | bMK | bRY | DNA | bMK | bRY |
| chr1 | 120,087,173 | - | - | 48 | 0 | 0 |
| chr2 | 132,185,944 | - | - | 55 | 0 | 0 |
| chr3 | 108,521,380 | - | - | 55 | 0 | 0 |
| chr4 | 101,370,280 | - | - | 53 | 0 | 0 |
| chr5 | 97,138,458 | - | - | 54 | 0 | 0 |
| chr6 | 92,843,685 | - | - | 54 | 0 | 0 |
| chr7 | 79,935,201 | - | - | 50 | 0 | 0 |
| chr8 | 79,356,790 | - | - | 55 | 0 | 0 |
| chr9 | 60,775,541 | - | - | 44 | 0 | 0 |
| chr10 | 71,296,346 | - | - | 53 | 0 | 0 |
| chr11 | 71,365,667 | - | - | 53 | 0 | 0 |
| chr12 | 69,997,164 | - | - | 53 | 0 | 0 |
| chr13 | 53,747,343 | - | - | 47 | 0 | 0 |
| chr14 | 47,963,542 | - | - | 45 | 0 | 0 |
| chr15 | 42,373,083 | - | - | 42 | 0 | 0 |
| chr16 | 39,888,607 | - | - | 44 | 0 | 0 |
| chr17 | 39,807,626 | - | - | 48 | 0 | 0 |
| chr18 | 43,199,905 | - | - | 54 | 0 | 0 |
| chr19 | 23,228,491 | - | - | 40 | 0 | 0 |
| chr20 | 34,203,432 | - | - | 53 | 0 | 0 |
| chr21 | 18,368,778 | - | - | 39 | 0 | 0 |
| chr22 | 17,821,310 | - | - | 35 | 0 | 0 |
| chrX | 65,910,905 | - | - | 42 | 0 | 0 |
| chrY | 3,578,376 | - | - | 6 | 0 | 0 |
| Sum | 1,514,967,552 | - | - | | | |

**Table S57.** Number and percentage of faithful mapping sites of 70 bp sequences mapped by BWA-MEM.

| Chromosome | Number | | | Percentage (%) | | |
|---|---|---|---|---|---|---|
| | DNA | bMK | bRY | DNA | bMK | bRY |
| chr1 | 177,926,804 | 150,978,437 | 115,969,538 | 71 | 61 | 47 |
| chr2 | 194,636,536 | 165,884,400 | 130,596,093 | 80 | 68 | 54 |
| chr3 | 161,965,868 | 137,592,204 | 107,389,267 | 82 | 69 | 54 |
| chr4 | 154,288,718 | 131,026,634 | 102,753,056 | 81 | 69 | 54 |
| chr5 | 144,934,506 | 123,074,298 | 96,484,054 | 80 | 68 | 53 |
| chr6 | 138,317,158 | 117,576,001 | 92,290,236 | 81 | 69 | 54 |
| chr7 | 120,074,044 | 101,655,952 | 78,986,993 | 75 | 64 | 50 |
| chr8 | 118,235,998 | 100,201,778 | 78,219,518 | 81 | 69 | 54 |
| chr9 | 89,435,293 | 76,037,765 | 59,197,057 | 65 | 55 | 43 |
| chr10 | 105,104,858 | 89,062,991 | 69,414,300 | 79 | 67 | 52 |
| chr11 | 105,505,949 | 89,671,804 | 68,734,819 | 78 | 66 | 51 |
| chr12 | 105,531,912 | 88,950,580 | 68,311,223 | 79 | 67 | 51 |
| chr13 | 80,442,530 | 68,676,465 | 54,457,940 | 70 | 60 | 48 |
| chr14 | 71,510,947 | 60,520,726 | 47,042,477 | 67 | 57 | 44 |
| chr15 | 62,146,092 | 52,866,330 | 40,800,110 | 61 | 52 | 40 |
| chr16 | 58,633,435 | 49,170,372 | 37,809,880 | 65 | 54 | 42 |
| chr17 | 57,985,305 | 48,771,644 | 37,402,636 | 70 | 59 | 45 |
| chr18 | 63,072,162 | 53,943,434 | 42,947,252 | 78 | 67 | 53 |
| chr19 | 37,714,204 | 30,068,219 | 20,951,263 | 64 | 51 | 36 |
| chr20 | 49,581,711 | 42,044,230 | 32,315,050 | 77 | 65 | 50 |
| chr21 | 27,274,356 | 23,135,544 | 18,299,625 | 58 | 50 | 39 |
| chr22 | 25,610,992 | 21,646,921 | 16,470,623 | 50 | 43 | 32 |
| chrX | 110,262,975 | 89,841,524 | 63,108,608 | 71 | 58 | 40 |
| chrY | 7,756,842 | 6,128,916 | 3,147,315 | 14 | 11 | 5 |
| Sum | 2,267,953,777 | 1,918,532,803 | 1,483,099,512 | | | |

**Table S58.** Number and percentage of faithful mapping sites of 100 bp sequences mapped by BWA-MEM.

| Chromosome | Number | | | Percentage (%) | | |
|---|---|---|---|---|---|---|
| | DNA | bMK | bRY | DNA | bMK | bRY |
| chr1 | 199,478,772 | 192,065,909 | 165,804,584 | 80 | 77 | 67 |
| chr2 | 214,632,296 | 207,647,865 | 182,181,521 | 89 | 86 | 75 |
| chr3 | 179,424,962 | 173,108,623 | 150,814,805 | 90 | 87 | 76 |
| chr4 | 170,638,850 | 164,722,179 | 143,081,558 | 90 | 87 | 75 |
| chr5 | 160,280,452 | 154,669,223 | 135,071,154 | 88 | 85 | 74 |
| chr6 | 153,624,448 | 148,179,962 | 128,763,964 | 90 | 87 | 75 |
| chr7 | 135,304,639 | 130,270,817 | 111,200,797 | 85 | 82 | 70 |
| chr8 | 130,624,079 | 126,135,087 | 110,148,650 | 90 | 87 | 76 |
| chr9 | 99,555,833 | 96,063,570 | 83,583,338 | 72 | 69 | 60 |
| chr10 | 117,645,534 | 113,371,328 | 97,963,290 | 88 | 85 | 73 |
| chr11 | 117,675,576 | 113,468,312 | 98,176,520 | 87 | 84 | 73 |
| chr12 | 119,073,947 | 114,217,830 | 97,893,352 | 89 | 86 | 73 |
| chr13 | 88,308,079 | 85,643,202 | 75,082,765 | 77 | 75 | 66 |
| chr14 | 79,972,639 | 77,044,236 | 66,487,734 | 75 | 72 | 62 |
| chr15 | 69,731,637 | 67,278,865 | 58,077,844 | 68 | 66 | 57 |
| chr16 | 67,052,038 | 64,248,640 | 54,555,727 | 74 | 71 | 60 |
| chr17 | 67,556,543 | 64,390,101 | 54,071,336 | 81 | 77 | 65 |
| chr18 | 68,886,426 | 66,891,859 | 59,290,352 | 86 | 83 | 74 |
| chr19 | 47,696,498 | 44,189,076 | 33,472,897 | 81 | 75 | 57 |
| chr20 | 55,485,463 | 53,416,046 | 46,524,861 | 86 | 83 | 72 |
| chr21 | 30,282,530 | 29,307,229 | 25,440,539 | 65 | 63 | 54 |
| chr22 | 29,689,556 | 28,371,461 | 23,901,822 | 58 | 56 | 47 |
| chrX | 126,741,328 | 120,783,672 | 97,652,094 | 81 | 77 | 63 |
| chrY | 10,014,742 | 9,412,374 | 5,912,176 | 17 | 16 | 10 |
| Sum | 2,539,382,925 | 2,444,905,080 | 2,105,154,566 | | | |

**Table S59.** Basic metrics of sequenced samples. To compare with BitSeq, sequencing data from ion torrent or illumina sequencer were down-sampled with read number near that of the same sample sequenced by BitSeq. CAP: Community-Acquired Pneumoniae.

| Sample | Read Number | | |
|---|---|---|---|
| | BitSeq | Ion Torrent | Illumina |
| Patient 1: NIPT Reference | 2594296 | 2594602 | - |
| Patient 2: NIPT Reference | 4602556 | 4602055 | - |
| Patient 3: NIPT Reference | 3016336 | 3014533 | - |
| Patient 4: NIPT Reference | 3164233 | 3162685 | - |
| Patient 5: NIPT Reference | 2328559 | 2328029 | - |
| Patient 6: NIPT Reference | 2111313 | 2131423 | - |
| Patient 7: NIPT Reference | 2713347 | 2713927 | - |
| Patient 8: NIPT Reference | 3853911 | 3851839 | - |
| Patient 9: NIPT Reference | 2691415 | 2691735 | - |
| Patient 10: NIPT Reference | 3626592 | 3623884 | - |
| Patient 11: NIPT Reference | 2875558 | 2873002 | - |
| Patient 12: NIPT Reference | 3068976 | 3065791 | - |
| Patient 13: NIPT Reference | 2226745 | 2225478 | - |
| Patient 14: NIPT Reference | 4662377 | 4659457 | - |
| Patient 15: NIPT Reference | 2670094 | 2669166 | - |
| Patient 16: NIPT Reference | 2598998 | 2597138 | - |
| Patient 17: NIPT Reference | 4247767 | 4244596 | - |
| Patient 18: NIPT Reference | 2968056 | 2966464 | - |
| Patient 19: NIPT Reference | 1625921 | 1625613 | - |
| Patient 20: NIPT Reference | 1802098 | 1801208 | - |
| Patient 21: NIPT Reference | 1953112 | 1951716 | - |
| Patient 22: NIPT Reference | 3150652 | 3150114 | - |
| Patient 23: NIPT Reference | 1815489 | 1820818 | - |
| Patient 24: NIPT Reference | 2227212 | 2226578 | - |
| Patient 25: NIPT Negative | 1791432 | 1792455 | - |
| Patient 26: NIPT Negative | 2469324 | 2471413 | - |
| Patient 27: NIPT Negative | 2934496 | 2933950 | - |
| Patient 28: NIPT Negative | 1590043 | 1590173 | - |
| Patient 29: 10% T21 mock | 3854635 | 3854467 | - |
| Patient 30: 5% T21 mock | 2298243 | 2298456 | - |
| Patient 31: 3.5% T21 mock | 3062991 | 3061723 | - |
| Patient 32: 2.5% T21 mock | 3002574 | 3000595 | - |
| Patient 33: 10% T13 mock | 1581030 | 1599279 | - |
| Patient 34: 5% T13 mock | 2987538 | 2985929 | - |
| Patient 35: 3.5% T13 mock | 3946439 | 3946603 | - |
| Patient 36: 2.5% T13 mock | 3734486 | 3733519 | - |
| Patient 37: 10% T18 mock | 2268176 | 2268157 | - |
| Patient 38: 5% T18 mock | 2363669 | 2362574 | - |

| | | | |
|---|---|---|---|
| Patient 39: 3.5% T18 mock | 2851004 | 2851756 | - |
| Patient 40: 2.5% T18 mock | 3680951 | 3683618 | - |
| Patient 41: Single-blinded NIPT | 4231466 | 4231214 | - |
| Patient 42: Single-blinded NIPT | 1563332 | 1563114 | - |
| Patient 43: Single-blinded NIPT | 1730517 | 1738079 | - |
| Patient 44: Single-blinded NIPT | 2154953 | 2162132 | - |
| Patient 45: Single-blinded NIPT | 3019251 | 3036386 | - |
| Patient 46: Single-blinded NIPT | 2030995 | 2031259 | - |
| Patient 47: Single-blinded NIPT | 2274493 | 2273373 | - |
| Patient 48: Single-blinded NIPT | 2435671 | 2439792 | - |
| Patient 49: Single-blinded NIPT | 1798844 | 1804056 | - |
| Patient 50: Single-blinded NIPT | 3361976 | 3373498 | - |
| Patient 51: Down Syndrome | 3559677 | 3559677 | - |
| Patient 52: normal male | 3086001 | 3086001 | - |
| Patient 53: DiGeorge Syndrome | 3003276 | 3003276 | 3003276 |
| Patient 54: Development Delay | 2240883 | 2240883 | 2240883 |
| Patient 55: Pediatric patient, anal swab | 7985374 | - | 8628545 |
| Patient 55: Pediatric patient, throat swab | 8000000 | - | 7700905 |
| Patient 56: CAP 1 | 1000000 | - | 1000000 |
| Patient 57: CAP 2 | 1000000 | - | 1000000 |
| Patient 58: CAP 3 | 1000000 | - | 1000000 |
| Patient 59: CAP 4 | 1000000 | - | 1000000 |
| Patient 60: CAP 5 | 1000000 | - | 1000000 |
| Patient 61: CAP 6 | 1000000 | - | 1000000 |
| Patient 62: CAP 7 | 1000000 | - | 1000000 |
| Patient 63: CAP 8 | 1000000 | - | 1000000 |
| Patient 64: CAP 9 | 1000000 | - | 1000000 |
| Patient 65: CAP 10 | 1000000 | - | 1000000 |
| Patient 66: CAP 11 | 1000000 | - | 1000000 |
| Patient 67: CAP 12 | 1000000 | - | 1000000 |
| Patient 68: CAP 13 | 1000000 | - | 1000000 |
| Patient 69: CAP 14 | 1000000 | - | 1000000 |
| Mouse embryonic stem cell RNA | 539155 | 560000 | 610000 |
| Mouse embryonic fibroblast RNA | 783455 | 810000 | 890000 |

### Captions for Movies

**Movie S1.** The fluorescence imaging process. After each cycle of reaction, the flow-cell was cooled down to room temperature and then the imaging process started. The fluorescence images were taken in tiles, and every cycle we took 171 (19 by 9) images to cover the whole flow-cell.

**Movie S2.** A small field of view demonstrating the cyclic reaction and fluorogenic signals. The total cycle number is 69 for this experiment, and the images are drifted due to the motion errors and thermal cycles. Through image registration every microwell is indexed, and two of which are labeled by red and green circles in the video. The signal intensity varies in each well through the cyclic sequencing reaction, representing the different length of elongation during each reaction cycle. Such intensity series can be quantified and then go through imaging processing and other correction procedure to generate the bit sequences.
